# Forces directing the systemic correlations of cell migration

**DOI:** 10.1101/2024.04.22.590476

**Authors:** Ildefonso M. De la Fuente, Jose Carrasco-Pujante, Borja Camino-Pontes, Maria Fedetz, Carlos Bringas, Alberto Pérez-Samartín, Gorka Pérez-Yarza, José I. López, Iker Malaina, Jesus M Cortes

## Abstract

Directional motility is an essential property of cells. Despite its enormous relevance in many fundamental physiological and pathological processes, how cells control their locomotion movements remains an unresolved question. Here we have addressed the systemic processes driving the directed locomotion of cells. Specifically, we have performed an exhaustive study analyzing the trajectories of 700 individual cells belonging to three different species (*Amoeba proteus*, *Metamoeba leningradensis* and *Amoeba borokensis*) in four different scenarios: in absence of stimuli, under an electric field (galvanotaxis), in a chemotactic gradient (chemotaxis), and under simultaneous galvanotactic and chemotactic stimuli. All movements were analyzed using advanced quantitative tools. The results show that the trajectories are mainly characterized by coherent integrative responses that operate at the global cellular scale. These systemic migratory movements depend on the cooperative non-linear interaction of most, if not all, molecular components of cells.

**Significance:** Cellular migration is a cornerstone issue in many human physiological and pathological processes. For years, the scientific attention has been focused on the individualized study of the diverse molecular parts involved in directional motility; however, locomotion movements have never been regarded as a systemic process that operates at a global cellular scale. In our quantitative experimental analysis essential ***systemic*** properties underlying locomotion movements were detected. Such emergent systemic properties are not found specifically in any of the molecular parts, partial mechanisms, or individual processes of the cell. Cellular displacements seem to be regulated by integrative processes operating at ***systemic*** level.

## Introduction

Self-locomotion is one of the most important complex behaviors of cells endowed with migratory responses. In the permanent struggle for survival free cells move efficiently to find food following adequate direction and speed, avoiding predators and adverse conditions. Cell motility is crucial for life in Metazoan organisms and fundamental to establish the appropriate organization of all multicellular organisms, playing a central role in a plethora of essential biological phenomena such as embryogenesis, morphogenesis, organogenesis, neural development, adult tissue remodeling, wound healing, immune responses, angiogenesis, tissue regeneration and repair, cell differentiation, etc. (1). Moreover, the deregulation of cell movements in humans is involved in many pathological processes such as chronic inflammatory diseases, atherosclerosis, vascular diseases, osteoporosis, congenital brain pathologies, hearing disorders, tumorigenesis and metastasis, among others (2–5).

Although cell locomotion was already observed in 1675 by van Leeuwenhoek in his respected pioneer microscopic studies (6), researchers and scholars have not yet come to unveil how cells move efficiently through diverse environments and migrate in the presence of complex cues.

Given its importance, a great attention has been focused on the study of the diverse molecular parts involved in directional motility. A relevant number of these experimental studies have unequivocally shown that locomotion movements are complex processes that involve practically all cellular components. So, directed movements are primarily driven by the cytoskeleton (the essential part of the locomotion system) which is a sophisticated dynamic structure formed by three main molecular components: actin microfilaments, microtubules, and intermediate filaments, all of them interacting in complex dynamic networks (7).

In particular, the activity of actin cytoskeleton networks is largely dependent on a wide variety of regulatory molecules such as small GTPases (8), integrins (9), and many post-translational modifications such as phosphorylation, acetylation, arginylation, oxidation, and others (10). In addition, the dynamic turnover of the actin filament networks is essential to regulate cell migration (11). Cytoskeleton networks are coupled with other complex regulated systems such as membrane surface receptors and signal transduction pathways which also participate in the control of locomotion movements (12). Energy is another essential element in cell motility; when cells move the cytoskeleton transforms chemical energy into mechanical forces (dynein cytoskeletal motor proteins) entailing considerable bioenergetic demands; for such a purpose the mitochondrial activity and the adenylate energy system are important regulators of directionality motion (13). Cell membrane activities are also necessary to implement an adequate migration (14, 15).

Recent studies have revealed the importance of autophagy (an intracellular process that controls protein and organelle degradation and recycling) in the control of locomotion (16). The turnover of focal adhesions also regulates cell spreading and migration (17). Calcium ions (Ca^2+^), which impact globally on almost every aspect of cellular life, play an important role in the control of directed movements (18). In this sense, the endoplasmic reticulum, a multifunctional signaling organelle which controls a wide range of cellular processes such as the entry and release of calcium ions, also participate in the regulation of cell locomotion (19, 20). Cell polarity is required for an adequate directionality motion and there are a lot of molecular processes that have been implicated in the intrinsic polarity status of cells; in this regard, the centrosomes positioning serves as a steering device for the directional movement (21, 22) and dynein together with other molecules regulates centrosomal orientation to establish and maintain cell polarity (23).

The Golgi apparatus (another important molecular processing center for modified proteins received from the endoplasmic reticulum) allows the remodeling of intracellular traffic processes towards the direction of movement; therefore, signals from the Golgi matrix play an important role in cell motility (24). The nucleus is very important for developing appropriate mechanical responses during cell migration, in fact this organelle behaves as a central mechanosensory structure, and its physical properties strongly connected to the cytoskeleton guarantee a proper cell migration (25, 26). Recently, it has been described that structural chromatin organization also has a key role in the cellular migration process (27). In addition, many molecules and corresponding processes are involved in directional movement of cells as, for instance, focal adhesion proteins (talin, paxillin, vinculin and others) (12), SCAR/WAVE proteins (28), actin-binding proteins (ABPs) (29), p21-activated kinases (PAKs, a family of serine/threonine kinases) (30), TORC2/PKB pathway (31), mitogen-activated protein kinases (MAPKs) (32), Arp2/3 complexes (33), WASP family proteins (34), Nck family of adaptor proteins (35), etc.

All this evidence suggests that cell migration is not a mere metabolic-molecular process which can be regulated by any of its individually considered components. Most, if not all, fundamental cellular physiological processes appear to be involved in cellular locomotion, which is indicative of the emergence of a global functional phenomenon in the cell. However, confirming the systemic nature of cellular locomotion represents a scientific challenge of great difficulty. Such verification requires multidisciplinary approaches that combine complex experimental studies with advanced quantitative methods.

Here, we have addressed the key question: Is cell migration a highly coordinated and integrated emergent process at the global cellular level? To answer this question, we have designed a large quantitative study to analyze the systemic trajectories of 700 individual cells belonging to three different species: *Amoeba proteus*, *Metamoeba leningradensis* and *Amoeba borokensis*. Such analysis has been performed under four different scenarios: in absence of stimuli, under chemotactic gradient (we have used a nFMLP peptide, which indicates to the amoebae the possible presence of food in their immediate environment), in an electric field (the electric membrane potential of cells enables predators like amoebas the detection of preys), and under complex external conditions such as simultaneous galvanotactic and chemotactic gradient stimuli.

To understand the forces driving the locomotion movement of the cell, all trajectories were analyzed using computational methods and advanced non-linear physical-mathematical tools rooted in Statistical Physics (Statistical Mechanics). These quantitative studies focused on some essential characteristics of the systemic dynamics underlying locomotion movements. The results indicate that a very complex dynamic structure emerges in the migratory movements of all the cells analyzed. Such structure is characterized by highly organized move-step sequences with very low entropy and high information, non-trivial long-range interdependence in the move-steps, strong anomalous super-diffusion dynamics, long-term memory effects with trend-reinforcing behavior, and efficient movements to explore the extracellular medium.

This outstanding cellular dynamic structure is a consequence of the emergent systemic dynamics occurring in the cell. The locomotion movements seem to depend on a complex integrated self-organized system carefully regulated at global level, arising from the cooperative non-linear interaction of most, if not all, cellular components. Such emergent systemic properties are not found specifically in any of the molecular parts, partial mechanisms, or individual processes of the cell.

The findings presented here open a new perspective to improve the conceptual framework of the cell, and to develop researches on the pathological deregulation of migratory movements combining systemic analysis with molecular studies.

## Results

The migratory trajectories of 700 individual cells belonging to the three species, *A. proteus, M. leningradensis* and *A. borokensis*, were recorded in four different scenarios: in absence of stimuli, under chemotactic gradient, in an electric field, and under simultaneous galvanotactic and chemotactic stimuli. Amoebae show robust movement in response to an electric field in a range between 300 mV/mm and 600 mV/mm (galvanotaxis). Under such conditions, practically all amoebae migrate towards the cathode (36). Likewise, these cells also exhibit chemotactic movements. More specifically, the peptide nFMLP (N-Formylmethionyl-leucyl-phenylalanine) secreted by bacteria may indicate that food might be in the near environment, provoking a strong chemotactic response (37).

All our experiments were performed in a specific set-up consisting of two standard electrophoresis blocks (17.5 cm long), two agar bridges, a power supply, and in the middle of the experimental platform, a structure of standard glass slide and covers where the cells were located (see *SI Appendix*, Fig. S1 and *Materials and Methods*). One electrophoresis block was directly plugged into a normal power supply and the other was connected to the first one through two agar bridges, thus preventing the direct contact of the anode and cathode with the medium (Chalkley’s simplified medium (36)) where the cells were placed. Specifically, the amoebae were arranged in the center of the structure of standard glass slide and covers (experimental chamber) and their migratory displacements were monitored. The glass experimental structure enabled the generation of a laminar flux allowing the electric current to pass through, on one hand, and generating an nFMLP peptide gradient, on the other (see *Materials and Methods* and *SI Appendix*, Figs. S1 and S2).

Prior to each experiment, all cells were starved for 24h. The individual migratory movements of each cell were recorded over periods of 30 minutes using a digital camera attached to a stereo microscope. The experiments on flat two-dimensional surfaces were always made with small groups of cells (no more than 9 cells per replication). The following basic experimental information data (BEID) is provided for each scenario: "Nr" the number of cells per replication, "Er" the number of experimental replications, and "N" is the total number of cells. Finally, the recorded trajectories were analyzed in the form of time series using advanced non-linear dynamic tools.

### 1. Cellular migratory movements without external stimulus

First, we recorded the locomotion trajectories of 153 individual cells belonging to the three species considered in a medium without any external influence (BEID: *A. proteus*: N=50, Er=7, Nr=7-8; *M. leningradensis*: N=51, Er=7, Nr=5-8; *A. borokensis*: N=52, Er=7, Nr=6-8). In Fig. 1*A*, a representative example of these amoebae migratory movements in absence of stimuli is depicted (for clarity only 60 cells were randomly taken from the total). It can be observed that after 30 minutes, cells have explored practically all the directions of the experimentation chamber. To quantitatively analyze cell directionality, we calculated the displacement cosine for each trajectory, 153 cells in total (Fig. 1*A*’). Values close to -1 indicate a preference towards the left, while values close to 1 suggest a preference towards the right. Our analysis showed that values ranged between -1 and 1, with a median/IQR of 0.052/1.397. Median/IQR values for each species were 0.417/1.244 (*A. proteus*), - 0.251/1.207 (*M. leningradensis*) and 0.126/1.232 (*A. borokensis*). These results indicate that in absence of stimuli cells moved randomly without any defined guidance.

**Fig. 1.**
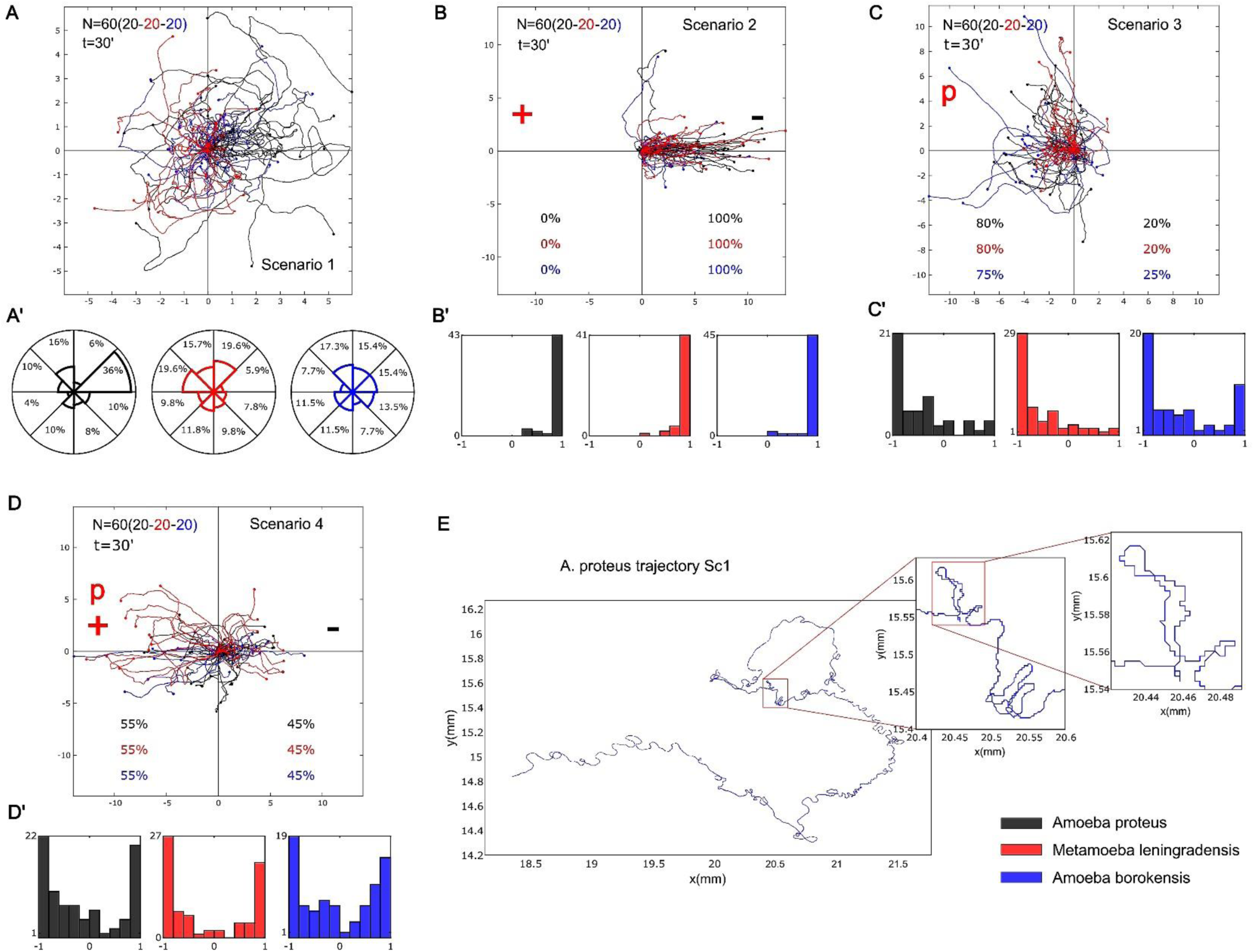
Representative migration trajectories of the three species under four experimental scenarios. *(A-D)* Migration behavior of the three species (in black *Amoeba proteus*, in red *Metamoeba leningradensis*, and in blue *Amoeba borokensis*) in the four experimental scenarios (“Scenario 1” absence of stimuli, “Scenario 2” presence of an electric field, “Scenario 3” presence of a chemotactic peptide gradient and “Scenario 4” simultaneous galvanotactic and chemotactic stimuli). *(A’)* the percentage of cells moving in any specific sector of the experimental chamber is represented as a polar histogram divided into 8 areas of π/4 angle amplitude each. *(B’–D’)* histograms of the displacement cosines from panels *(B–D)*, respectively. *(E)* Digitized cell trajectory with two inserts highlighting displacements regions of interest. “N” is the total number of cells; “t”, is the time of galvanotaxis or chemotaxis; “p”, is the chemotactic peptide (nFMLP); “+”, represents the anode; “−”, represents the cathode; “Sc1” Scenario One (absence of stimuli). Both the x and y-axis show the distance in mm, and the initial location of each cell has been placed at the center of the diagram.

### 2. Cell migration under galvanotaxis conditions

The migratory trajectories of 147 cells belonging to the three species were recorded under an external controlled direct-current electric field of about 300-600 mV/mm. In Fig. 1*B*, a representative example of the migratory movements of 60 cells is depicted. They show an unequivocal systemic response consisting of the migration to the cathode which has been placed on the right side of the set-up. The overall median/IQR value of the displacement cosines of all 147 cells (Fig. 1*B*’) was 0.989/0.0732. This finding confirmed that a fundamental behavior characterized by an unequivocal directionality towards the cathode had emerged under these galvanotactic conditions. The median/IQR values for each species were 0.993/0.023 (*A. proteus*), 0.978/0.098 (*M. leningradensis*) and 0.985/0.083 (*A. borokensis*). We compared the distributions of the values for the displacement cosines under galvanotaxis with the values obtained in the experiment without stimuli using the Wilcoxon rank-sum test. The results indicated that both behaviors were significantly different for the three species and that the galvanotactic cellular behavior is highly unlikely to be obtained by chance (p-values: 10^-9^, 10^-15^, and 10^-12^; Z: -5.854, -7.789, and -6.845 for *A. proteus*, *M. leningradensis*, and *A. borokensis*, respectively). BEID: *A. proteus*: N=49, Er=7, Nr=6-8; *M. leningradensis*: N=48, Er=7, Nr=6-8; *A. borokensis*: N=50, Er=8, Nr=3-9.

### 3. Cell locomotion under chemotaxis conditions

The migratory behavior of 166 cells belonging to the three species considered was recorded under conditions of chemotactic gradient. All the amoebae were exposed for 30 minutes to an nFMLP peptide gradient which had been placed on the left side of the set-up. In Fig. 1*C* a representative example of these migratory trajectories with 60 cells is depicted. Under these conditions 78.31% of all studied amoebae showed locomotion movements towards the attractant peptide.

The displacement angle cosines of the 166 individual trajectories ranged from -1 to 1, with a median/IQR value of -0.672/0.874. Median/IQR values for each species were -0.652/0.745 (*A. proteus*), -0.774/0.676 (*M. leningradensis*) and -0.513/1.17 (*A. borokensis*), indicating that they exhibited a single fundamental behavior of movement towards the peptide (Fig. 1*C*’). The Wilcoxon rank-sum test showed significant differences between the cosine values obtained with and without chemotactic stimulus (p-values: 10^-7^, 0.003, and 0.016; Z: 5.0841, 2.9379, and 2.403 for *A. proteus*, *M. leningradensis*, and *A. borokensis*, respectively), and between the cosine values with chemotactic gradient and with the presence of an electric field (p-values: 10^-15^, 10^-17^, and 10^-14^; Z: 7.998, 8.554, and 7.677 for *A. proteus*, *M. leningradensis*, and *A. borokensis*, respectively). This confirmed that the systemic locomotion behavior under the chemotactic gradient was completely different from both the absence of stimuli and the presence of an electric field. BEID: *A. proteus*: N=51, Er=8, Nr=5-7; *M. leningradensis*: N=60, Er=9, Nr=5-8; *A. borokensis*: N=55, Er=10, Nr=5-7.

### 4. Cellular displacement under simultaneous galvanotactic and chemotactic stimuli

Once the locomotion movements of the cells were recorded under the three previous independent experimental scenarios (without stimuli, under galvanotaxis and under chemotaxis), we studied the trajectories of 234 cells under simultaneous galvanotactic and chemotactic stimuli. For such a purpose, the nFMLP peptide was arranged on the left of the set-up (in the anode area) and the cathode was placed on the right. In Fig. 1*D*, a representative example of these locomotion movements (60 cells in total) is depicted. Under these complex external conditions, the results showed that 42% of the amoebae migrated towards the cathode while the remaining 58% moved towards the peptide (anode).

The displacement cosines of the 234 cells had an overall median/IQR value of -0.286/1.659. More specifically, the values for each species were (-0.315/1.591, median/IQR) for *A. proteus*, (- 0.542/1.811, median/IQR) for *M. leningradensis*, and (-0.137/1.453, median/IQR) for *A. borokensis*. This analysis quantitatively verified that two main cellular migratory behaviors had emerged in the experiment, one towards the anode and another towards the cathode (Fig. 1*D*’). The statistical analysis (Wilcoxon rank-sum test) confirmed the presence of these two different behaviors for *A. proteus* (p-value=10^-14^; Z=7.672), *M. leningrandensis* (p-value=10^-13^; Z=7.226) and *A. borokensis* (p-value=10^-14^; Z=7.5549). BEID: *Amoeba proteus*: N=83, Er=12, Nr=6-8; *Metamoeba leningradensis*: N=73, Er=11, Nr=5-8; *Amoeba borokensis*: N=78, Er=12, Nr=4-8.

### 5. Long-range interdependence in the move-steps of cellular migratory displacements

An essential characteristic of systemic behavior in complex systems is the presence of dynamics with strong long-range correlations (38). One of the most recognized tools to analyze the presence of these correlations in time series (migratory trajectories here) is the “root mean square fluctuation” (“rmsf” analysis), a classical method in Statistical Mechanics based on the ideas raised by Gibbs (39) and Einstein (40).

Long-range interdependence can be detected by a power-law relation such that *F*(*l*)∼*l*^*α*^. Where *l* is the number of steps. For uncorrelated data, the fluctuation exponent *α* is about 0.5, whereas *α* > 0.5 or *α* < 0.5 indicate respectively the presence of positive or negative long-range correlations (*Materials and Methods*). In Fig. 2*A*, an illustrative “rmsf” analysis for the locomotion movements of three representative cells belonging to each species considered under simultaneous chemotactic and galvanotactic stimuli (*A. proteus* and *M. leningradensis*) and under chemotactic conditions (*A. borokensis*) is depicted.

**Fig. 2.**
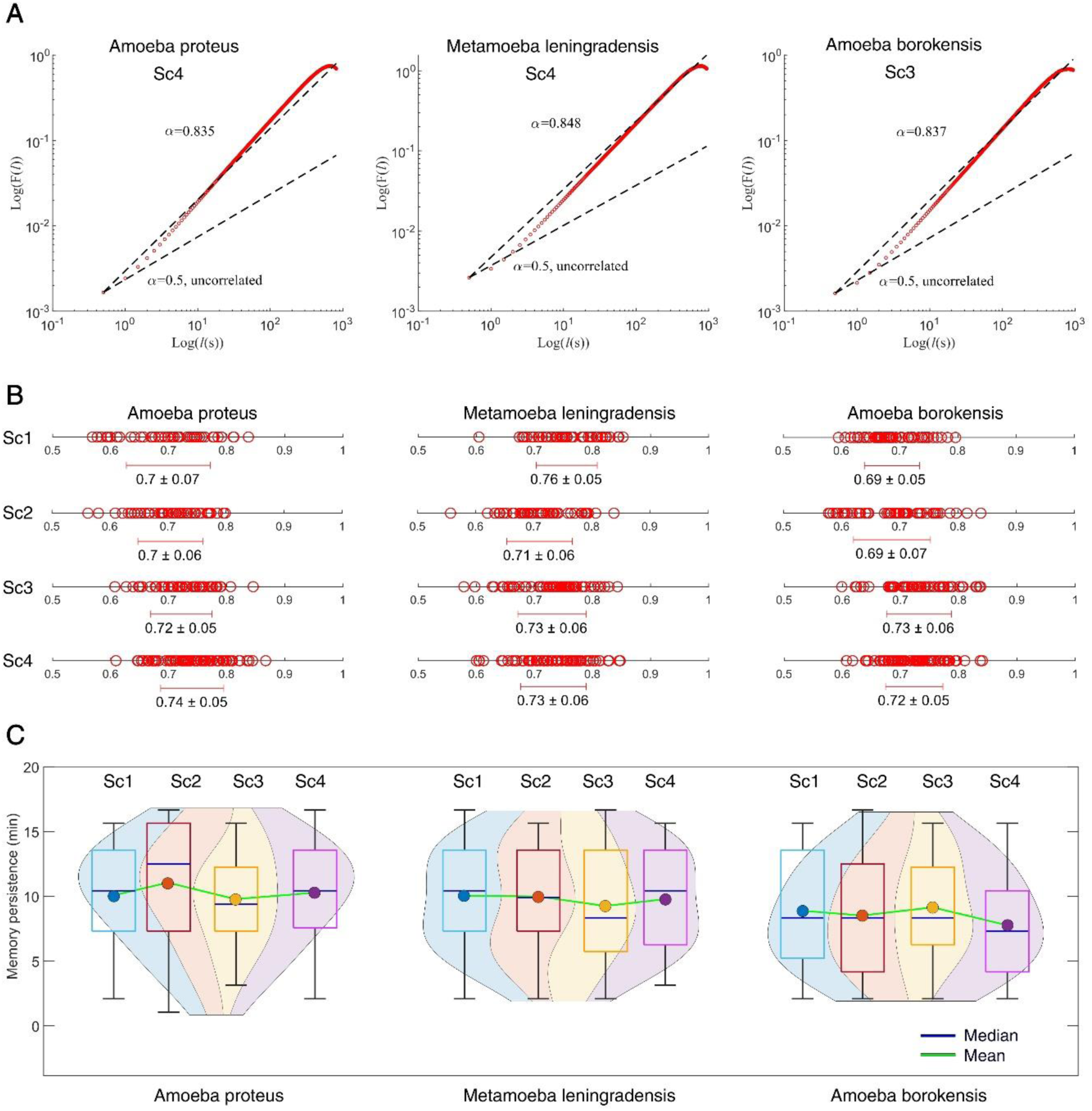
Long-range interdependence in the move-steps of cellular migratory displacements. *(A)* Log-log plot of RMSF *F* versus *l* step for a representative cell of each species. The slope was α=0.835 for *A. proteus*, α=0.848 for *M. leningradensis* and α=0.837 for *A. borokensis*, indicating the presence of strong long-range interdependence in the move-steps of all of them. *(B)* Diagram representing the values (and the overall average ± SD) of all the scaling exponents α from cells belonging to each species (*A. proteus*, *M. leningradensis* and *A. borokensis*) under each experimental scenario (Sc1-Sc4). *(C)* Violin plots showing the estimated distribution, median and average memory persistence values from cellular trajectories. “Sc1” Scenario One, absence of stimuli; “Sc2” Scenario Two, presence of an electric field; “Sc3” Scenario Three, presence of a chemotactic peptide gradient and “Sc4” Scenario Four, simultaneous galvanotactic and chemotactic stimuli.

The results of “rmsf” analysis of the 700 experimental cell trajectories are shown in Fig. 2*B* (for more details see *SI Appendix*, Table S1). All the migratory trajectories exhibit non-trivial correlations in their cellular move-step migratory fluctuations. Specifically, we found that the scaling exponent α of the “rmsf” had a median/IQR value of 0.723/0.08 for *A. proteus*, 0.734/0.081 for *M. leningradensis*, and 0.707/0.076 for *A. borokensis*. The values of the “rmsf” analysis of the total experimental migratory trajectories analyzed ranged from 0.557 to 0.867, with a median/IQR of 0.723/0.08, whereas the values of the scaling exponent α of all shuffled trajectories ranged from 0.351 to 0.644, with a median/IQR value of 0.468/0.069 (see *SI Appendix*, Table S2 for more details). Moreover, a Wilcoxon test comparing measured exponents to those from shuffled trajectories revealed highly significant long-range correlations in our data (p-value≅0, Z= -32.312), indicating the improbability of chance occurrence.

We also calculated the time duration of the correlations regime and found that all cells exhibited long-range coordination over periods ranging from 1.042 to 16.667 minutes with a median/IQR value of 9.375/7.292 minutes. These findings indicate strong dependences of past movements lasting approximately 1125/875 (median/IQR) move-steps (Fig. 2*C* and *SI Appendix*, Table S3). *A. proteus* cells exhibited long-range correlations up to a median/IQR duration of 10.417/6.25 minutes, *M. leningradensis* cells showed 9.375/7.292 minutes and *A. borokensis* cells 8.333/6.25 minutes, thus highlighting the influence of previous trajectory values on each cellular move-step. These results show the presence of non-trivial long-range correlations in all migration trajectories.

### 6. Strong anomalous migratory dynamics in cellular locomotion

Another characteristic of the migratory movement of cells is their strong anomalous dynamics. This property is directly related to anomalous super-diffusion, a complex process with a high non-linear relationship to time which also corresponds to efficient systemic directional trajectories (41, 42).

One of the best methods to determine such dynamic property is the Mean Square Displacement (MSD), a method proposed by Einstein (43) and later by Smoluchowski (44). This Statistical Mechanics tool allows to quantify the amount of space explored by the amoebae during their locomotion. According to this procedure (see *Materials and Methods*), the anomalous diffusion exponent *β* is commonly used to refer to whether normal (Brownian, *β* = 1) or anomalous diffusion (*β* ≠ 1) is observed. The dynamics of sub-diffusion and super-diffusion correspond to 0 < *β* < 1 and *β* > 1, respectively.

In Fig. 3*A*, we depicted an MSD analysis for the locomotion movements of three representative cells belonging to each species under absence of stimuli. The results of MSD analysis of the 700 experimental cells (see Fig. 3*B*, Fig. 3*C* and *SI Appendix*, Table S4) show that practically all trajectories exhibit strong anomalous migratory dynamics. For experimental trajectories, the variable *β*, which characterizes the behavior of the diffusion process, had a median/IQR value of 1.899/0.127 for *A. proteus* cells, of 1.878/0.168 for *M. leningradensis* cells, and of 1.845/0.146 for *A. borokensis* cells. These values suggest an anomalous super-diffusive process, a complex behavior which appears to govern the three groups of cell trajectories. The values for experimental trajectories of the anomalous diffusion exponent *β* ranged from 1.092 to 2.022, median/IQR value of 1.874/0.15, whereas the values for shuffled trajectories ranged from -0.007 to 0.006, with a median/IQR value of 10^-4^/0.002 (see Fig. 3*B* and *SI Appendix*, Table S5). A Wilcoxon test comparing anomalous diffusion exponents from shuffled showed that our results are extremely unlikely to be obtained by chance (p-value≅0, Z=32.392).

**Fig. 3.**
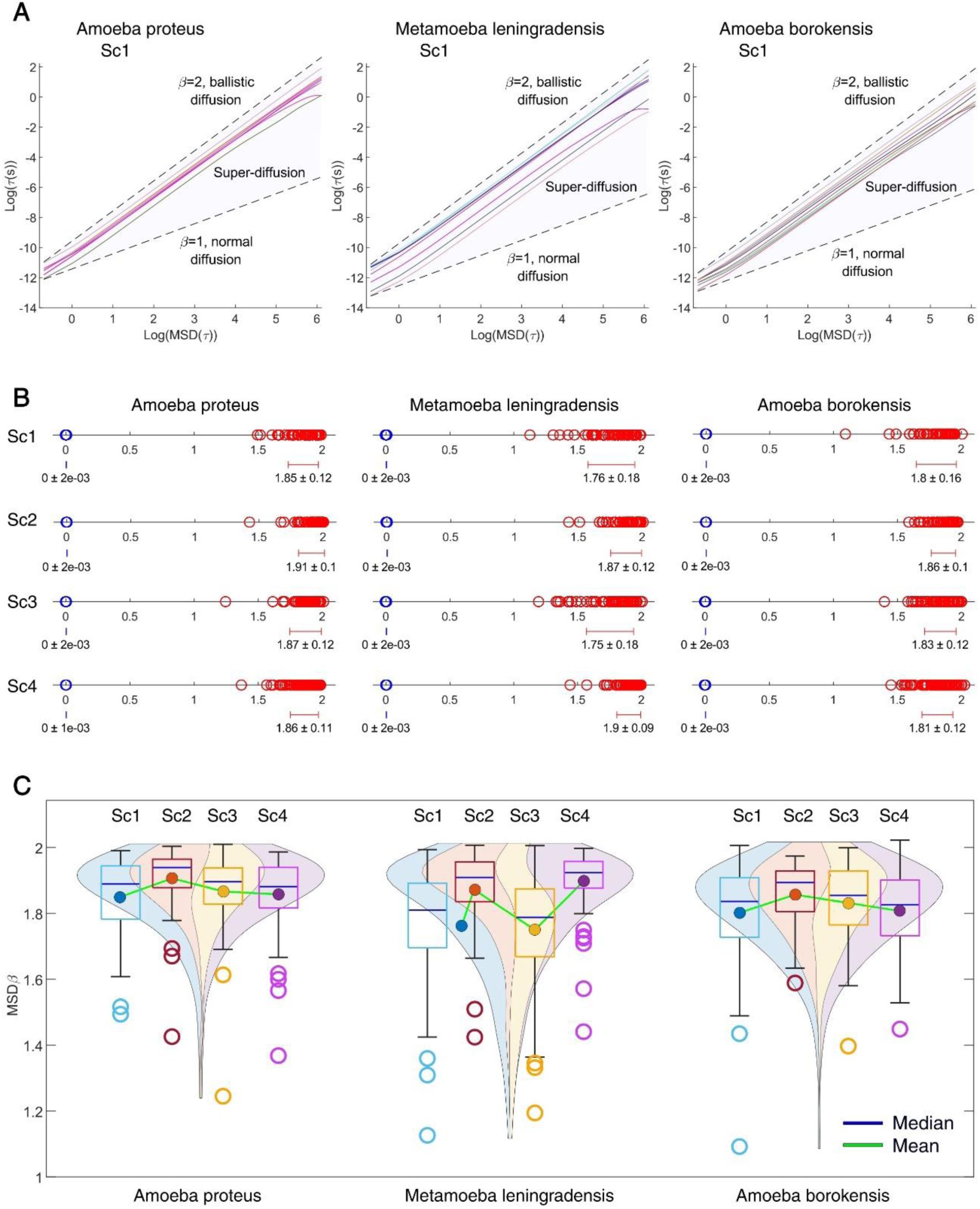
Strong anomalous migratory dynamics in cellular locomotion. *(A)* Graphics showing the value of the exponent β by fitting log-log plots of MSD as a function of the time interval τ, for 8 prototypic cells of each species (*A. proteus*, *M. leningradensis* and *A. borokensis*). β=1 indicates normal diffusion while β=2 indicates ballistic diffusion. The grey region defines the area of super-diffusion, which is a complex process with a high non-linear relationship to time, within which all the experimental values fall. *(B)* Diagram representing all values of the β exponents (and the overall average ± SD) for all cells of the three species in each experimental scenario (Sc1-Sc4) experimental values in red and shuffled values in blue. The shuffling step extinguished the long-term correlation structure, causing the sharp division between the experimental and shuffled value distributions (p-value≅0, Z=-32.3921) for all species and experimental conditions. *(C)* Estimated distribution, median and mean MSD β exponent values from experimental trajectories are illustrated using violin plots. “Sc1” Scenario One, absence of stimuli; “Sc2” Scenario Two, presence of an electric field; “Sc3” Scenario Three, presence of a chemotactic peptide gradient and “Sc4” Scenario Four, simultaneous galvanotactic and chemotactic stimuli.

### 7. Complexity and information in cellular migration

To assess the information content within locomotion trajectories we implemented the Approximate Entropy (ApEn), a robust approximation of the Kolmogorov–Sinai (K-S) entropy (45, 46), providing insight into the complex migratory behavior that emerges from the cellular system.

In Fig. 4*A* (*SI Appendix*, Tables S6 and S7), the results of the ApEn estimation for the 700 cellular trajectories are shown. The heatmaps display the Approximate K-S entropy for all experimental (upper row) and shuffled trajectories (bottom row) from each species, calculated for 72 different time windows (intervals) of increasing length (interval duration was increased by 25 seconds at every iteration). Intervals present ApEn values that vary from 10^-4^ (in blue) to 0.515 (red) for experimental trajectories and from 0.141 (in blue) to 2.126 (in red) for shuffled trajectories. These findings allow to observe that practically all the experimental series exhibit extremely low entropy.

**Fig. 4.**
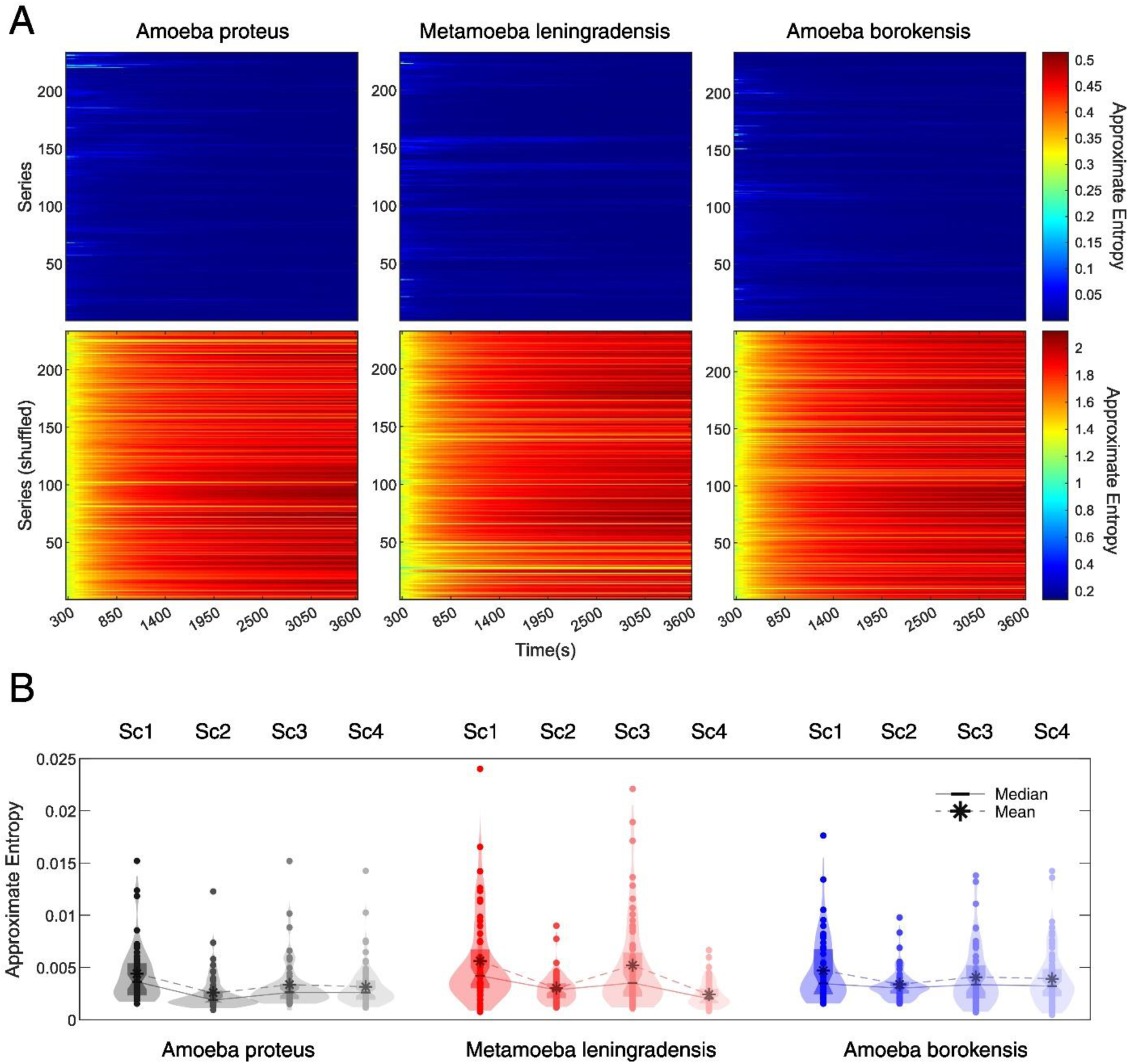
Complexity and information in cellular migration. *(A)* Heatmaps for the Approximate Entropy values of all 700 experimental (upper row panels) and shuffled (bottom row panels) cell trajectories from each species (*A. proteus*, *M. leningradensis* and *A. borokensis*). Each row in every panel corresponds to a single cell, while in the 72 columns the endpoint of the Approximate Entropy calculation is represented, increased in 25 seconds at every iteration. *(B)* Violin plots illustrate the estimated distribution, mean and median Approximate Entropy values for all experimental cell trajectories. “Sc1” Scenario One, absence of stimuli; “Sc2” Scenario Two, presence of an electric field; “Sc3” Scenario Three, presence of a chemotactic peptide gradient and “Sc4” Scenario Four, simultaneous galvanotactic and chemotactic stimuli.

In Fig. 4*B*, (*SI Appendix*, Table S6), the decrease in entropy that occurs from SC1 to SC2-SC4 denotes how in absence of stimuli (SC1) cells maximize entropy, but when there is a more defined directionality cellular trajectories become more focused (less entropic displacements). Specifically, we found that the ApEn values of experimental trajectories exhibited a narrow range of low values displaying a median/IQR of 0.003/0.002 for *A. proteus*, 0.003/0.003 for *M. leningradensis* and 0.003/0.003 for *A. borokensis*. The ApEn analysis for all experimental migratory trajectories obtained under the four scenarios showed a median/IQR ApEn value of 0.003/0.002 with values ranging from 10^-4^ to 0.024. In Fig. 4*B*, it is also showed that the high regularity and order observed in the experimental migration series vanished after shuffling. The ApEn values for shuffled trajectories displayed a range of very high values (from 1.252 to 2.126, median/IQR equal to 1.966/0.156) relative to the values for experimental trajectories (*SI Appendix*, Table S7).

The whole analysis confirms the presence of a complex structure characterized by high information in the move-step sequences in the migration trajectories of all cells. Furthermore, the statistical analysis revealed that this complex dynamic structure observed in the move-step trajectories was highly unlikely to occur by chance, as indicated by a p-value≅0 and Z=-32.392, results of a Wilcoxon test comparing the respective ApEn value distributions of experimental and shuffled trajectories.

### 8. Long-term memory effects in cellular migratory movements

Long-range memory effects or persistence is another main characteristic of the systemic cellular migratory movements in unicellular organisms (47, 48). The Detrended Fluctuation Analysis (DFA) (see *Materials and Methods*) is a well-known technique for measuring persistent effects in physiological time series.

For a given observation scale ℓ, DFA calculates the function (ℓ) to quantify the fluctuations of the time series around the local trend. If the time series displays scaling properties, then *F*(ℓ) ∼ ℓ^γ^ asymptotically, where γ represents the scaling exponent. This exponent is commonly estimated as the slope of a linear fit in the log(F(n)) versus log(ℓ) plot. Thus, γ serves as a measure of the strength of long-term memory effects and characterizes the underlying dynamical system. Specifically, values close to 0.5 indicate the absence of long-range correlations, while when 1.5< γ <2, the process exhibits positive long-range persistence (49) (Fig. 5*A*). Through the application of this quantitative method, we identified the presence of long-range persistence in all experimental trajectories (*SI Appendix*, Table S8), with a γ overall median/IQR value of 1.784/0.107. Specifically, the median/IQR DFA scaling parameter γ was found to be 1.795/0.087 for *A. proteus*, 1.785/0.138 for *M. leningradensis*, and 1.775/0.097 for *A. borokensis* (Fig. 5*B* and *SI Appendix*, Table S8), thus indicating that all the move-step trajectories exhibit trend-reinforcing memory.

**Fig. 5.**
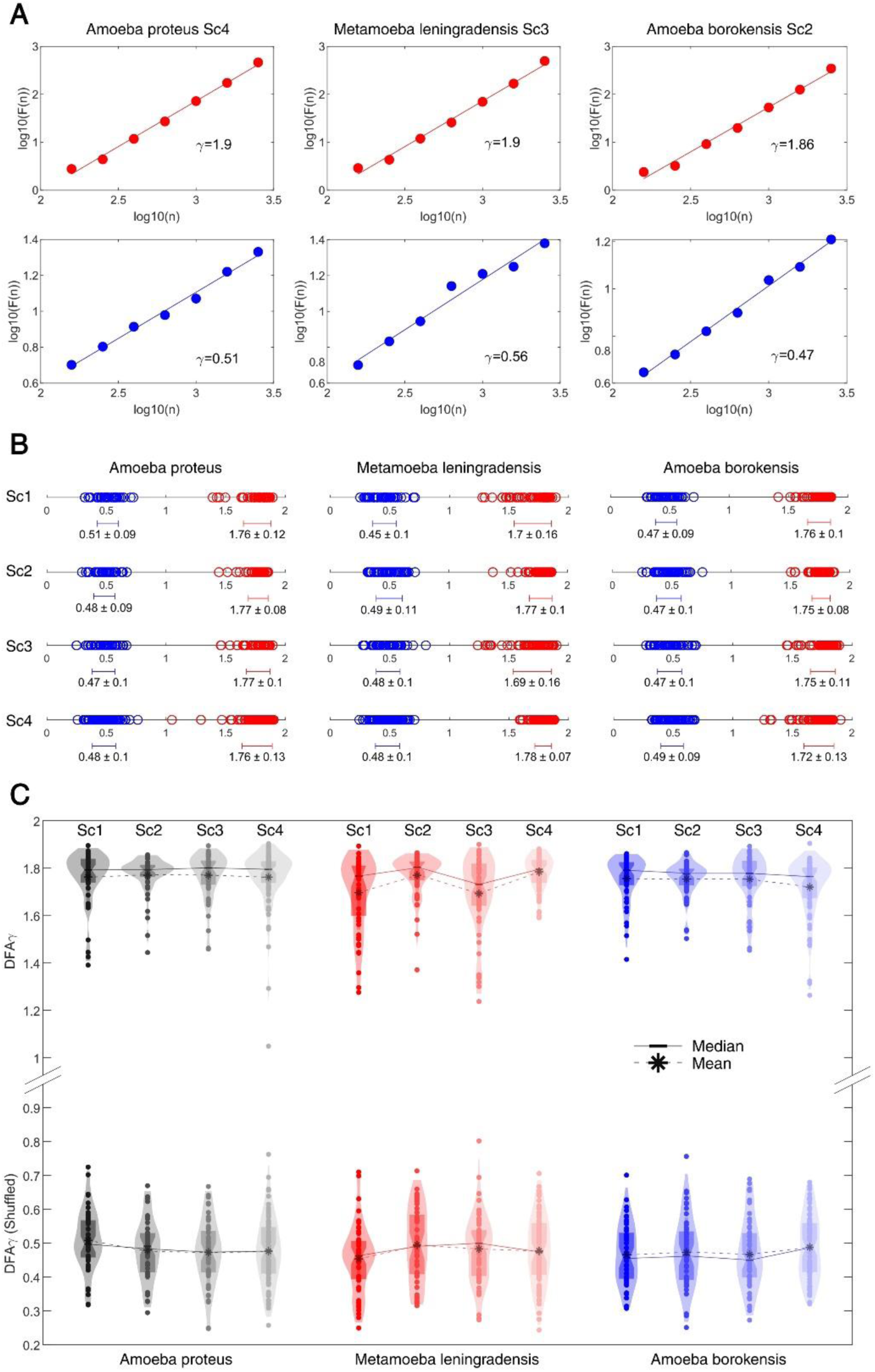
Long-term memory effects in cellular migratory movements. *(A)* Log-log plot of the detrended fluctuation parameter *F(n)* versus window size *n* for a prototype cell from each species. The scaling exponent γ was γ=1.9 for the *A. proteus* cell, γ=1.9 for the *M. leningradensis* cell and γ=1.86 for the *A. borokensis* cell, indicating long-range memory effects in all the species. The shuffling procedure removed the memory information contained in the original trajectories, causing γ values to drop to γ=0.51 for the representative *A. proteus* cell, γ=0.56 for the *M. leningradensis* cell and γ=0.47 for the *A. borokensis* cell. *(B)* Diagram displaying all the values of the scaling exponent γ (and the overall average ± SD) in all cells, separately for each species (*A. proteus*, *M. leningradensis* and *A. borokensis*) and experimental scenario (Sc1-Sc4). Values of the scaling exponent γ belonging to shuffled time series are depicted in blue, while exponents corresponding to experimental trajectories are depicted in red. *(C)* Violin plots illustrate the estimated distribution, mean and median scaling exponent γ values for all experimental (upper row) and shuffled (bottom row) cell trajectories. “Sc1” Scenario One, absence of stimuli; “Sc2” Scenario Two, presence of an electric field; “Sc3” Scenario Three, presence of a chemotactic peptide gradient and “Sc4” Scenario Four, simultaneous galvanotactic and chemotactic stimuli.

In order to assess the reliability of the DFA analysis, we conducted a random shuffling procedure on 700 time series. The results demonstrated that the strong correlation values observed in the experimental migration series vanished after shuffling (refer to Fig. 5*B*, Fig. 5*C* and *SI Appendix*, Table S9 for more information), with γ overall median/IQR of 0.477/0.135. This finding confirms that the complex locomotion structure, characterized by well-organized move-step sequences and persistent dynamics observed in the migration trajectories of the three cell groups, is not attributable to a random chance (p-value ≅ 0, Z=32.392).

### 9. Kinematic properties in cellular locomotion trajectories

To quantify some kinematic properties of the cell migration trajectories, we studied the Intensity of the response (IR), the directionality ratio (DR) and the average speed (AS) of amoebae (Fig. 6*A*, *B*, and *C*).

**Fig. 6.**
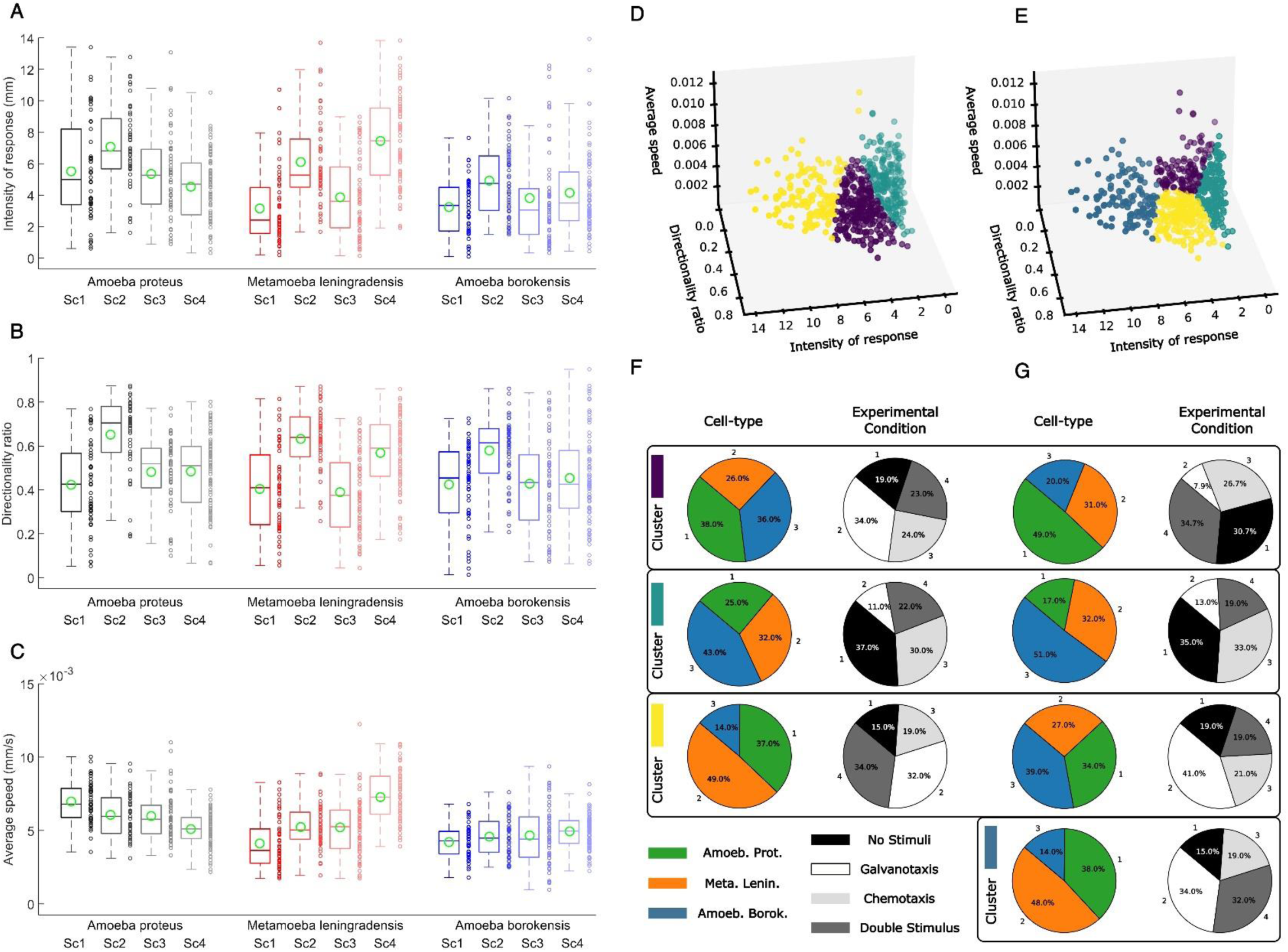
Kinematic properties and clustering analysis of cellular locomotion trajectories. *(A-C)* Group boxplots of the distributions of all experimental values for three main cytokinetic metrics, revealing basic data about the cell migration characteristics -*(A)* Intensity of the response, *(B)* Directionality ratio and *(C)* Average speed-of the three species used in this experiment (*A. proteus*, *M. leningradensis* and *A. borokensis*) under all four experimental scenarios (Sc1, Sc2, Sc3 and Sc4). *(D-G)* Unsupervised clustering was performed using *k-means* on experiments defined by 3D vectors of three metrics. *(D)* Three clusters are depicted (yellow, purple, and green), with each dot representing a different cellular motion experiment (combining three cell types and four experimental scenarios). *(E)* Similar to *(D)* but with four clusters. Note that the clusters in Panel *(D)* do not have the same colours as in Panel *(E)*, although three of the four colours are the same. *(F)* Characterization of the three clusters (panel *D*) in terms of cell types and experimental conditions. *(G)* Similar to *(F)* but with four clusters (panel *D*).

The IR is associated to the space explored by the cell, and in particular, we quantified the module of the trajectories to represent the strength of the response. In this case, the median/IQR IR was 5.322/3.5 for *A. proteus*, 4.961/5.053 for *M. leningradensis* and 3.575/3.072 for *A. borokensis*. Next, we studied the DR, which quantifies the trajectory straightness, ranging between 0 (for fully curved trajectories) and 1 (for fully straight trajectories), by considering the start and end point of the trajectory. The values ranged between 0.052 and 0.875 (median/IQR 0.525/0.307) for *A. proteus*, 0.043 and 0.871 (median/IQR 0.508/0.315) for *M. leningradensis* and 0.013 and 0.949 for *A. borokensis* (median/IQR 0.469/0.296).

Finally, we calculated the average speed (AS) of the trajectories, which ranged between 0.002 and 0.011 mm/s (median/IQR 0.006/0.002) for *A. proteus*, 0.002 and 0.012 mm/s (median/IQR 0.006/0.003) for *M. leningradensis* and 0.001 and 0.009 mm/s (median/IQR 0.005/0.002) for *A. borokensis*.

As it can be observed from the p-values derived from Kruskal-Wallis analyses there is a remarkable variability regarding kinetic properties, both between species (for example, the p-values comparing the IR, DR and AS in Scenario 4 were 10^-13^, 10^-4^ and 10^-16^, respectively) and between scenarios (for example, the p-values of *A. proteus* for IR, DR and AS compared among all four scenarios were 10^-6^, 10^-9^ and 10^-10^, respectively); for more information, see Fig. 6 and *SI Appendix*, Table S10.

Fig. 6*D-G* shows a clustering analysis performed on all kinematic properties considered. Each cluster was characterized by the proportion of cell types and experimental condition present in each group. The performance of the obtained clustering solution was assessed using the Silhouette Coefficient, which estimates all the differences between intra-cluster points minus the distances between inter-cluster points. A higher Silhouette index indicates a model with better defined clusters. The implementation was achieved using the silhouette score implemented in Scikit-Learn. The three clusters identified in Fig. 6 yielded a Silhouette coefficient of 0.366. Similarly, the four clusters identified provided a Silhouette coefficient of 0.369. Another alternative clustering analysis by means of a hierarchical agglomerative method (*SI Appendix*, Fig. S5) showed that the distinction between cell-types or experimental conditions remains unchanged when varying the clustering strategy, which indicates the robustness of the findings.

The high variability of the statistics of the kinematic parameters, combined with the cluster analysis, indicates a high level of heterogeneity in any of the three metrics used. This heterogeneity exists both among cell types and among different experimental conditions and suggests that the behavior is individual in each one and hence they cannot be separated into groups.

### 10. Dynamic structure in migratory movements

Finally, we have represented all the main metrics considered in our study, such as RMSF Alpha, RMSF correlation time (measured in move steps or in minutes), DFA Gamma, MSD Beta, and Approximate Entropy by comparing them with those data obtained in the corresponding shuffling procedures (Fig. 7*A-C*). As evident in each panel, the metrics’ shuffled and non-shuffled values could be distinctly grouped and differentiated. This suggests that the inherent systemic information structure was disrupted during the shuffling process. The trajectories of the experimentally observed cells are completely differentiated from the cells whose trajectories lost systemic properties.

**Fig. 7.**
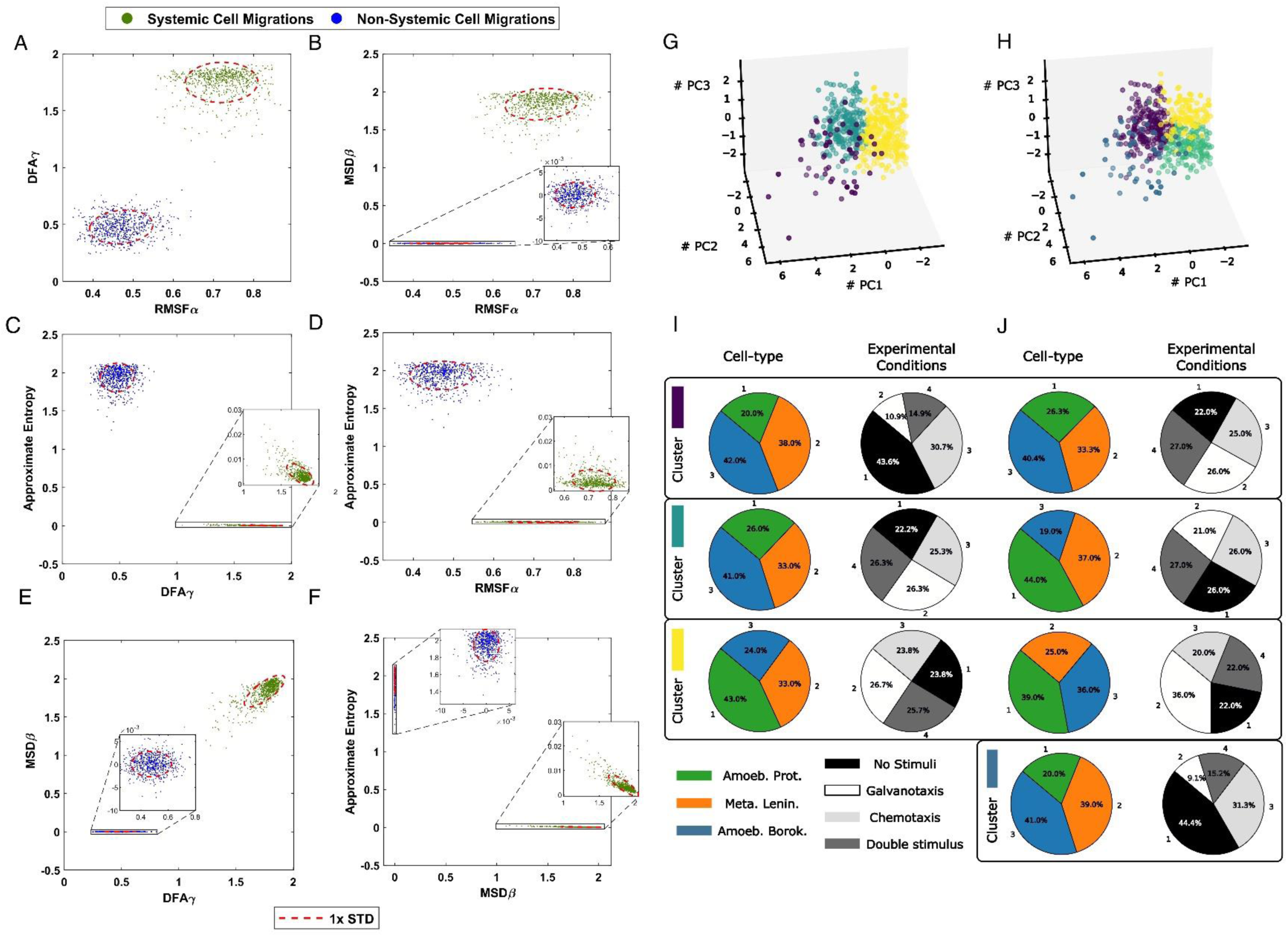
Systemic metrics and clustering analysis of cellular migratory displacements. *(A-F)* Analysis of the 700 experimental cell trajectories with the main metrics considered in our work such as RMSF Alpha, RMSF correlation time, DFA Gamma, MSD Beta, and Approximate Entropy is compared with corresponding shuffled data in which the systemic informational structure was completely lost. *(G-J)* Clustering analysis of cellular motion experiments does not distinguish between cell types and experimental conditions when using non-linear advanced movement metrics. Similar to Fig. 6 *(D-G)* but defining each experiment using the 5D main vectors of metrics. K-means clustering was performed using the first three principal components (PC1, PC2, and PC3), which accounted for 92.54% of the total variance. The interpretation of the different panels is the same as in Fig. 6, but the variables used for clustering are now different.

In Fig. 7*G-J* a clustering analysis of the main metrics was performed. The three clusters related to cell type identified in Fig. 7 yielded a Silhouette coefficient of 0.347. Similarly, the four clusters identified related to experimental conditions provided a Silhouette coefficient of 0.311. This unsupervised clustering analysis combined to the small variability of the results, appears to be quite homogeneous, regardless the cell type or scenario considered, since its quantitative aspects show no dependency on either cell type or experimental condition. Moreover, we utilized an alternative approach, specifically hierarchical agglomerative clustering (*SI Appendix*, Fig. S6), and the resulted profile remained consistent when the different clustering methods were applied, highlighting the robustness of the results.

These findings suggest the emergence of a highly intricate dynamic structure within the migratory patterns of all examined cell trajectories. Furthermore, this structure appears to be an inherent aspect of cell locomotion, irrespective of species or environmental conditions. Indeed, cluster analysis of all quantitative parameters revealed no reliance on either cell type or experimental context, suggesting the potential universality of this behavior.

## Discussion

Cellular migration is a cornerstone issue in many essential physiological and pathological processes. Despite its enormous relevance, how cells control their locomotion movements is a still unresolved question which represents a fundamental challenge in contemporary biology.

For years, the scientific attention has been focused on the individualized study of the diverse molecular parts involved in cell migration; however, locomotion movements have never been regarded as a systemic process that operates at a global cellular scale.

Here, we have addressed the integrative systemic dynamics involved in the regulation of directional motility. To this end, we have studied the migratory displacements of 700 single cells, belonging to three different species (*Amoeba proteus*, *Metamoeba leningradensis* and *Amoeba borokensis*), in four different scenarios: in absence of stimuli, under chemotactic gradient, in an electric field and under complex external conditions such as simultaneous galvanotactic and chemotactic gradient stimuli. The experimental trajectories, obtained on flat two-dimensional surfaces, have been quantitatively studied using a multidisciplinary approach to understand how integrative systemic forces drive the locomotion movement of cells.

First, we have analyzed the long-range interdependence in the move-steps using the “rmsf” method, a classical Statistical Mechanics approach based on the concepts developed by Gibbs and Einstein (39, 40), later developed and used to quantify biological processes (50–52). Our results demonstrate that each move-step ahead at a given point is strongly influenced by its preceding displacements, indicating that strong dependences of past movements lasting approximately 1,137 move-steps over periods averaging 9.5 minutes do exist. Practically, all the 700 unicellular organisms analyzed in the four experimental conditions exhibited non-trivial long-range correlations in their directional trajectories, which represents a key characteristic of the systemic dynamic movements emerging in the cell system (47).

Anomalous behavior is another characteristic of the migratory movements that we have identified using the MSD method also proposed by Einstein (43). Migratory dynamics that do not result in a linear MSD can be considered as non-trivial. Specifically, the anomalous nature of cell migration can be detected by super-diffusion, a physical phenomenon detected in the trajectories of all the 700 cells analyzed. Likewise, the MSD is a proxy for the surface area explored by the cell over time and is a measure related to the overall migration efficiency. The cellular displacements analyzed correspond to efficient movements during the exploration of the extracellular medium (41, 42, 53). The strong manifestation of the anomalous nature of cell migration can be caused by temporal memory effects as a consequence of the non-trivial long-range correlations in the cellular move-steps (53–56).

We have also quantified the regularity and unpredictability of the fluctuations over the migratory displacements (45, 46). The obtained results show high levels of information in all the analyzed trajectories. This finding, together with the previous ones, confirms the presence of a very complex structure in the migratory move-step sequences. Entropy is directly related to the complexity of the system dynamics, and the very low level of entropy in the directional movements indicates that the migration patterns are organized on a level of complexity that is above the individual components of the cell system. Such complex dynamic structure observed in the trajectories was highly unlikely to occur by chance (practically, p-value ≅ 0).

In addition, we have verified the presence of long-term memory effects in the cellular migratory movements. The results of the DFA fluctuation analysis (57) show that the scaling exponent γ displays a total average (± SD) of 1.749 ± 0.117, indicating that the move-steps trajectories exhibit a trend-reinforcing memory, that is, if the directional movements in the past show an increase in a set of their move-step values it is very likely to be followed by an increasing trend in the future; and vice versa, a decreasing trend in the past, is likely to be continued in the future. In other words, the evolution of the cell system trajectories is strongly influenced by previous system movements over long periods of time. Long-term memory is a key concept, closely related to long-term correlations, widely developed in Physics with a robust and formal Mathematical construction. Therefore, the results we have obtained with DFA also validate the presence of strong long-range correlations in the locomotion movements.

The quantitative studies carried out here unequivocally show that a very complex dynamic structure emerges in the migratory movements of all the analyzed cell systems. Such structure is characterized by highly organized move-step sequences with very low level of entropy and high information, non-trivial long-range interdependence in the move-steps, strong anomalous super-diffusion dynamics, long-term memory effects with trend-reinforcing behavior, and efficient movements to explore the extracellular medium. The outstanding detected dynamic structure underlies all the migration trajectories of 700 cells of three different species analyzed under the four experimental scenarios. On the other hand, the results of the two types of clustering analysis performed suggested the potential universality of this complex systemic structure in the cellular locomotion movements.

These essential characteristics of the locomotion movements are a consequence of the self-organized dynamics intrinsic to all unicellular organisms. Cells are sophisticated systems conformed by the mutual interactions of millions of molecules and hundreds of thousands of macromolecular subcellular structures, following an intricate interplay that challenges the human mind (58). They are open systems that operate far from the thermodynamic equilibrium and exchange energy-matter with the environment (59–61). Under these conditions, non-linear enzymatic interactions and irreversible metabolic processes allow the cell system to become spatially and temporally self-organized (58, 62–66). If cells reach the equilibrium, their sophisticated dynamic functionality and molecular order disappears and they die.

Briefly described, the essential energy-matter flow generates a negative entropy variation inside the cell which corresponds to an emergent positive increment in the information of the system (67). Such information increases the complexity, producing collective functional patterns, highly ordered macrostructures, and complex self-organized behaviors as for instance molecular-metabolic rhythms and spatial traveling waves (61, 68).

These emergent non-equilibrium molecular dynamics supported by permanent energy dissipation (continuously exporting entropy to the external medium) are known as self-organized dissipative structures (69, 70). The principles of self-organization through energy dissipation were conceived and developed by the Nobel Prize Laureate in Chemistry Ilya Prigogine (38).

Intensive studies over the last six decades have demonstrated that cells are very complex self-organized dissipative systems (58, 60, 66, 67, 71–73) in which integrated processes and systemic properties at different levels of organization and complexity do appear. At a basic level, dissipative molecular behaviors emerge, for example, in shaping actin polymerization waves involved in the cytoskeleton activities during cell migration (74, 75), in self-organized oscillations in actin networks (76), in myosin dynamics (77), in microtubular behavior (78), and in intracellular calcium rhythms (79). At the highest level, complex systemic properties such as directional mobility, integral growth, reproduction, sensitivity to the external medium, adaptive responses, and evolution do occur (58). To note, these strong emergent properties cannot be found in their individual molecular components or in their single molecular-metabolic processes (66).

Systemic dynamics are emergent integrative processes in all unicellular organisms, and such behaviors are a consequence of the self-organization of the biochemical system as a whole (80). From collective metabolic-molecular constituents, all of them interacting nonlinearly with each other, emerge basic coherent self-organized structures and functional ordered patterns which originate a cell system that increases at different levels its structural and functional complexity driven by energy dissipation and molecular information processing (58, 63, 64).

A critical attribute of these dissipative self-organized systems is the interacting dynamics exhibiting long-range correlations (81–83). In his Nobel Lecture dissertation, Ilya Prigogine stressed that the main feature of systemic dynamics in dissipative systems is the emergence of long-range correlations (38).

On the other hand, long-term correlations have also been showed in different metabolic processes such as calcium-activated potassium channels (84), intracellular transport pathway of Chlamydomonas (85), glycolytic studies (86–89), NADPH series (90), and metabolic networks (91,92), attractor dynamics (93), neural activity (94). In additions, complex emergent systemic behaviors have been observed in cellular locomotion movements (95,96).

Until now, cell migration has never been considered as a systemic property, but in this work we have shown evidences that indicate the existence of complex integrative dynamic processes at a global cellular scale involved in directional motility.

First, we have highlighted how a relevant number of experimental studies have verified that most of the fundamental cellular physiological-molecular activities in the cell are implicated in their locomotion movements, which is indicative of a phenomenon that operates at a systemic level.

We have also collected previous evidences found over the last decades showing that unicellular organisms constitute self-organized systems characterized by emergent molecular-metabolic processes and integrative responses operating at a systemic level. These self-organized processes are mainly characterized by exhibiting coherent behaviors with long-range correlations, a ubiquitous characteristic of all dissipative systems.

Our quantitative analysis has shown that all cell systems analyzed here display a kind of dynamic migration structure characterized by non-trivial long-range correlations, strong anomalous super-diffusion dynamics, long-term memory effects, highly organized move-step sequences with very low level of entropy and high information, and efficient movements to explore the environment. Such characteristics correspond to critically self-organized systems. The locomotion trajectories change continuously, since they exhibit random magnitudes that vary over time, but these stochastic movements shape a dynamic structure whose defining characteristics are preserved in all the conditions analyzed. This movement structure corresponds to complex behavior belonging to a self-organized cell system, in which the emergent systemic dynamics drive the locomotion movement of cells.

Cellular locomotion seems to be regulated by complex integrated self-organized dynamics, carefully regulated at a systemic level, which depends on the cooperative non-linear interaction of most, if not all, cellular components. The systemic properties responsible of this locomotor behavior are not found specifically in any of their singular molecular parts, partial mechanisms, or individual processes in the cell.

Our quantitative experimental results together with the remarkable amount of evidences provided by other scientific analyses seem to indicate that migration is a systemic property of cells. This fact does not invalidate the importance of studying the influence of individual metabolic-molecular pathways on cell migration.

Cell migration is a central issue in many human physiological and pathological processes. We consider that new researches combining migratory systemic dynamics with molecular studies are crucial for the development of next-generation, efficient cellular therapies for migration disorders. In addition, the findings presented here open a new perspective for improving the conceptual framework of the cell, the most complex molecular system in Nature.

## Materials and Methods

### Cell cultures

The three species of free-living amoebae were cultured in Ø100 x 20 mm Petri dishes (Corning^®^ CLS430167) at 21°C. *Amoeba proteus* (Carolina Biological Supply, #131306) and *Metamoeba leningradensis* (CCAP: Culture Collection of Algae and Protozoa, Oban, Scotland, United Kingdom, catalogue number 1503/6) were grown in Chalkley’s simplified medium (36) (NaCl, 1.4 mM; KCl, 0.026 mM; CaCl2, 0.01 mM) with heat-treated wheat grains. *Amoeba borokensis* (ACCIC (97): Amoebae Cultures Collection of Institute of Cytology in Saint Petersburg) were grown in Prescott & Carrier media (98) and fed every fourth day 0.5 ml of a mixed *Chilomonas sp.* (Carolina Biological Supply^®^ #131734) and *Colpydium sp.* (ACCIC culture. *Chilomonas sp.* and *Colpydium sp.* were grown in Prescott & Carrier media as well.

### Experimental set-up

A schematic drawing of the experimental set-up is shown in *SI Appendix*, Figs. S1 and S2. Two electrophoresis blocks (BIO-RAD Mini-Sub cell GT) were interconnected by two ∼12 cm long agar bridges (2% agar in 0.5 N KCl in) and the first one was plugged into a BIO-RAD Model 1000/500 power supply unit. Atop the central elevated platform of the second block, a custom-devised experimental glass chamber was placed. By using agar bridges and removing all the electrodes from the second electrophoresis block we avoided direct contact between the media, where the cells moved, and any metallic electrodes, thus preventing ionic contamination.

The experimental glass chamber consisted of a modified 25 x 75 x 1 mm standard glass slide and three additional, smaller glass pieces (one central piece measuring 24 × 3 × 0.17 mm and two flanking pieces of 24 × 40 × 0.17 mm each) crafted by carefully trimming three 24 x 60 x 0.17 mm cover glasses (*SI Appendix*, Figs. S1 and S2). To build the modified standard glass slide, two 24 x 60 x 0.1 mm cover glasses were stuck using silicone to a standard glass slide and let dry for 24h, whereupon the 20 × 60 × 0.17 mm overhanging parts from both cover glasses were cut off, leaving two 4 × 60 × 0.17 mm glass sheets as the chamber’s sideway walls. All the experiments were conducted within the experimental glass chamber, which allowed to establish a laminar flow when closed and to place or extract the cells when opened.

Fresh agar bridges were used in every experimental replica to avoid contamination and conductivity loss (leak of KCL into the simplified Chalkley’s medium, which has a much lower osmotic concentration than the agar bridges). Furthermore, one electrophoresis block was used exclusively for chemotaxis and another one for galvanotaxis, while the whole set-up (both the electrophoresis blocks and the glass chamber) was scrupulously cleaned and reassembled after each experimental replica.

Ahead of every experiment, the modified glass slide was adhered to the top of the central platform of the second electrophoresis block using a droplet of olive oil to prevent both the medium and the electric current from passing beneath the experimental glass chamber. Following this, the 24 × 3 × 0.17 mm central glass piece was gently placed atop the middle of the modified glass slide. Next, amoebae were washed in fresh simplified Chalkley’s medium and placed beneath the central piece of the glass chamber, where they were left to attach to the surface of the modified glass slide for ∼2 minutes. This step needs to be performed in less than 15 seconds, for the amoebae will immediately start attaching to the inner surface of the plastic micropipette tip, and any further pipetting will henceforth potentially damage their cellular membrane. *Metamoeba leningradensis* cells attached faster and stronger to the plastic micropipette tips than the other two species, thus being especially susceptible to being damaged. Subsequently, the two 24 × 40 × 0.17 mm lateral sliding glasses were laid flanking the central piece and slightly overhanging the electrophoresis block wells. Each well was carefully filled with 75 ml of clean simplified Chalkley’s medium, and the two 24 × 40 × 0.1 mm lateral sliding glasses were gently poked down using a micropipette tip until they contacted the medium, which spread beneath them by surface tension. Finally, the two lateral glass pieces were longitudinally slid till they touched the central piece where the amoebae laid, thus closing the glass chamber, and establishing a connection between the media in both wells.

All experimental replicates were performed with a maximum 9 cells and lasted for exactly 30 minutes, during which cell behavior was recorded using a digital camera. In preparation for each experiment, amoebae were starved for 24 hours in fresh, nonnutritive simplified Chalkley’s medium. Only cells deemed healthy (motile and rod-shaped) were chosen for the experiments. Deviations from optimal culture and experimental conditions, as well as mechanical issues in the recording system, rendered ⪅ 9% of the experimental replicates invalid. Those replicates have not been considered in this investigation.

### Galvanotactic stimulus

The galvanotactic stimulus is traumatic for the amoebae, and thus can affect their response. Four key steps were taken to minimize those effects by ensuring optimal current intensity and voltage in all the experiments where a galvanotactic stimulus was applied: 1. The power supply unit was programmed to keep the voltage at 60V; 2. The flow sectional area of the experimental glass chamber was adjusted by modifying the amount of silicone used to glue the longitudinal walls of the modified glass slide, which determined their height; 3. A variable 1 MΩ resistor and a microammeter were installed in series, in that order (*SI Appendix*, Fig. S3), and the intensity of the electric current was manually corrected when needed in real time by turning the variable resistor’s screw to tune the global resistance of the set-up; thus, a stable 60V electric potential and optimal current intensity values of 70-74 µA (*A. proteus*), 70-80 µA (*M. leningradensis*) and 68-75 µA (*A. borokensis*) were kept throughout the galvanotactic experiments; 4. Lastly, the current was stopped immediately once the 30-minute image recording finished.

We found some amoebae populations displayed an anomalous, inverted or even null response to the galvanotactic stimulus. We therefore carried out a 5-minute galvanotactic test prior to any experiments where amoebae were exposed to galvanotaxis for the first time, using intensity and voltage values within the optimal ranges provided.

### Chemotactic stimulus & gradient calculation

To establish the chemotactic peptide gradient, we added 750 µl of 2 × 10^−4^ M nFMLP (Sigma-Aldrich, #F3506) to one well of the second electrophoresis block to obtain a working peptide concentration of 2 × 10^−6^ M. The medium in that well was stirred immediately after adding the peptide to properly mix the peptide until the amoebae began to respond to it.

To assess the nFMLP peptide gradient concentration, an experiment was carried where 60 µL of medium were sampled at 0, 2, 5, 10, 15, 20, and 30 minutes since the addition of the nFMLP peptide. Samples were taken from the center of the experimental glass chamber through a small gap opened between the central glass piece and one of its flanking glass pieces by slightly moving this sliding lateral glass. Known fluorescein-tagged peptide concentration values from a standard curve were used to extrapolate nFMLP concentration at each time point (*SI Appendix*, Fig. S4). Two samples were taken at each time point, and the whole procedure was replicated three times, yielding a total of 6 measurements for each time point. Fluorescence was measured at 460/528 excitation/emission wavelengths on 96 well glass bottom black plates (Cellvis, #P96-1.5H-N) using a SynergyHTX plate reader (BIOTEK) following the protocol established by Green and Sambrook (99).

### Experiment recording and cell tracking

Experiments were recorded with a digital camera attached to an SM-2T stereomicroscope. Two frames were captured every second for 30 minutes (3600 frames). Individual cell movements (tracks) were manually digitized using the TrackMate (100) plugin from FIJI (ImageJ) and saved as a list of (x, y) coordinate tuples. Manual tracking was chosen over automated and semi-automated alternatives due to the well-known imprecision of such tools (101).

### Approximation Entropy calculation

Approximation Entropy (ApEn) is a statistic developed by Pincus et al. in 1991 (45) as a measure of relative regularity in data series. In Fig. 4*A*, Approximation Entropy ApEn values corresponding to intervals of length <300 data points were not represented in the heatmaps, as ApEn requires at least 10^m^ data points (m=2 in our calculations) to produce meaningful results (45), and these are known to be unreliable in time series of length ≤200 points (102).

### Clustering numerical experiments

The clustering analyses were implemented using our custom code in Python 3.9.13 and Scikit Learn 1.0.2. Combining all experiments across the three cell types (*M. leningradensis*, *A. Proteus*, and *A. borokensis*) and the four experimental conditions (no stimuli, galvanotaxis, chemotaxis, and simultaneous galvanotactic and chemotactic stimuli), unsupervised clustering was performed on all experiments of cellular movement, each defined by a vector of different movement metrics (MM). In the first clustering analysis (Fig. 6), experiments were characterized by the 3D vector of MMs: Intensity of the response (mm), Directionality Ratio, and Average Speed (mm/s). In the second clustering analysis (Fig. 6), experiments were identified using the 5D vector of MMs: RMSF Alpha, RMSF correlation time (move-steps), DFA Gamma, MSD Beta, and Approximate Entropy. Before performing the clustering analysis, and to ensure a comparable visualization to the first analysis, we first applied Principal Component Analyses (PCA) and subsequently applied the clustering strategy to the first three components (PC1, PC2, and PC3), which explained the 92.54% of the total variance in the data. For the second analysis, the clustering results did not change when using RMSF correlation time measured in minutes instead of move steps, as both metrics are highly correlated. Although the best clustering solution (measured by the Silhouette Coefficient) was obtained in both analyses for the number of two clusters, Figs. 6 and 7 display 3 and 4 clusters with the purpose of exploring potential associations with, respectively, cell type and experimental condition. After normalizing the different values using z-scores across all possible experiments, we applied the *k-means* algorithm to obtain 3 and 4 clusters. Subsequently, each cluster was characterized by the proportion of cell types and experimental condition present in each cluster. The performance of the obtained clustering solution was assessed using the *Silhouette Coefficient*, which estimates all the differences between intra-cluster points minus the distances between inter-cluster points. A higher Silhouette index indicates a model with better defined clusters, while a smaller index suggests more heterogeneity in the data, and no well-separated groups. The implementation was achieved using the silhouette score implemented in Scikit-Learn library in Python. To show robustness of the clustering solution and that our results were not dependent on the clustering strategy employed, we repeated the same analyses as in Figs. 6 and 7 but using *hierarchical agglomerative* clustering and depicting the solutions of 3 and 4 clusters (*SI Appendix*, Figs. S5 and S6).

### Statistical analysis

First, the normality of the distribution of our quantitative data was assessed using the Kolmogorov-Smirnoff test for single samples. Given that normality was rejected, the significance of our quantitative results was estimated through the Kruskal-Wallis test for groups and the Wilcoxon rank-sum test for pairs. Since these two are non-parametric tests, results are represented as median/IQ instead of mean ± SD. In addition to the p-values the Z statistics have been reported.

## Data and code availability

The data and code generated by this study are publicly accesible from the Zenodo repository at https://doi.org/10.5281/zenodo.10974258.

Any additional information required to reanalyze the data reported in this paper is available from the lead contact upon request.

## Acknowledgments

We would like to thank Florentino Onandía Yague for valuable advice related to the experimental setup. This work was supported by grant US21/27 from the University of Basque Country (UPV/EHU) and Basque Center of Applied Mathematics. In addition, this work was supported by Basque Government funding, grant IT456-22.

## Author contributions

JC-P, and MF: performed the experiments; CB, MF and IMDF: designed the setup; CB methodology with experimental glass chamber; MF and JC-P: cell cultures and cellular behavior advice; JC-P: performed the digitalization of trajectories; IM, JMC, IMDF: designed quantitative analysis; JC-P,IM, JCM and BC-P: performed the quantitative studies; GPY, JIL, AP-S, JC-P, IM, JMC and IMDF: analysis and design of the research mapping; all authors wrote the manuscript and agreed with its submission; IMDF: conceived, designed and directed the investigation.

## Supporting information

Appendix 01 (PDF)

## Supplemental information

**Figure S1.**
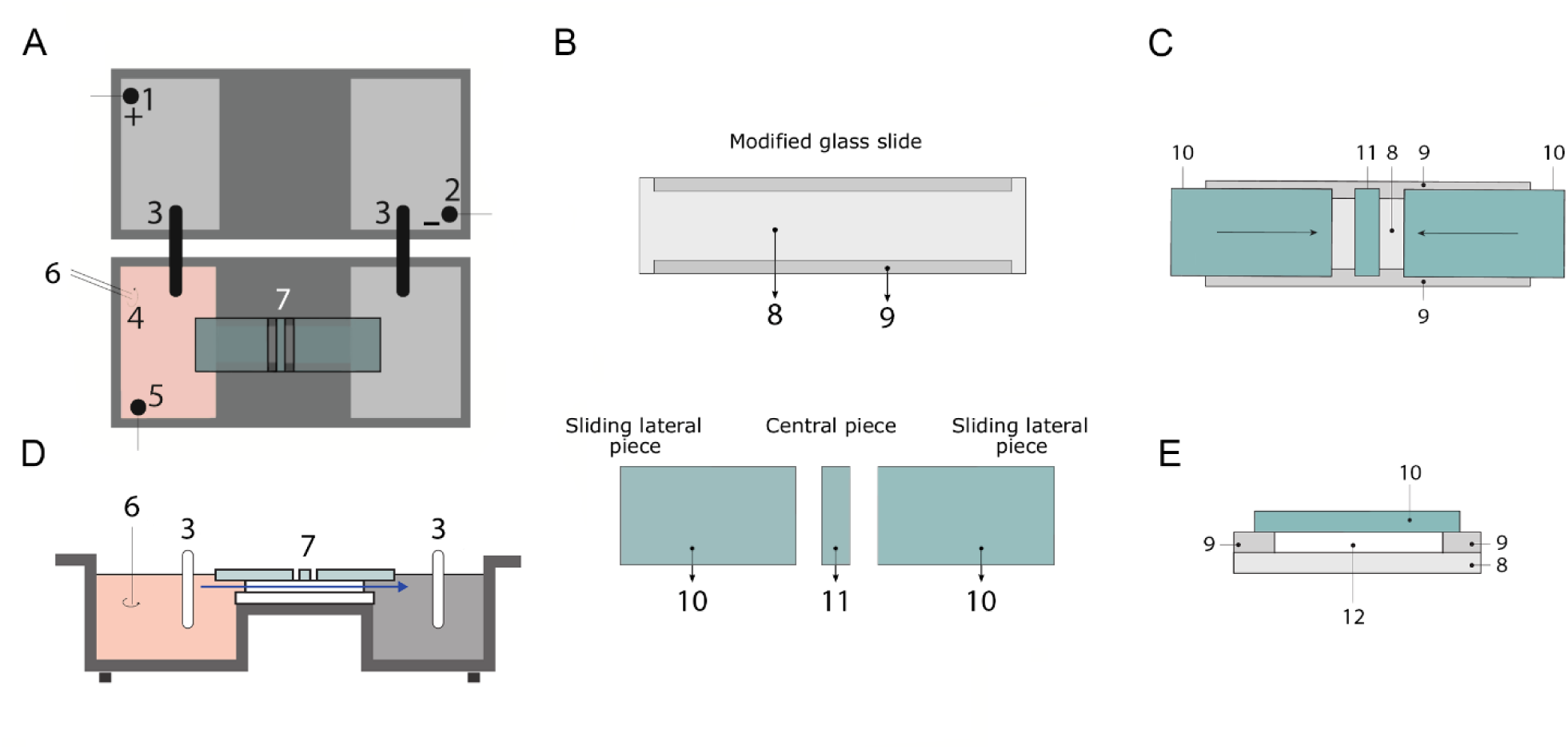
Experimental set-up layout. (A and D) Top and lateral views of the experimental setup. 1: anode; 2: cathode; 3: agar + KCl bridges; 4: chemotactic peptide; 5: electrode used to probe the electric field; 6: stirrer used to properly mix the peptide; 7: experimental glass chamber, the blue arrow signals the trajectory and direction of the laminar flow. (B) Top view of the glass pieces that compose the experimental glass chamber. 8: 75 × 25 mm standard glass slide; 9: longitudinal trimmed glasses; 10: sliding lateral glasses; 11: central piece of glass underneath which the cells are placed. (C) Top view of the experimental glass chamber. (E) Axial section of the experimental chamber. 12: flow sectional area. The experimental chamber can be opened and closed by longitudinally displacing #10, allowing to place or remove cells when open and establishing a laminar flow of medium through #12 when closed (See *Materials and Methods* for further details).

**Figure S2.**
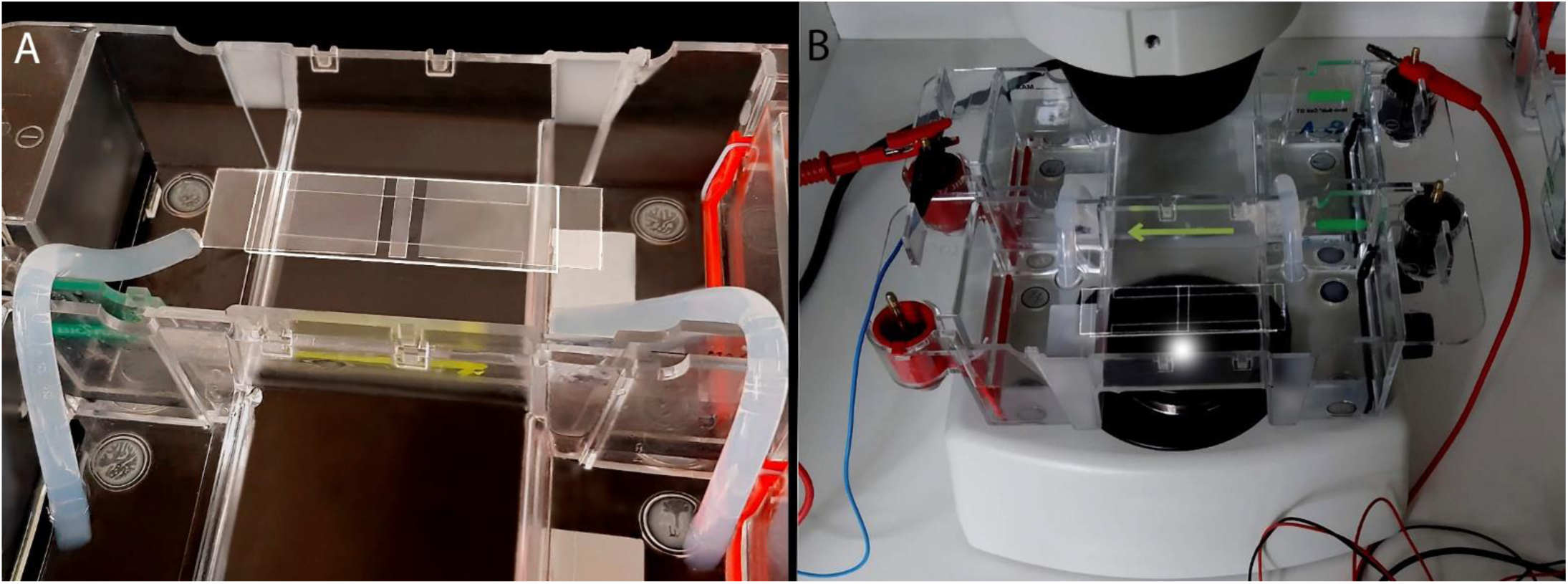
Experimental set-up, related to Figure S1. (A and B) All elements sketched and described in Figure S1 are depicted here under experimental conditions.

**Figure S3.**
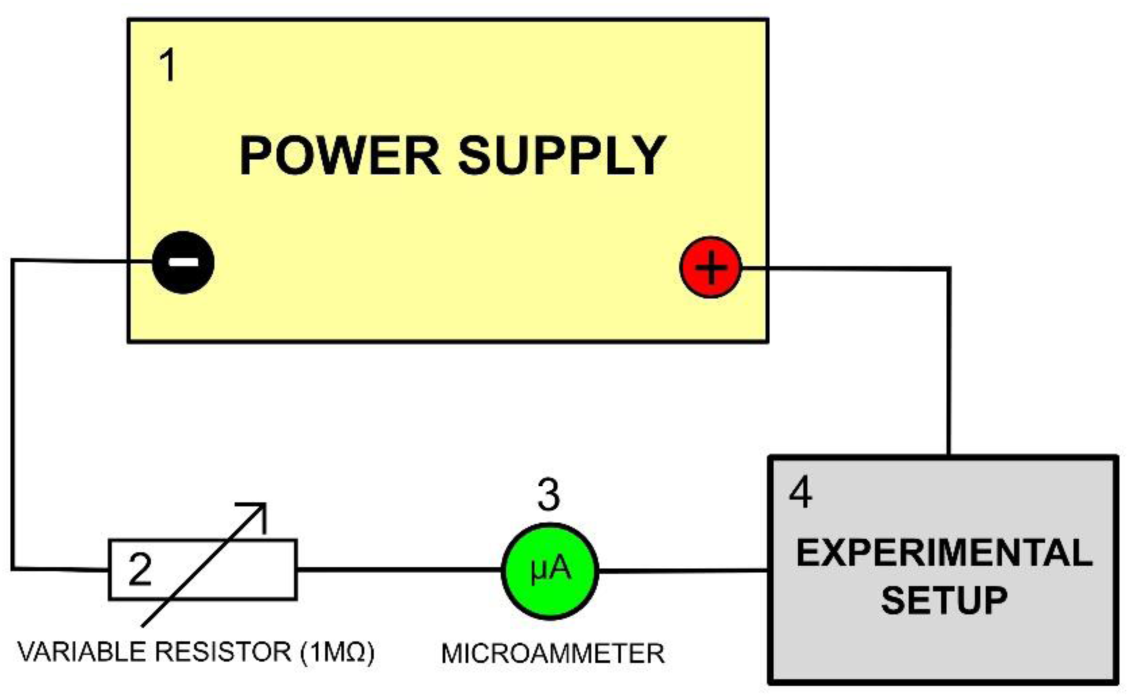
Auxiliary electric circuitry. This circuit allows the electric current that circulates through the experimental system to be monitored and adjusted as necessary. 1: BIO-RAD model 1000/500 power supply unit; 2: 1 MΩ variable resistor to adjust the intensity of the electric current; 3: Microammeter to monitor current intensity flowing through the system; 4: Experimental set-up (see Figures S1 and S2 for more detail).

**Figure S4.**
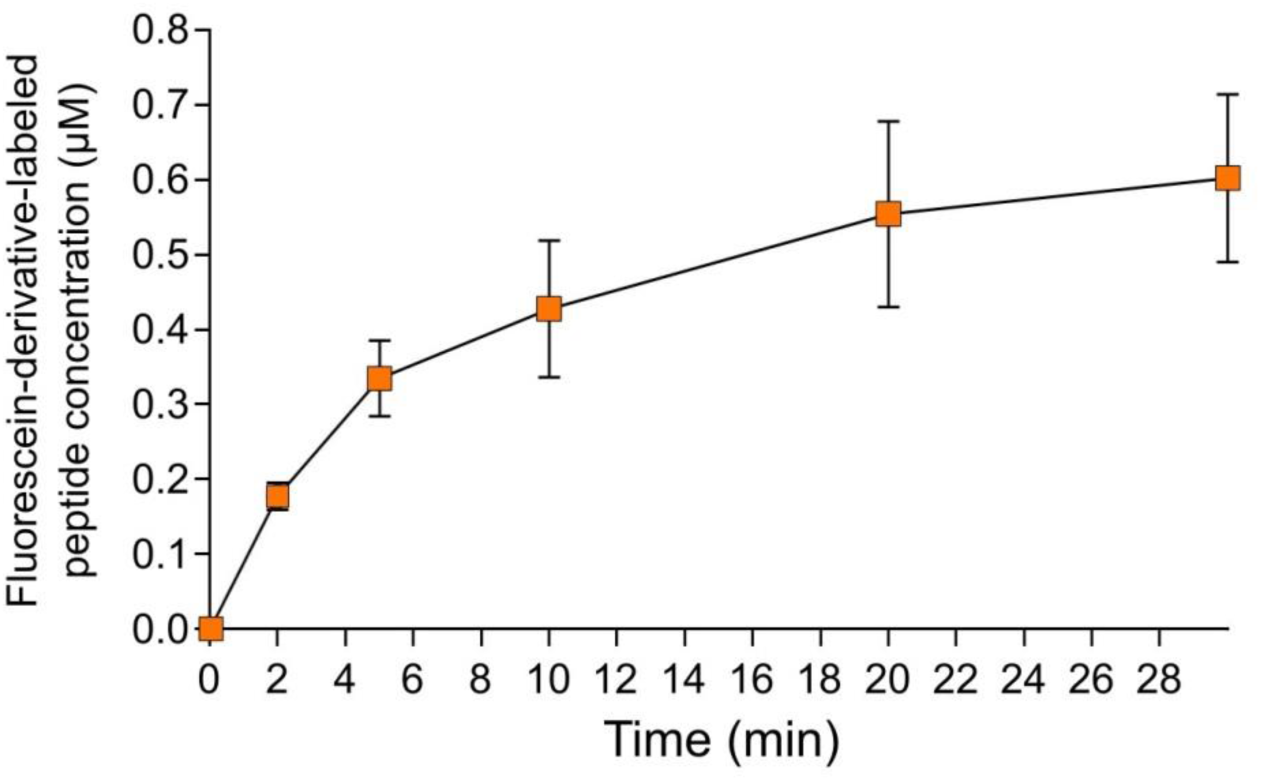
Peptide gradient concentration measurement. Average fluorescein-tagged peptide concentration as a function of time, measured in the center of the experimental glass chamber at 0, 2, 5, 10, 20 and 30 min. Each data point represents the average (± SD) of six measurements (duplicate sampling in three separate experimental replicates) taken at the location where the amoebae were placed. Peptide concentration rises to ∼ 0.2 μM two minutes after the laminar flow is established, and further to 0.6 μM towards the end of the experiment (30 min since laminar flow is established).

**Figure S5:**
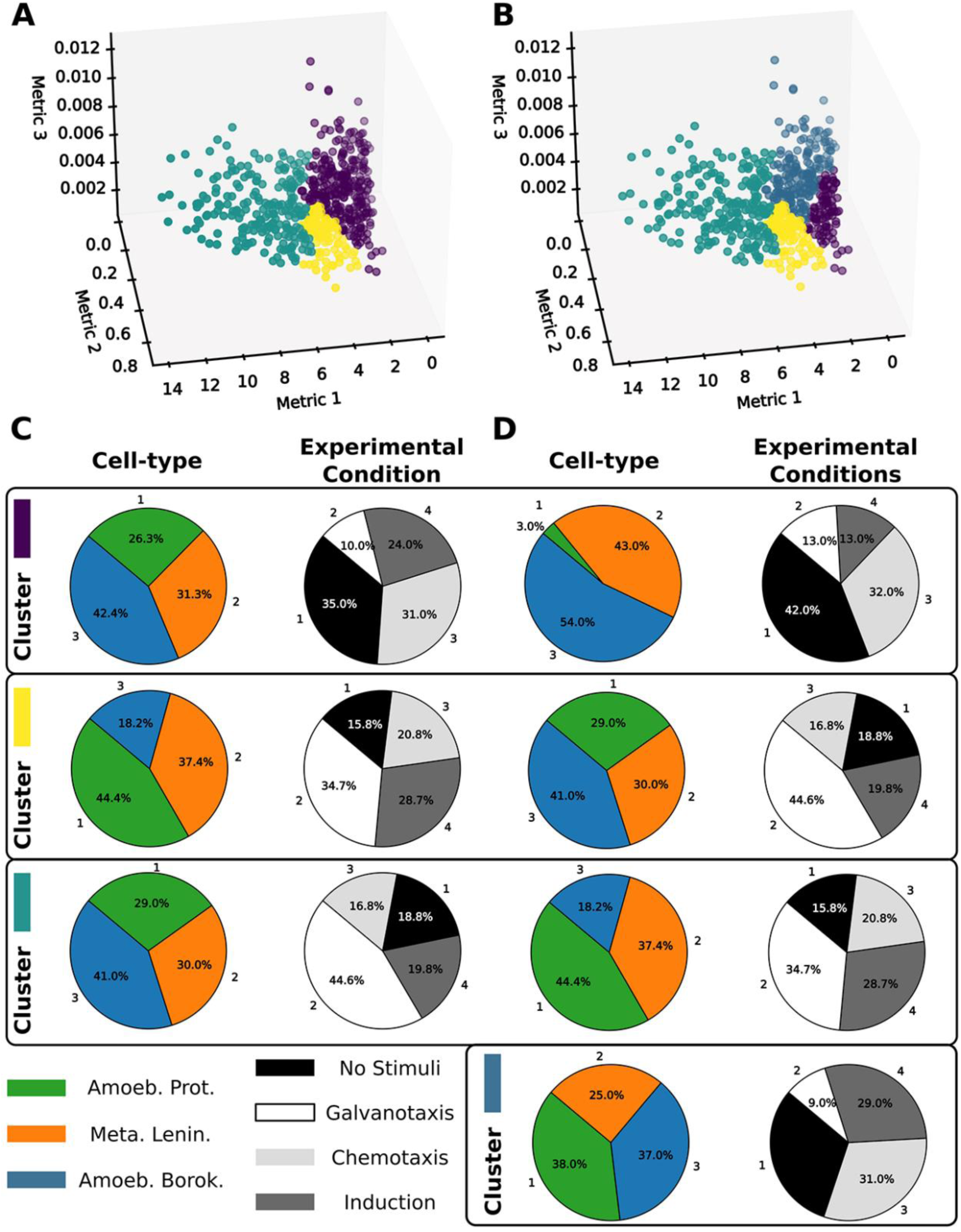
Similar to Fig. 6, but employing a different clustering strategy, namely hierarchical agglomerative clustering. The clustering characterization regarding the distinction between cell-types or experimental conditions remains unchanged when varying the clustering strategy, indicating the robustness of the findings.

**Figure S6:**
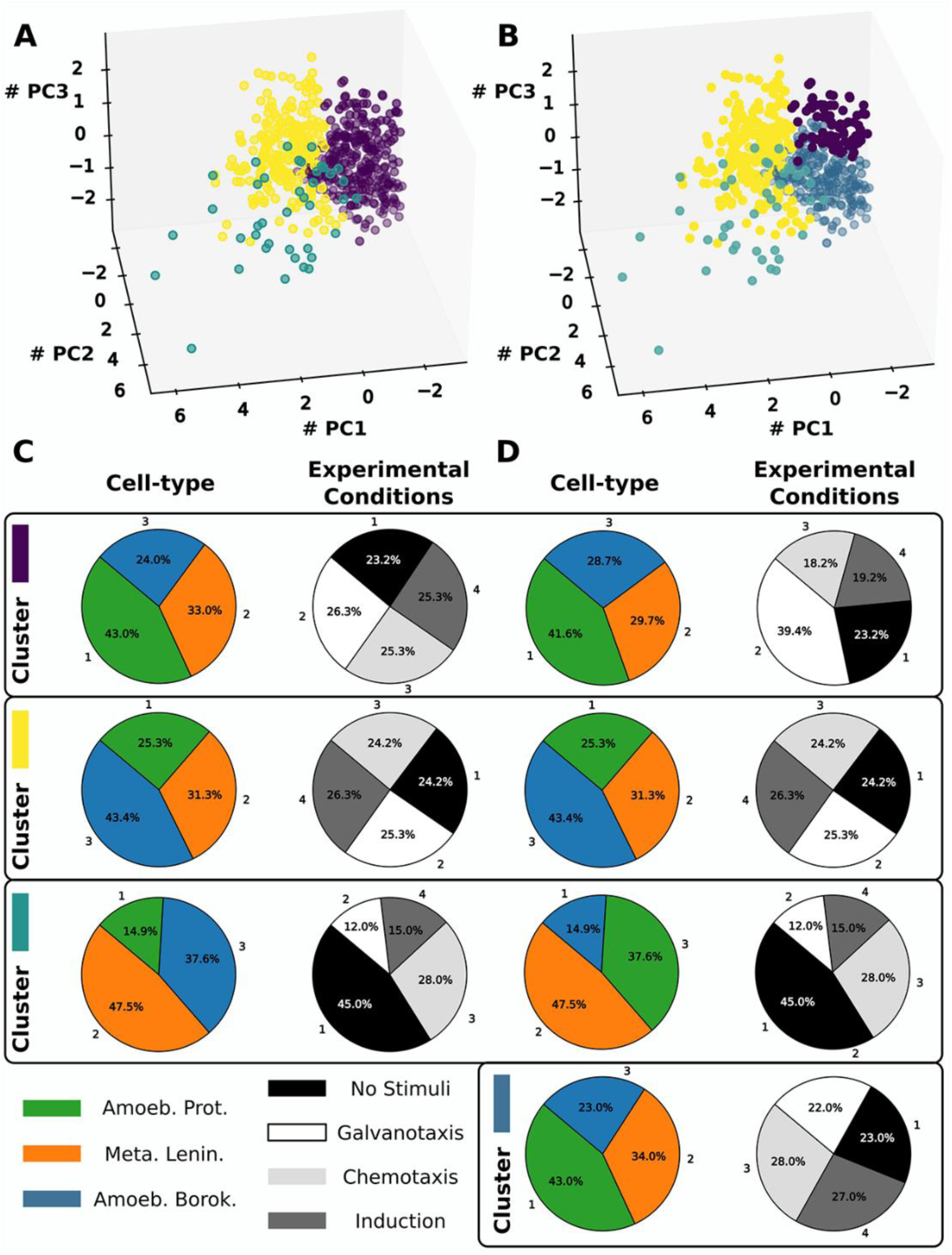
Similar to Fig. 7, but employing a different clustering strategy, namely hierarchical agglomerative clustering.

**Table S1.** Results of “rmsf” analysis (scaling exponent α) of the 700 experimental cell trajectories.

| Cell number | Amoeba proteus |  |  |  | Metamoeba leningradensis |  |  |  | Amoeba borokensis |  |  |  |
| --- | --- | --- | --- | --- | --- | --- | --- | --- | --- | --- | --- | --- |
|  | Sc1 | Sc2 | Sc3 | Sc4 | Sc1 | Sc2 | Sc3 | Sc4 | Sc1 | Sc2 | Sc3 | Sc4 |
| 1 | 0.791 | 0.718 | 0.783 | 0.682 | 0.808 | 0.653 | 0.797 | 0.755 | 0.684 | 0.645 | 0.645 | 0.670 |
| 2 | 0.748 | 0.774 | 0.693 | 0.733 | 0.853 | 0.788 | 0.731 | 0.812 | 0.648 | 0.757 | 0.636 | 0.691 |
| 3 | 0.748 | 0.677 | 0.846 | 0.666 | 0.708 | 0.679 | 0.647 | 0.796 | 0.658 | 0.681 | 0.747 | 0.746 |
| 4 | 0.709 | 0.562 | 0.753 | 0.723 | 0.737 | 0.666 | 0.763 | 0.778 | 0.631 | 0.641 | 0.679 | 0.781 |
| 5 | 0.837 | 0.747 | 0.707 | 0.706 | 0.736 | 0.620 | 0.755 | 0.846 | 0.758 | 0.629 | 0.686 | 0.694 |
| 6 | 0.740 | 0.668 | 0.607 | 0.800 | 0.844 | 0.684 | 0.727 | 0.717 | 0.663 | 0.769 | 0.622 | 0.750 |
| 7 | 0.672 | 0.621 | 0.650 | 0.750 | 0.695 | 0.759 | 0.714 | 0.730 | 0.739 | 0.765 | 0.710 | 0.684 |
| 8 | 0.811 | 0.720 | 0.707 | 0.835 | 0.817 | 0.721 | 0.814 | 0.715 | 0.639 | 0.759 | 0.758 | 0.782 |
| 9 | 0.719 | 0.707 | 0.708 | 0.787 | 0.756 | 0.642 | 0.828 | 0.766 | 0.724 | 0.737 | 0.722 | 0.731 |
| 10 | 0.812 | 0.666 | 0.715 | 0.653 | 0.686 | 0.711 | 0.787 | 0.761 | 0.782 | 0.734 | 0.743 | 0.733 |
| 11 | 0.703 | 0.693 | 0.568 | 0.804 | 0.721 | 0.633 | 0.685 | 0.771 | 0.689 | 0.731 | 0.732 | 0.784 |
| 12 | 0.611 | 0.714 | 0.699 | 0.710 | 0.749 | 0.676 | 0.783 | 0.741 | 0.642 | 0.691 | 0.705 | 0.670 |
| 13 | 0.715 | 0.691 | 0.773 | 0.756 | 0.748 | 0.691 | 0.653 | 0.604 | 0.750 | 0.606 | 0.713 | 0.775 |
| 14 | 0.698 | 0.686 | 0.790 | 0.782 | 0.674 | 0.771 | 0.757 | 0.729 | 0.682 | 0.698 | 0.689 | 0.758 |
| 15 | 0.599 | 0.772 | 0.757 | 0.682 | 0.784 | 0.807 | 0.747 | 0.643 | 0.714 | 0.585 | 0.632 | 0.723 |
| 16 | 0.736 | 0.796 | 0.638 | 0.700 | 0.707 | 0.739 | 0.803 | 0.738 | 0.738 | 0.709 | 0.688 | 0.740 |
| 17 | 0.652 | 0.653 | 0.782 | 0.799 | 0.604 | 0.557 | 0.710 | 0.801 | 0.677 | 0.762 | 0.743 | 0.680 |
| 18 | 0.635 | 0.727 | 0.677 | 0.771 | 0.681 | 0.734 | 0.733 | 0.605 | 0.689 | 0.746 | 0.707 | 0.607 |
| 19 | 0.560 | 0.637 | 0.757 | 0.721 | 0.799 | 0.685 | 0.714 | 0.612 | 0.721 | 0.734 | 0.684 | 0.644 |
| 20 | 0.757 | 0.693 | 0.719 | 0.738 | 0.763 | 0.663 | 0.732 | 0.715 | 0.606 | 0.683 | 0.687 | 0.746 |
| 21 | 0.593 | 0.700 | 0.806 | 0.664 | 0.796 | 0.699 | 0.787 | 0.723 | 0.628 | 0.590 | 0.774 | 0.680 |
| 22 | 0.753 | 0.785 | 0.748 | 0.764 | 0.822 | 0.640 | 0.768 | 0.690 | 0.628 | 0.685 | 0.732 | 0.796 |
| 23 | 0.785 | 0.690 | 0.668 | 0.714 | 0.753 | 0.708 | 0.660 | 0.695 | 0.648 | 0.707 | 0.624 | 0.707 |
| 24 | 0.641 | 0.609 | 0.648 | 0.731 | 0.781 | 0.790 | 0.674 | 0.696 | 0.680 | 0.710 | 0.684 | 0.655 |
| 25 | 0.776 | 0.713 | 0.775 | 0.785 | 0.748 | 0.648 | 0.760 | 0.756 | 0.661 | 0.618 | 0.721 | 0.716 |
| 26 | 0.669 | 0.691 | 0.761 | 0.664 | 0.739 | 0.783 | 0.773 | 0.729 | 0.622 | 0.797 | 0.719 | 0.696 |
| 27 | 0.615 | 0.690 | 0.679 | 0.771 | 0.764 | 0.666 | 0.656 | 0.780 | 0.663 | 0.641 | 0.735 | 0.764 |
| 28 | 0.778 | 0.722 | 0.754 | 0.656 | 0.824 | 0.685 | 0.754 | 0.716 | 0.674 | 0.714 | 0.599 | 0.617 |
| 29 | 0.749 | 0.799 | 0.722 | 0.759 | 0.708 | 0.732 | 0.749 | 0.692 | 0.657 | 0.646 | 0.728 | 0.688 |
| 30 | 0.693 | 0.754 | 0.707 | 0.804 | 0.682 | 0.838 | 0.726 | 0.789 | 0.615 | 0.705 | 0.743 | 0.671 |
| 31 | 0.595 | 0.734 | 0.732 | 0.726 | 0.746 | 0.713 | 0.665 | 0.666 | 0.704 | 0.593 | 0.704 | 0.685 |
| 32 | 0.599 | 0.580 | 0.752 | 0.774 | 0.767 | 0.682 | 0.597 | 0.731 | 0.796 | 0.632 | 0.721 | 0.777 |
| 33 | 0.568 | 0.642 | 0.723 | 0.703 | 0.720 | 0.782 | 0.578 | 0.669 | 0.676 | 0.580 | 0.830 | 0.674 |
| 34 | 0.744 | 0.740 | 0.718 | 0.746 | 0.679 | 0.633 | 0.721 | 0.737 | 0.685 | 0.712 | 0.718 | 0.746 |
| 35 | 0.745 | 0.710 | 0.687 | 0.769 | 0.764 | 0.732 | 0.627 | 0.730 | 0.666 | 0.610 | 0.681 | 0.688 |
| 36 | 0.683 | 0.770 | 0.731 | 0.749 | 0.806 | 0.725 | 0.657 | 0.710 | 0.743 | 0.577 | 0.710 | 0.664 |
| 37 | 0.737 | 0.768 | 0.729 | 0.726 | 0.807 | 0.792 | 0.645 | 0.692 | 0.772 | 0.694 | 0.768 | 0.666 |
| 38 | 0.740 | 0.784 | 0.781 | 0.676 | 0.714 | 0.764 | 0.843 | 0.721 | 0.694 | 0.815 | 0.771 | 0.729 |
| 39 | 0.707 | 0.732 | 0.729 | 0.844 | 0.793 | 0.695 | 0.761 | 0.646 | 0.746 | 0.646 | 0.807 | 0.736 |
| 40 | 0.732 | 0.636 | 0.708 | 0.721 | 0.809 | 0.742 | 0.739 | 0.681 | 0.715 | 0.700 | 0.730 | 0.694 |
| 41 | 0.716 | 0.774 | 0.728 | 0.749 | 0.756 | 0.794 | 0.783 | 0.703 | 0.765 | 0.786 | 0.781 | 0.783 |
| 42 | 0.778 | 0.745 | 0.789 | 0.700 | 0.699 | 0.696 | 0.759 | 0.662 | 0.595 | 0.601 | 0.728 | 0.702 |
| 43 | 0.657 | 0.660 | 0.627 | 0.695 | 0.743 | 0.715 | 0.802 | 0.600 | 0.639 | 0.604 | 0.782 | 0.697 |
| 44 | 0.577 | 0.688 | 0.742 | 0.762 | 0.755 | 0.689 | 0.782 | 0.768 | 0.669 | 0.626 | 0.837 | 0.726 |
| 45 | 0.599 | 0.631 | 0.707 | 0.805 | 0.785 | 0.677 | 0.810 | 0.747 | 0.694 | 0.632 | 0.765 | 0.640 |
| 46 | 0.583 | 0.677 | 0.705 | 0.795 | 0.736 | 0.683 | 0.767 | 0.761 | 0.720 | 0.678 | 0.789 | 0.755 |
| 47 | 0.688 | 0.718 | 0.770 | 0.729 | 0.814 | 0.735 | 0.694 | 0.778 | 0.686 | 0.674 | 0.740 | 0.705 |
| 48 | 0.740 | 0.647 | 0.690 | 0.763 | 0.830 | 0.728 | 0.821 | 0.792 | 0.663 | 0.730 | 0.829 | 0.837 |
| 49 | 0.762 | 0.758 | 0.654 | 0.704 | 0.728 |  | 0.745 | 0.705 | 0.701 | 0.839 | 0.775 | 0.670 |
| 50 | 0.676 |  | 0.741 | 0.725 | 0.845 |  | 0.710 | 0.679 | 0.733 | 0.693 | 0.806 | 0.774 |
| 51 |  |  | 0.745 | 0.678 | 0.722 |  | 0.730 | 0.754 | 0.668 |  | 0.766 | 0.730 |
| 52 |  |  |  | 0.646 |  |  | 0.752 | 0.764 | 0.649 |  | 0.751 | 0.641 |
| 53 |  |  |  | 0.703 |  |  | 0.707 | 0.692 |  |  | 0.786 | 0.723 |
| 54 |  |  |  | 0.609 |  |  | 0.770 | 0.757 |  |  | 0.835 | 0.727 |
| 55 |  |  |  | 0.666 |  |  | 0.683 | 0.848 |  |  | 0.801 | 0.795 |
| 56 |  |  |  | 0.674 |  |  | 0.734 | 0.756 |  |  |  | 0.702 |
| 57 |  |  |  | 0.748 |  |  | 0.685 | 0.828 |  |  |  | 0.752 |
| 58 |  |  |  | 0.794 |  |  | 0.664 | 0.772 |  |  |  | 0.733 |
| 59 |  |  |  | 0.747 |  |  | 0.737 | 0.772 |  |  |  | 0.695 |
| 60 |  |  |  | 0.744 |  |  | 0.630 | 0.654 |  |  |  | 0.728 |
| 61 |  |  |  | 0.810 |  |  |  | 0.722 |  |  |  | 0.684 |
| 62 |  |  |  | 0.799 |  |  |  | 0.692 |  |  |  | 0.687 |
| 63 |  |  |  | 0.671 |  |  |  | 0.746 |  |  |  | 0.678 |
| 64 |  |  |  | 0.867 |  |  |  | 0.806 |  |  |  | 0.768 |
| 65 |  |  |  | 0.783 |  |  |  | 0.775 |  |  |  | 0.758 |
| 66 |  |  |  | 0.664 |  |  |  | 0.747 |  |  |  | 0.732 |
| 67 |  |  |  | 0.713 |  |  |  | 0.691 |  |  |  | 0.810 |
| 68 |  |  |  | 0.713 |  |  |  | 0.689 |  |  |  | 0.737 |
| 69 |  |  |  | 0.760 |  |  |  | 0.690 |  |  |  | 0.707 |
| 70 |  |  |  | 0.647 |  |  |  | 0.812 |  |  |  | 0.776 |
| 71 |  |  |  | 0.735 |  |  |  | 0.827 |  |  |  | 0.794 |
| 72 |  |  |  | 0.823 |  |  |  | 0.766 |  |  |  | 0.700 |
| 73 |  |  |  | 0.663 |  |  |  | 0.802 |  |  |  | 0.751 |
| 74 |  |  |  | 0.733 |  |  |  |  |  |  |  | 0.794 |
| 75 |  |  |  | 0.821 |  |  |  |  |  |  |  | 0.775 |
| 76 |  |  |  | 0.838 |  |  |  |  |  |  |  | 0.840 |
| 77 |  |  |  | 0.808 |  |  |  |  |  |  |  | 0.701 |
| 78 |  |  |  | 0.755 |  |  |  |  |  |  |  | 0.736 |
| 79 |  |  |  | 0.794 |  |  |  |  |  |  |  |  |
| 80 |  |  |  | 0.775 |  |  |  |  |  |  |  |  |
| 81 |  |  |  | 0.727 |  |  |  |  |  |  |  |  |
| 82 |  |  |  | 0.746 |  |  |  |  |  |  |  |  |
| 83 |  |  |  | 0.793 |  |  |  |  |  |  |  |  |

**Table S2.** Results of “rmsf” analysis (scaling exponent α) of the 700 shuffled cell trajectories.

| Cell number | Amoeba proteus |  |  |  | Metamoeba leningradensis |  |  |  | Amoeba borokensis |  |  |  |
| --- | --- | --- | --- | --- | --- | --- | --- | --- | --- | --- | --- | --- |
|  | Sc1 | Sc2 | Sc3 | Sc4 | Sc1 | Sc2 | Sc3 | Sc4 | Sc1 | Sc2 | Sc3 | Sc4 |
| 1 | 0.433 | 0.477 | 0.440 | 0.423 | 0.473 | 0.453 | 0.477 | 0.553 | 0.484 | 0.493 | 0.436 | 0.459 |
| 2 | 0.463 | 0.517 | 0.492 | 0.539 | 0.474 | 0.448 | 0.533 | 0.407 | 0.548 | 0.449 | 0.415 | 0.449 |
| 3 | 0.446 | 0.538 | 0.469 | 0.500 | 0.584 | 0.493 | 0.434 | 0.449 | 0.485 | 0.466 | 0.513 | 0.448 |
| 4 | 0.502 | 0.523 | 0.428 | 0.502 | 0.459 | 0.478 | 0.490 | 0.466 | 0.391 | 0.462 | 0.429 | 0.413 |
| 5 | 0.540 | 0.452 | 0.516 | 0.422 | 0.448 | 0.529 | 0.435 | 0.467 | 0.467 | 0.463 | 0.515 | 0.467 |
| 6 | 0.460 | 0.544 | 0.457 | 0.474 | 0.423 | 0.414 | 0.410 | 0.494 | 0.407 | 0.508 | 0.509 | 0.488 |
| 7 | 0.414 | 0.495 | 0.502 | 0.595 | 0.432 | 0.450 | 0.494 | 0.487 | 0.444 | 0.589 | 0.512 | 0.457 |
| 8 | 0.533 | 0.473 | 0.400 | 0.459 | 0.468 | 0.452 | 0.509 | 0.400 | 0.539 | 0.475 | 0.451 | 0.481 |
| 9 | 0.583 | 0.460 | 0.389 | 0.455 | 0.393 | 0.381 | 0.470 | 0.583 | 0.528 | 0.590 | 0.644 | 0.456 |
| 10 | 0.459 | 0.435 | 0.453 | 0.479 | 0.538 | 0.511 | 0.411 | 0.467 | 0.446 | 0.515 | 0.456 | 0.472 |
| 11 | 0.471 | 0.397 | 0.490 | 0.400 | 0.431 | 0.418 | 0.460 | 0.534 | 0.431 | 0.479 | 0.399 | 0.504 |
| 12 | 0.386 | 0.487 | 0.479 | 0.417 | 0.448 | 0.424 | 0.485 | 0.433 | 0.531 | 0.473 | 0.498 | 0.470 |
| 13 | 0.516 | 0.475 | 0.408 | 0.489 | 0.479 | 0.365 | 0.470 | 0.451 | 0.481 | 0.568 | 0.556 | 0.461 |
| 14 | 0.432 | 0.509 | 0.566 | 0.545 | 0.492 | 0.495 | 0.425 | 0.422 | 0.398 | 0.532 | 0.369 | 0.471 |
| 15 | 0.573 | 0.546 | 0.390 | 0.393 | 0.486 | 0.501 | 0.429 | 0.490 | 0.480 | 0.449 | 0.445 | 0.470 |
| 16 | 0.525 | 0.487 | 0.464 | 0.502 | 0.472 | 0.456 | 0.505 | 0.480 | 0.643 | 0.460 | 0.468 | 0.484 |
| 17 | 0.423 | 0.535 | 0.545 | 0.401 | 0.492 | 0.444 | 0.488 | 0.481 | 0.516 | 0.547 | 0.447 | 0.507 |
| 18 | 0.477 | 0.504 | 0.420 | 0.408 | 0.361 | 0.437 | 0.527 | 0.373 | 0.565 | 0.455 | 0.487 | 0.403 |
| 19 | 0.452 | 0.444 | 0.380 | 0.438 | 0.476 | 0.526 | 0.383 | 0.401 | 0.384 | 0.432 | 0.424 | 0.418 |
| 20 | 0.472 | 0.525 | 0.436 | 0.520 | 0.421 | 0.499 | 0.399 | 0.418 | 0.442 | 0.460 | 0.454 | 0.521 |
| 21 | 0.575 | 0.476 | 0.436 | 0.425 | 0.428 | 0.595 | 0.368 | 0.393 | 0.559 | 0.536 | 0.455 | 0.479 |
| 22 | 0.397 | 0.591 | 0.475 | 0.530 | 0.502 | 0.492 | 0.538 | 0.391 | 0.601 | 0.477 | 0.561 | 0.438 |
| 23 | 0.488 | 0.467 | 0.504 | 0.461 | 0.476 | 0.397 | 0.530 | 0.585 | 0.446 | 0.569 | 0.475 | 0.495 |
| 24 | 0.511 | 0.541 | 0.430 | 0.381 | 0.480 | 0.575 | 0.587 | 0.476 | 0.484 | 0.469 | 0.386 | 0.442 |
| 25 | 0.454 | 0.393 | 0.465 | 0.422 | 0.454 | 0.415 | 0.434 | 0.369 | 0.435 | 0.476 | 0.480 | 0.436 |
| 26 | 0.440 | 0.494 | 0.527 | 0.555 | 0.464 | 0.418 | 0.392 | 0.405 | 0.449 | 0.368 | 0.523 | 0.545 |
| 27 | 0.471 | 0.397 | 0.453 | 0.502 | 0.487 | 0.433 | 0.432 | 0.412 | 0.473 | 0.515 | 0.593 | 0.482 |
| 28 | 0.505 | 0.441 | 0.436 | 0.433 | 0.448 | 0.468 | 0.504 | 0.428 | 0.422 | 0.490 | 0.453 | 0.437 |
| 29 | 0.502 | 0.486 | 0.405 | 0.498 | 0.481 | 0.555 | 0.432 | 0.462 | 0.470 | 0.569 | 0.476 | 0.483 |
| 30 | 0.397 | 0.477 | 0.525 | 0.449 | 0.516 | 0.522 | 0.498 | 0.520 | 0.447 | 0.477 | 0.400 | 0.473 |
| 31 | 0.500 | 0.433 | 0.454 | 0.435 | 0.384 | 0.421 | 0.502 | 0.455 | 0.464 | 0.467 | 0.477 | 0.491 |
| 32 | 0.480 | 0.400 | 0.477 | 0.408 | 0.392 | 0.536 | 0.568 | 0.482 | 0.458 | 0.421 | 0.459 | 0.438 |
| 33 | 0.498 | 0.482 | 0.492 | 0.587 | 0.476 | 0.495 | 0.621 | 0.476 | 0.503 | 0.438 | 0.493 | 0.489 |
| 34 | 0.478 | 0.421 | 0.454 | 0.519 | 0.450 | 0.497 | 0.543 | 0.419 | 0.596 | 0.475 | 0.430 | 0.434 |
| 35 | 0.426 | 0.562 | 0.416 | 0.443 | 0.517 | 0.405 | 0.578 | 0.543 | 0.391 | 0.423 | 0.510 | 0.482 |
| 36 | 0.432 | 0.490 | 0.475 | 0.374 | 0.494 | 0.378 | 0.510 | 0.439 | 0.482 | 0.420 | 0.435 | 0.445 |
| 37 | 0.477 | 0.444 | 0.415 | 0.444 | 0.384 | 0.442 | 0.573 | 0.462 | 0.379 | 0.477 | 0.462 | 0.395 |
| 38 | 0.487 | 0.449 | 0.437 | 0.596 | 0.483 | 0.477 | 0.412 | 0.531 | 0.476 | 0.467 | 0.480 | 0.413 |
| 39 | 0.455 | 0.469 | 0.473 | 0.477 | 0.404 | 0.435 | 0.431 | 0.427 | 0.476 | 0.448 | 0.409 | 0.459 |
| 40 | 0.516 | 0.557 | 0.514 | 0.483 | 0.490 | 0.461 | 0.388 | 0.495 | 0.503 | 0.451 | 0.535 | 0.550 |
| 41 | 0.567 | 0.422 | 0.455 | 0.432 | 0.403 | 0.510 | 0.400 | 0.470 | 0.504 | 0.451 | 0.440 | 0.481 |
| 42 | 0.488 | 0.442 | 0.510 | 0.477 | 0.436 | 0.504 | 0.492 | 0.534 | 0.444 | 0.431 | 0.409 | 0.422 |
| 43 | 0.444 | 0.436 | 0.462 | 0.476 | 0.419 | 0.484 | 0.448 | 0.483 | 0.463 | 0.485 | 0.427 | 0.416 |
| 44 | 0.448 | 0.449 | 0.503 | 0.410 | 0.568 | 0.454 | 0.472 | 0.414 | 0.551 | 0.491 | 0.524 | 0.476 |
| 45 | 0.477 | 0.502 | 0.488 | 0.508 | 0.393 | 0.444 | 0.466 | 0.501 | 0.487 | 0.504 | 0.415 | 0.429 |
| 46 | 0.529 | 0.461 | 0.459 | 0.422 | 0.466 | 0.550 | 0.451 | 0.486 | 0.451 | 0.424 | 0.484 | 0.501 |
| 47 | 0.354 | 0.426 | 0.527 | 0.425 | 0.458 | 0.429 | 0.611 | 0.441 | 0.423 | 0.473 | 0.559 | 0.518 |
| 48 | 0.431 | 0.480 | 0.442 | 0.523 | 0.406 | 0.460 | 0.441 | 0.365 | 0.466 | 0.548 | 0.562 | 0.509 |
| 49 | 0.497 | 0.498 | 0.535 | 0.472 | 0.441 |  | 0.411 | 0.404 | 0.421 | 0.395 | 0.454 | 0.504 |
| 50 | 0.406 |  | 0.558 | 0.448 | 0.469 |  | 0.407 | 0.425 | 0.432 | 0.536 | 0.446 | 0.556 |
| 51 |  |  | 0.494 | 0.479 | 0.508 |  | 0.479 | 0.520 | 0.497 |  | 0.428 | 0.521 |
| 52 |  |  |  | 0.500 |  |  | 0.617 | 0.522 | 0.480 |  | 0.447 | 0.364 |
| 53 |  |  |  | 0.434 |  |  | 0.563 | 0.469 |  |  | 0.552 | 0.417 |
| 54 |  |  |  | 0.470 |  |  | 0.457 | 0.393 |  |  | 0.368 | 0.503 |
| 55 |  |  |  | 0.451 |  |  | 0.483 | 0.414 |  |  | 0.390 | 0.523 |
| 56 |  |  |  | 0.409 |  |  | 0.453 | 0.465 |  |  |  | 0.490 |
| 57 |  |  |  | 0.527 |  |  | 0.481 | 0.443 |  |  |  | 0.449 |
| 58 |  |  |  | 0.568 |  |  | 0.520 | 0.481 |  |  |  | 0.411 |
| 59 |  |  |  | 0.427 |  |  | 0.404 | 0.408 |  |  |  | 0.483 |
| 60 |  |  |  | 0.476 |  |  | 0.520 | 0.443 |  |  |  | 0.427 |
| 61 |  |  |  | 0.466 |  |  |  | 0.429 |  |  |  | 0.380 |
| 62 |  |  |  | 0.493 |  |  |  | 0.419 |  |  |  | 0.520 |
| 63 |  |  |  | 0.558 |  |  |  | 0.376 |  |  |  | 0.432 |
| 64 |  |  |  | 0.565 |  |  |  | 0.351 |  |  |  | 0.470 |
| 65 |  |  |  | 0.571 |  |  |  | 0.526 |  |  |  | 0.490 |
| 66 |  |  |  | 0.408 |  |  |  | 0.426 |  |  |  | 0.512 |
| 67 |  |  |  | 0.451 |  |  |  | 0.507 |  |  |  | 0.490 |
| 68 |  |  |  | 0.525 |  |  |  | 0.456 |  |  |  | 0.499 |
| 69 |  |  |  | 0.528 |  |  |  | 0.404 |  |  |  | 0.552 |
| 70 |  |  |  | 0.446 |  |  |  | 0.430 |  |  |  | 0.431 |
| 71 |  |  |  | 0.435 |  |  |  | 0.576 |  |  |  | 0.379 |
| 72 |  |  |  | 0.411 |  |  |  | 0.447 |  |  |  | 0.402 |
| 73 |  |  |  | 0.411 |  |  |  | 0.473 |  |  |  | 0.450 |
| 74 |  |  |  | 0.520 |  |  |  |  |  |  |  | 0.515 |
| 75 |  |  |  | 0.446 |  |  |  |  |  |  |  | 0.438 |
| 76 |  |  |  | 0.534 |  |  |  |  |  |  |  | 0.456 |
| 77 |  |  |  | 0.447 |  |  |  |  |  |  |  | 0.516 |
| 78 |  |  |  | 0.549 |  |  |  |  |  |  |  | 0.446 |
| 79 |  |  |  | 0.388 |  |  |  |  |  |  |  |  |
| 80 |  |  |  | 0.457 |  |  |  |  |  |  |  |  |
| 81 |  |  |  | 0.436 |  |  |  |  |  |  |  |  |
| 82 |  |  |  | 0.535 |  |  |  |  |  |  |  |  |
| 83 |  |  |  | 0.613 |  |  |  |  |  |  |  |  |

**Table S3.** Long-range correlation duration values for the 700 experimental cell trajectories.

| Cell number | Amoeba proteus |  |  |  | Metamoeba leningradensis |  |  |  | Amoeba borokensis |  |  |  |
| --- | --- | --- | --- | --- | --- | --- | --- | --- | --- | --- | --- | --- |
|  | Sc1 | Sc2 | Sc3 | Sc4 | Sc1 | Sc2 | Sc3 | Sc4 | Sc1 | Sc2 | Sc3 | Sc4 |
| 1 | 2.083 | 14.583 | 11.458 | 10.417 | 13.542 | 15.625 | 15.625 | 4.167 | 14.583 | 10.417 | 14.583 | 5.208 |
| 2 | 13.542 | 15.625 | 6.250 | 5.208 | 12.500 | 7.292 | 7.292 | 10.417 | 12.500 | 8.333 | 8.333 | 5.208 |
| 3 | 14.583 | 11.458 | 14.583 | 4.167 | 5.208 | 7.292 | 16.667 | 14.583 | 9.375 | 11.458 | 10.417 | 4.167 |
| 4 | 8.333 | 7.292 | 8.333 | 3.125 | 7.292 | 12.500 | 5.208 | 9.375 | 6.250 | 3.125 | 13.542 | 2.083 |
| 5 | 9.375 | 15.625 | 5.208 | 14.583 | 6.250 | 3.125 | 13.542 | 13.542 | 14.583 | 4.167 | 2.083 | 11.458 |
| 6 | 10.417 | 15.625 | 11.458 | 5.208 | 11.458 | 4.167 | 8.333 | 10.417 | 4.167 | 14.583 | 4.167 | 3.125 |
| 7 | 7.292 | 16.667 | 15.625 | 4.167 | 4.167 | 15.625 | 6.250 | 13.542 | 13.542 | 3.125 | 10.417 | 11.458 |
| 8 | 12.500 | 15.625 | 9.375 | 13.542 | 13.542 | 6.250 | 6.250 | 6.250 | 10.417 | 4.167 | 7.292 | 3.125 |
| 9 | 5.208 | 8.333 | 3.125 | 7.292 | 4.167 | 5.208 | 6.250 | 8.333 | 9.375 | 10.417 | 8.333 | 4.167 |
| 10 | 5.208 | 16.667 | 8.333 | 14.583 | 8.333 | 7.292 | 12.500 | 8.333 | 15.625 | 13.542 | 6.250 | 6.250 |
| 11 | 3.125 | 13.542 | 8.333 | 14.583 | 11.458 | 14.583 | 6.250 | 6.250 | 7.292 | 15.625 | 7.292 | 7.292 |
| 12 | 13.542 | 13.542 | 10.417 | 12.500 | 5.208 | 7.292 | 14.583 | 6.250 | 14.583 | 6.250 | 15.625 | 3.125 |
| 13 | 5.208 | 5.208 | 11.458 | 12.500 | 4.167 | 7.292 | 11.458 | 7.292 | 13.542 | 6.250 | 6.250 | 4.167 |
| 14 | 5.208 | 12.500 | 8.333 | 14.583 | 14.583 | 7.292 | 3.125 | 6.250 | 15.625 | 10.417 | 4.167 | 4.167 |
| 15 | 13.542 | 10.417 | 6.250 | 10.417 | 14.583 | 11.458 | 13.542 | 3.125 | 2.083 | 8.333 | 13.542 | 4.167 |
| 16 | 2.083 | 10.417 | 12.500 | 7.292 | 8.333 | 10.417 | 2.083 | 5.208 | 12.500 | 3.125 | 6.250 | 9.375 |
| 17 | 14.583 | 7.292 | 9.375 | 12.500 | 10.417 | 15.625 | 4.167 | 5.208 | 3.125 | 7.292 | 8.333 | 7.292 |
| 18 | 3.125 | 7.292 | 13.542 | 3.125 | 12.500 | 5.208 | 3.125 | 10.417 | 2.083 | 4.167 | 12.500 | 11.458 |
| 19 | 9.375 | 15.625 | 12.500 | 11.458 | 11.458 | 13.542 | 4.167 | 14.583 | 6.250 | 9.375 | 11.458 | 14.583 |
| 20 | 13.542 | 14.583 | 9.375 | 3.125 | 5.208 | 13.542 | 5.208 | 4.167 | 8.333 | 12.500 | 7.292 | 12.500 |
| 21 | 12.500 | 2.083 | 11.458 | 11.458 | 7.292 | 11.458 | 15.625 | 6.250 | 6.250 | 2.083 | 7.292 | 14.583 |
| 22 | 15.625 | 11.458 | 7.292 | 10.417 | 10.417 | 13.542 | 3.125 | 14.583 | 6.250 | 12.500 | 6.250 | 3.125 |
| 23 | 13.542 | 2.083 | 5.208 | 10.417 | 12.500 | 7.292 | 3.125 | 13.542 | 3.125 | 4.167 | 12.500 | 9.375 |
| 24 | 15.625 | 2.083 | 4.167 | 8.333 | 10.417 | 15.625 | 13.542 | 6.250 | 4.167 | 5.208 | 5.208 | 5.208 |
| 25 | 10.417 | 9.375 | 10.417 | 13.542 | 2.083 | 14.583 | 11.458 | 7.292 | 5.208 | 13.542 | 8.333 | 10.417 |
| 26 | 6.250 | 5.208 | 10.417 | 16.667 | 8.333 | 9.375 | 5.208 | 6.250 | 14.583 | 14.583 | 5.208 | 8.333 |
| 27 | 14.583 | 14.583 | 9.375 | 4.167 | 13.542 | 8.333 | 13.542 | 14.583 | 9.375 | 2.083 | 15.625 | 15.625 |
| 28 | 14.583 | 1.042 | 13.542 | 11.458 | 13.542 | 9.375 | 8.333 | 13.542 | 2.083 | 11.458 | 10.417 | 9.375 |
| 29 | 14.583 | 11.458 | 14.583 | 12.500 | 15.625 | 10.417 | 5.208 | 6.250 | 7.292 | 7.292 | 6.250 | 8.333 |
| 30 | 10.417 | 12.500 | 5.208 | 13.542 | 16.667 | 14.583 | 6.250 | 10.417 | 8.333 | 8.333 | 7.292 | 5.208 |
| 31 | 7.292 | 13.542 | 7.292 | 13.542 | 9.375 | 10.417 | 12.500 | 12.500 | 4.167 | 12.500 | 8.333 | 7.292 |
| 32 | 3.125 | 15.625 | 12.500 | 13.542 | 12.500 | 8.333 | 14.583 | 12.500 | 15.625 | 4.167 | 4.167 | 3.125 |
| 33 | 14.583 | 11.458 | 12.500 | 10.417 | 15.625 | 7.292 | 9.375 | 10.417 | 2.083 | 5.208 | 10.417 | 11.458 |
| 34 | 13.542 | 15.625 | 6.250 | 12.500 | 15.625 | 7.292 | 15.625 | 12.500 | 6.250 | 14.583 | 8.333 | 9.375 |
| 35 | 9.375 | 6.250 | 9.375 | 10.417 | 8.333 | 2.083 | 4.167 | 5.208 | 11.458 | 16.667 | 6.250 | 13.542 |
| 36 | 8.333 | 13.542 | 8.333 | 9.375 | 15.625 | 11.458 | 12.500 | 3.125 | 14.583 | 11.458 | 12.500 | 12.500 |
| 37 | 9.375 | 8.333 | 11.458 | 13.542 | 9.375 | 15.625 | 4.167 | 6.250 | 5.208 | 9.375 | 7.292 | 5.208 |
| 38 | 9.375 | 15.625 | 10.417 | 10.417 | 8.333 | 11.458 | 4.167 | 5.208 | 14.583 | 14.583 | 13.542 | 10.417 |
| 39 | 9.375 | 13.542 | 11.458 | 10.417 | 9.375 | 14.583 | 12.500 | 10.417 | 6.250 | 13.542 | 11.458 | 10.417 |
| 40 | 7.292 | 11.458 | 7.292 | 10.417 | 10.417 | 6.250 | 13.542 | 10.417 | 3.125 | 3.125 | 7.292 | 5.208 |
| 41 | 15.625 | 15.625 | 9.375 | 9.375 | 6.250 | 13.542 | 7.292 | 6.250 | 11.458 | 8.333 | 4.167 | 10.417 |
| 42 | 6.250 | 2.083 | 8.333 | 4.167 | 3.125 | 13.542 | 11.458 | 4.167 | 7.292 | 8.333 | 6.250 | 8.333 |
| 43 | 15.625 | 4.167 | 11.458 | 2.083 | 5.208 | 6.250 | 9.375 | 5.208 | 6.250 | 7.292 | 12.500 | 3.125 |
| 44 | 15.625 | 5.208 | 7.292 | 6.250 | 8.333 | 6.250 | 13.542 | 3.125 | 9.375 | 7.292 | 15.625 | 13.542 |
| 45 | 10.417 | 6.250 | 15.625 | 6.250 | 9.375 | 4.167 | 14.583 | 12.500 | 5.208 | 7.292 | 7.292 | 8.333 |
| 46 | 2.083 | 15.625 | 11.458 | 9.375 | 10.417 | 11.458 | 10.417 | 14.583 | 2.083 | 3.125 | 10.417 | 4.167 |
| 47 | 7.292 | 13.542 | 5.208 | 10.417 | 7.292 | 12.500 | 7.292 | 14.583 | 15.625 | 4.167 | 5.208 | 10.417 |
| 48 | 10.417 | 15.625 | 12.500 | 12.500 | 14.583 | 9.375 | 15.625 | 13.542 | 13.542 | 7.292 | 11.458 | 3.125 |
| 49 | 14.583 | 15.625 | 14.583 | 7.292 | 6.250 |  | 15.625 | 15.625 | 14.583 | 12.500 | 15.625 | 3.125 |
| 50 | 10.417 |  | 12.500 | 10.417 | 15.625 |  | 8.333 | 13.542 | 11.458 | 6.250 | 8.333 | 3.125 |
| 51 |  |  | 4.167 | 3.125 | 15.625 |  | 6.250 | 15.625 | 7.292 |  | 11.458 | 12.500 |
| 52 |  |  |  | 9.375 |  |  | 14.583 | 11.458 | 10.417 |  | 12.500 | 6.250 |
| 53 |  |  |  | 12.500 |  |  | 10.417 | 13.542 |  |  | 3.125 | 8.333 |
| 54 |  |  |  | 15.625 |  |  | 7.292 | 6.250 |  |  | 14.583 | 8.333 |
| 55 |  |  |  | 11.458 |  |  | 6.250 | 15.625 |  |  | 11.458 | 9.375 |
| 56 |  |  |  | 5.208 |  |  | 8.333 | 15.625 |  |  |  | 3.125 |
| 57 |  |  |  | 11.458 |  |  | 7.292 | 14.583 |  |  |  | 12.500 |
| 58 |  |  |  | 13.542 |  |  | 7.292 | 15.625 |  |  |  | 2.083 |
| 59 |  |  |  | 10.417 |  |  | 3.125 | 15.625 |  |  |  | 7.292 |
| 60 |  |  |  | 10.417 |  |  | 15.625 | 10.417 |  |  |  | 5.208 |
| 61 |  |  |  | 10.417 |  |  |  | 14.583 |  |  |  | 7.292 |
| 62 |  |  |  | 11.458 |  |  |  | 11.458 |  |  |  | 7.292 |
| 63 |  |  |  | 3.125 |  |  |  | 13.542 |  |  |  | 3.125 |
| 64 |  |  |  | 14.583 |  |  |  | 12.500 |  |  |  | 3.125 |
| 65 |  |  |  | 11.458 |  |  |  | 7.292 |  |  |  | 9.375 |
| 66 |  |  |  | 11.458 |  |  |  | 6.250 |  |  |  | 10.417 |
| 67 |  |  |  | 4.167 |  |  |  | 6.250 |  |  |  | 11.458 |
| 68 |  |  |  | 14.583 |  |  |  | 12.500 |  |  |  | 7.292 |
| 69 |  |  |  | 9.375 |  |  |  | 6.250 |  |  |  | 12.500 |
| 70 |  |  |  | 15.625 |  |  |  | 13.542 |  |  |  | 11.458 |
| 71 |  |  |  | 8.333 |  |  |  | 5.208 |  |  |  | 5.208 |
| 72 |  |  |  | 11.458 |  |  |  | 4.167 |  |  |  | 14.583 |
| 73 |  |  |  | 13.542 |  |  |  | 5.208 |  |  |  | 4.167 |
| 74 |  |  |  | 15.625 |  |  |  |  |  |  |  | 7.292 |
| 75 |  |  |  | 14.583 |  |  |  |  |  |  |  | 4.167 |
| 76 |  |  |  | 15.625 |  |  |  |  |  |  |  | 14.583 |
| 77 |  |  |  | 15.625 |  |  |  |  |  |  |  | 4.167 |
| 78 |  |  |  | 8.333 |  |  |  |  |  |  |  | 12.500 |
| 79 |  |  |  | 6.250 |  |  |  |  |  |  |  |  |
| 80 |  |  |  | 4.167 |  |  |  |  |  |  |  |  |
| 81 |  |  |  | 13.542 |  |  |  |  |  |  |  |  |
| 82 |  |  |  | 11.458 |  |  |  |  |  |  |  |  |
| 83 |  |  |  | 14.583 |  |  |  |  |  |  |  |  |

**Table S4.** Results of MSD analysis of the 700 experimental cell trajectories.

| Cell number | Amoeba proteus |  |  |  | Metamoeba leningradensis |  |  |  | Amoeba borokensis |  |  |  |
| --- | --- | --- | --- | --- | --- | --- | --- | --- | --- | --- | --- | --- |
|  | Sc1 | Sc2 | Sc3 | Sc4 | Sc1 | Sc2 | Sc3 | Sc4 | Sc1 | Sc2 | Sc3 | Sc4 |
| 1 | 1.655 | 1.962 | 1.693 | 1.850 | 1.926 | 2.007 | 1.895 | 1.833 | 1.833 | 1.671 | 1.790 | 1.792 |
| 2 | 1.871 | 1.903 | 1.809 | 1.666 | 1.893 | 1.963 | 1.789 | 1.943 | 1.778 | 1.839 | 1.738 | 1.905 |
| 3 | 1.761 | 1.956 | 1.928 | 1.692 | 1.993 | 1.691 | 1.332 | 1.940 | 1.836 | 1.666 | 1.854 | 1.774 |
| 4 | 1.894 | 1.964 | 1.967 | 1.815 | 1.610 | 1.424 | 1.525 | 1.991 | 1.804 | 1.786 | 1.844 | 1.564 |
| 5 | 1.817 | 1.912 | 1.974 | 1.986 | 1.867 | 1.919 | 1.851 | 1.954 | 1.877 | 1.589 | 1.623 | 1.843 |
| 6 | 1.901 | 1.948 | 1.930 | 1.870 | 1.886 | 1.509 | 1.866 | 1.930 | 1.852 | 1.790 | 1.655 | 1.732 |
| 7 | 1.949 | 1.939 | 1.613 | 1.709 | 1.705 | 1.923 | 1.823 | 1.967 | 1.882 | 1.743 | 1.862 | 1.597 |
| 8 | 1.907 | 1.940 | 1.976 | 1.942 | 1.986 | 1.957 | 1.992 | 1.981 | 1.880 | 1.805 | 1.904 | 1.802 |
| 9 | 1.766 | 1.800 | 1.869 | 1.756 | 1.943 | 1.901 | 2.006 | 1.967 | 1.841 | 1.683 | 1.770 | 1.775 |
| 10 | 1.494 | 1.964 | 1.915 | 1.945 | 1.753 | 1.918 | 1.932 | 1.830 | 1.833 | 1.926 | 1.711 | 1.870 |
| 11 | 1.782 | 1.874 | 1.900 | 1.938 | 1.865 | 1.735 | 1.809 | 1.954 | 1.838 | 1.920 | 1.871 | 1.760 |
| 12 | 1.864 | 1.670 | 1.949 | 1.865 | 1.816 | 1.918 | 1.910 | 1.900 | 1.594 | 1.769 | 1.726 | 1.886 |
| 13 | 1.759 | 1.919 | 1.897 | 1.982 | 1.839 | 1.976 | 1.827 | 1.920 | 1.846 | 1.929 | 1.811 | 1.828 |
| 14 | 1.709 | 1.961 | 1.840 | 1.964 | 1.796 | 1.877 | 1.863 | 1.879 | 1.668 | 1.946 | 1.869 | 1.695 |
| 15 | 1.920 | 1.825 | 1.966 | 1.970 | 1.954 | 1.687 | 1.915 | 1.929 | 1.671 | 1.705 | 1.871 | 1.847 |
| 16 | 1.741 | 1.921 | 1.881 | 1.820 | 1.797 | 1.989 | 1.945 | 1.959 | 1.942 | 1.883 | 1.844 | 1.449 |
| 17 | 1.833 | 2.003 | 1.796 | 1.977 | 1.786 | 1.664 | 1.694 | 1.943 | 1.737 | 1.853 | 1.680 | 1.818 |
| 18 | 1.651 | 1.860 | 1.874 | 1.881 | 1.707 | 1.918 | 1.623 | 1.811 | 1.729 | 1.821 | 1.803 | 1.901 |
| 19 | 1.664 | 1.957 | 1.926 | 1.862 | 1.883 | 1.969 | 1.692 | 1.964 | 1.919 | 1.772 | 1.874 | 1.869 |
| 20 | 1.909 | 1.986 | 1.896 | 1.846 | 1.905 | 1.929 | 1.623 | 1.953 | 1.930 | 1.915 | 1.834 | 1.901 |
| 21 | 1.774 | 1.925 | 1.856 | 1.917 | 1.126 | 1.982 | 1.693 | 1.950 | 1.684 | 1.939 | 1.691 | 1.881 |
| 22 | 1.869 | 1.922 | 1.844 | 1.802 | 1.896 | 1.786 | 1.743 | 1.892 | 1.629 | 1.945 | 1.924 | 1.774 |
| 23 | 1.953 | 1.918 | 1.880 | 1.902 | 1.699 | 1.904 | 1.571 | 1.896 | 1.913 | 1.967 | 1.864 | 1.730 |
| 24 | 1.944 | 1.973 | 1.849 | 1.822 | 1.736 | 1.980 | 1.847 | 1.923 | 1.435 | 1.941 | 1.927 | 1.795 |
| 25 | 1.866 | 1.878 | 1.903 | 1.918 | 1.963 | 1.915 | 1.965 | 1.971 | 1.682 | 1.935 | 1.772 | 1.810 |
| 26 | 1.608 | 1.938 | 1.782 | 1.852 | 1.820 | 1.772 | 1.876 | 1.931 | 1.919 | 1.736 | 1.932 | 1.786 |
| 27 | 1.966 | 1.811 | 1.891 | 1.884 | 1.713 | 1.896 | 1.784 | 1.934 | 1.725 | 1.936 | 1.873 | 1.835 |
| 28 | 1.899 | 1.965 | 1.965 | 1.760 | 1.838 | 1.974 | 1.898 | 1.859 | 1.489 | 1.634 | 1.397 | 1.528 |
| 29 | 1.909 | 1.960 | 1.970 | 1.958 | 1.309 | 1.955 | 1.606 | 1.997 | 1.625 | 1.896 | 1.894 | 1.823 |
| 30 | 1.973 | 1.929 | 1.691 | 1.951 | 1.580 | 1.990 | 1.704 | 1.858 | 1.836 | 1.856 | 1.580 | 1.908 |
| 31 | 1.968 | 1.426 | 1.706 | 1.813 | 1.785 | 1.827 | 1.933 | 1.799 | 1.928 | 1.812 | 1.795 | 1.954 |
| 32 | 1.959 | 1.693 | 1.983 | 1.845 | 1.841 | 1.914 | 1.490 | 1.981 | 1.894 | 1.857 | 1.976 | 1.911 |
| 33 | 1.858 | 2.000 | 2.010 | 1.368 | 1.360 | 1.947 | 1.451 | 1.948 | 1.945 | 1.908 | 1.947 | 1.705 |
| 34 | 1.887 | 1.973 | 1.938 | 1.875 | 1.648 | 1.858 | 1.425 | 1.971 | 1.826 | 1.837 | 1.981 | 1.936 |
| 35 | 1.897 | 1.778 | 1.911 | 1.946 | 1.424 | 1.901 | 1.345 | 1.915 | 1.893 | 1.944 | 1.607 | 1.957 |
| 36 | 1.946 | 1.821 | 1.987 | 1.841 | 1.950 | 1.845 | 1.780 | 1.961 | 1.928 | 1.924 | 1.902 | 1.863 |
| 37 | 1.906 | 1.829 | 1.978 | 1.899 | 1.642 | 1.870 | 1.194 | 1.944 | 1.892 | 1.870 | 1.641 | 1.696 |
| 38 | 1.990 | 1.954 | 1.913 | 1.893 | 1.707 | 1.892 | 1.767 | 1.987 | 2.006 | 1.918 | 1.967 | 1.608 |
| 39 | 1.807 | 1.961 | 1.943 | 1.939 | 1.627 | 1.922 | 1.804 | 1.979 | 1.947 | 1.891 | 1.848 | 1.720 |
| 40 | 1.948 | 1.969 | 1.826 | 1.902 | 1.474 | 1.918 | 1.764 | 1.903 | 1.805 | 1.834 | 1.739 | 1.773 |
| 41 | 1.894 | 1.925 | 1.936 | 1.831 | 1.560 | 1.986 | 1.799 | 1.926 | 1.834 | 1.915 | 1.928 | 1.712 |
| 42 | 1.925 | 1.877 | 1.916 | 1.756 | 1.618 | 1.895 | 1.785 | 1.989 | 1.904 | 1.928 | 1.781 | 1.783 |
| 43 | 1.944 | 1.992 | 1.923 | 1.794 | 1.770 | 1.858 | 1.711 | 1.962 | 1.708 | 1.974 | 1.763 | 1.629 |
| 44 | 1.516 | 1.899 | 1.785 | 1.889 | 1.853 | 1.820 | 1.856 | 1.905 | 1.810 | 1.882 | 1.933 | 1.803 |
| 45 | 1.979 | 1.968 | 1.245 | 1.821 | 1.926 | 1.885 | 1.921 | 1.827 | 1.706 | 1.957 | 1.977 | 1.577 |
| 46 | 1.806 | 1.949 | 1.795 | 1.979 | 1.882 | 1.718 | 1.841 | 1.944 | 1.092 | 1.923 | 1.993 | 1.630 |
| 47 | 1.887 | 1.971 | 1.784 | 1.965 | 1.695 | 1.792 | 1.787 | 1.940 | 1.943 | 1.946 | 1.999 | 1.820 |
| 48 | 1.891 | 1.983 | 1.890 | 1.962 | 1.821 | 1.962 | 1.956 | 1.923 | 1.823 | 1.912 | 1.964 | 1.688 |
| 49 | 1.950 | 1.973 | 1.833 | 1.617 | 1.951 |  | 1.872 | 1.810 | 1.895 | 1.901 | 1.960 | 1.825 |
| 50 | 1.876 |  | 1.860 | 1.939 | 1.936 |  | 1.869 | 1.571 | 1.951 | 1.910 | 1.823 | 1.838 |
| 51 |  |  | 1.792 | 1.829 | 1.810 |  | 1.642 | 1.880 | 1.931 |  | 1.942 | 1.845 |
| 52 |  |  |  | 1.752 |  |  | 1.868 | 1.907 | 1.797 |  | 1.837 | 1.979 |
| 53 |  |  |  | 1.857 |  |  | 1.963 | 1.856 |  |  | 1.763 | 1.965 |
| 54 |  |  |  | 1.566 |  |  | 1.936 | 1.916 |  |  | 1.970 | 1.863 |
| 55 |  |  |  | 1.748 |  |  | 1.775 | 1.959 |  |  | 1.861 | 1.950 |
| 56 |  |  |  | 1.687 |  |  | 1.765 | 1.898 |  |  |  | 1.868 |
| 57 |  |  |  | 1.935 |  |  | 1.694 | 1.888 |  |  |  | 1.909 |
| 58 |  |  |  | 1.890 |  |  | 1.645 | 1.957 |  |  |  | 1.842 |
| 59 |  |  |  | 1.934 |  |  | 1.364 | 1.896 |  |  |  | 2.002 |
| 60 |  |  |  | 1.600 |  |  | 1.431 | 1.729 |  |  |  | 1.846 |
| 61 |  |  |  | 1.864 |  |  |  | 1.904 |  |  |  | 1.551 |
| 62 |  |  |  | 1.884 |  |  |  | 1.874 |  |  |  | 1.903 |
| 63 |  |  |  | 1.935 |  |  |  | 1.725 |  |  |  | 1.693 |
| 64 |  |  |  | 1.923 |  |  |  | 1.938 |  |  |  | 1.612 |
| 65 |  |  |  | 1.940 |  |  |  | 1.978 |  |  |  | 1.770 |
| 66 |  |  |  | 1.950 |  |  |  | 1.840 |  |  |  | 1.717 |
| 67 |  |  |  | 1.675 |  |  |  | 1.441 |  |  |  | 1.862 |
| 68 |  |  |  | 1.865 |  |  |  | 1.749 |  |  |  | 1.939 |
| 69 |  |  |  | 1.916 |  |  |  | 1.976 |  |  |  | 1.995 |
| 70 |  |  |  | 1.723 |  |  |  | 1.891 |  |  |  | 2.022 |
| 71 |  |  |  | 1.907 |  |  |  | 1.837 |  |  |  | 1.890 |
| 72 |  |  |  | 1.972 |  |  |  | 1.878 |  |  |  | 1.785 |
| 73 |  |  |  | 1.778 |  |  |  | 1.709 |  |  |  | 1.936 |
| 74 |  |  |  | 1.929 |  |  |  |  |  |  |  | 1.925 |
| 75 |  |  |  | 1.811 |  |  |  |  |  |  |  | 1.870 |
| 76 |  |  |  | 1.916 |  |  |  |  |  |  |  | 1.907 |
| 77 |  |  |  | 1.968 |  |  |  |  |  |  |  | 1.787 |
| 78 |  |  |  | 1.980 |  |  |  |  |  |  |  | 1.902 |
| 79 |  |  |  | 1.958 |  |  |  |  |  |  |  |  |
| 80 |  |  |  | 1.849 |  |  |  |  |  |  |  |  |
| 81 |  |  |  | 1.860 |  |  |  |  |  |  |  |  |
| 82 |  |  |  | 1.870 |  |  |  |  |  |  |  |  |
| 83 |  |  |  | 1.949 |  |  |  |  |  |  |  |  |

**Table S5.**
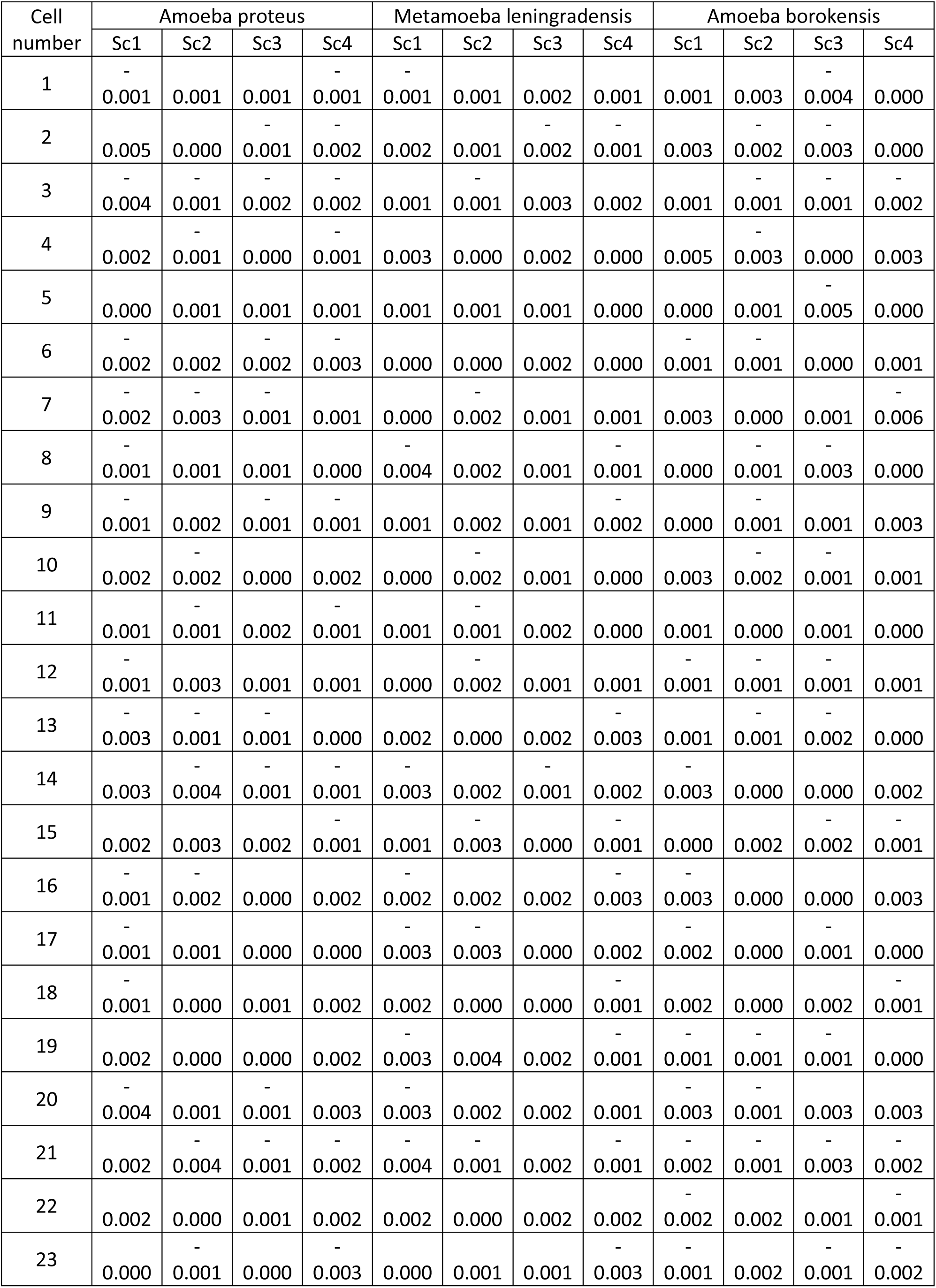
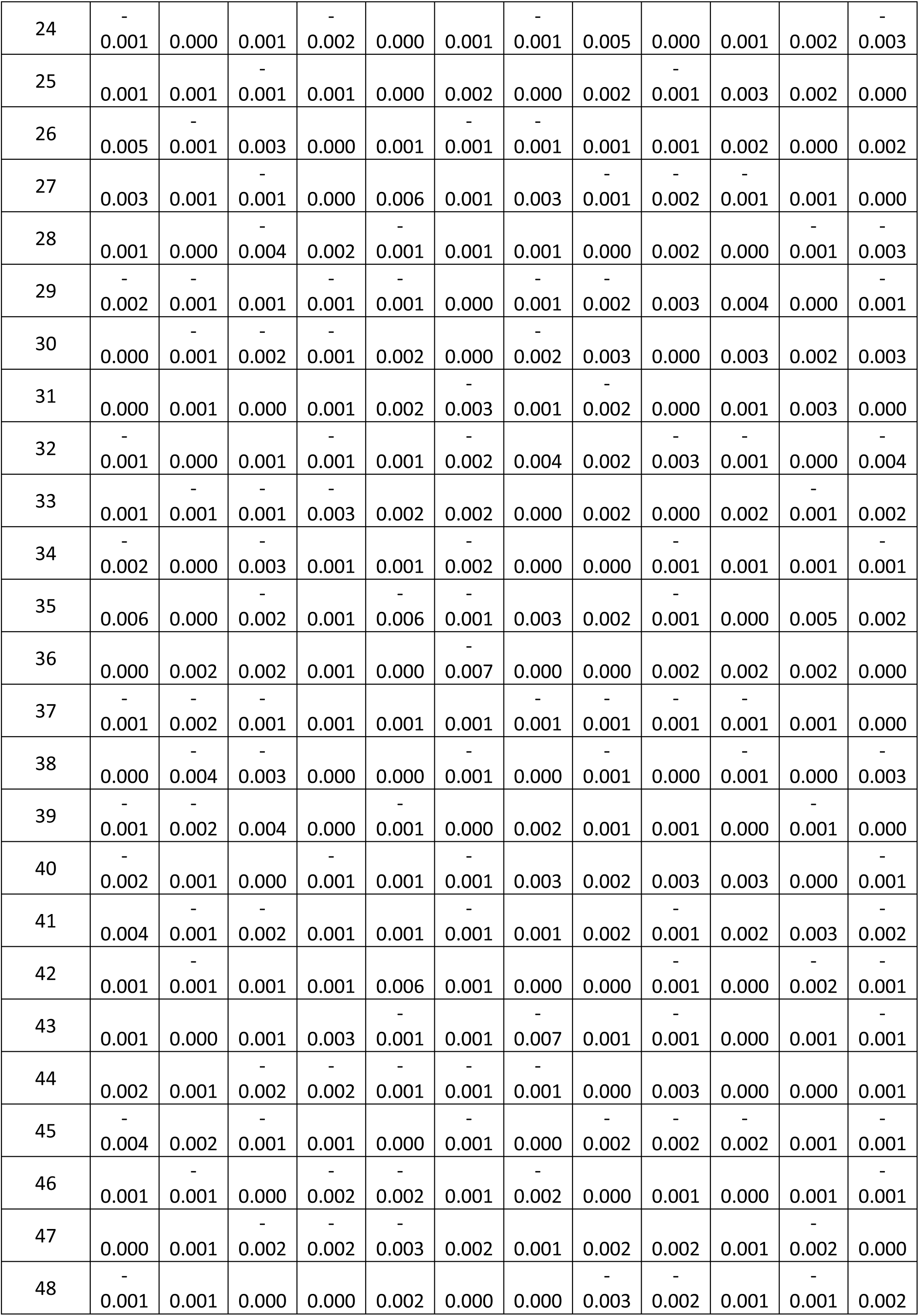
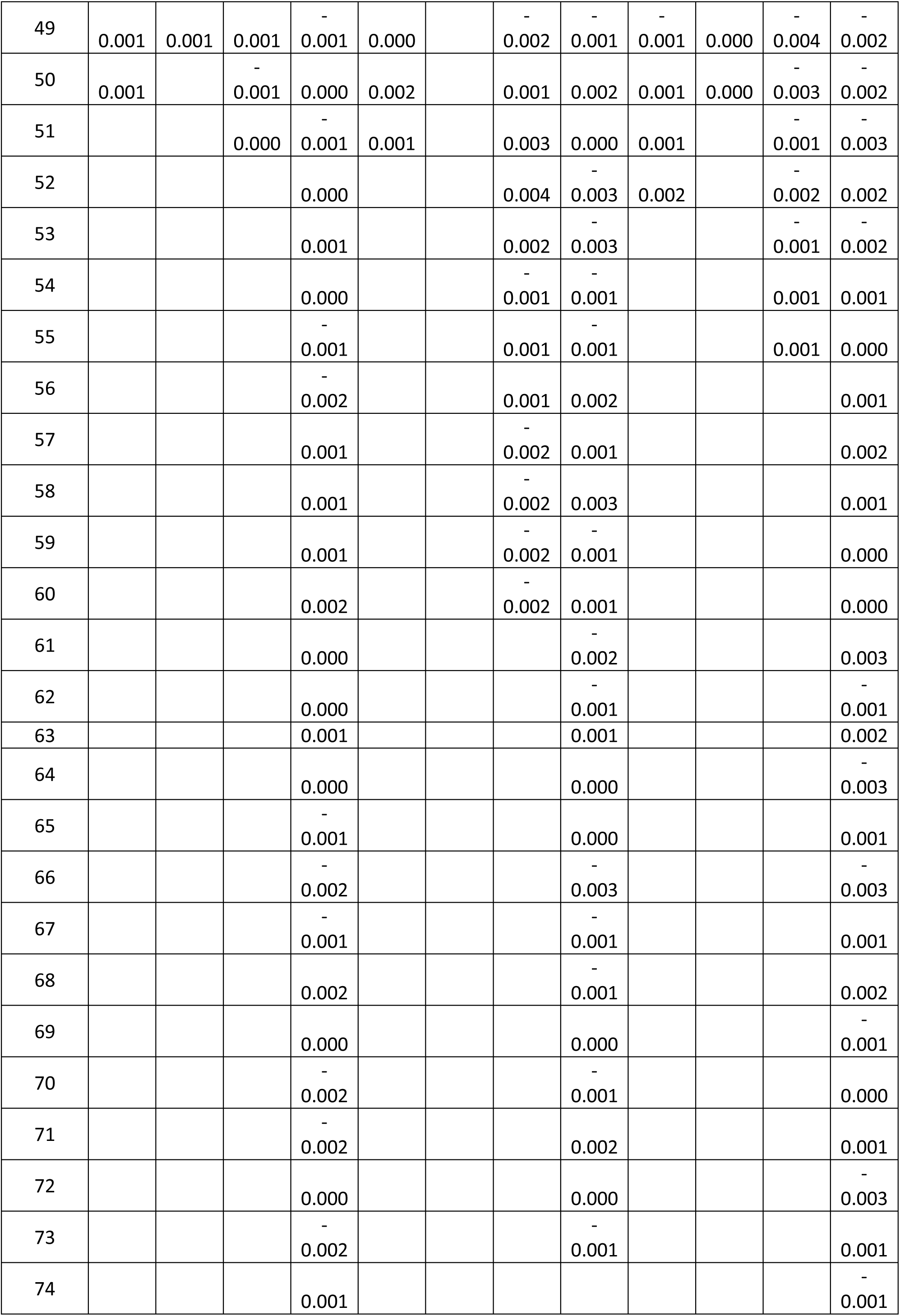
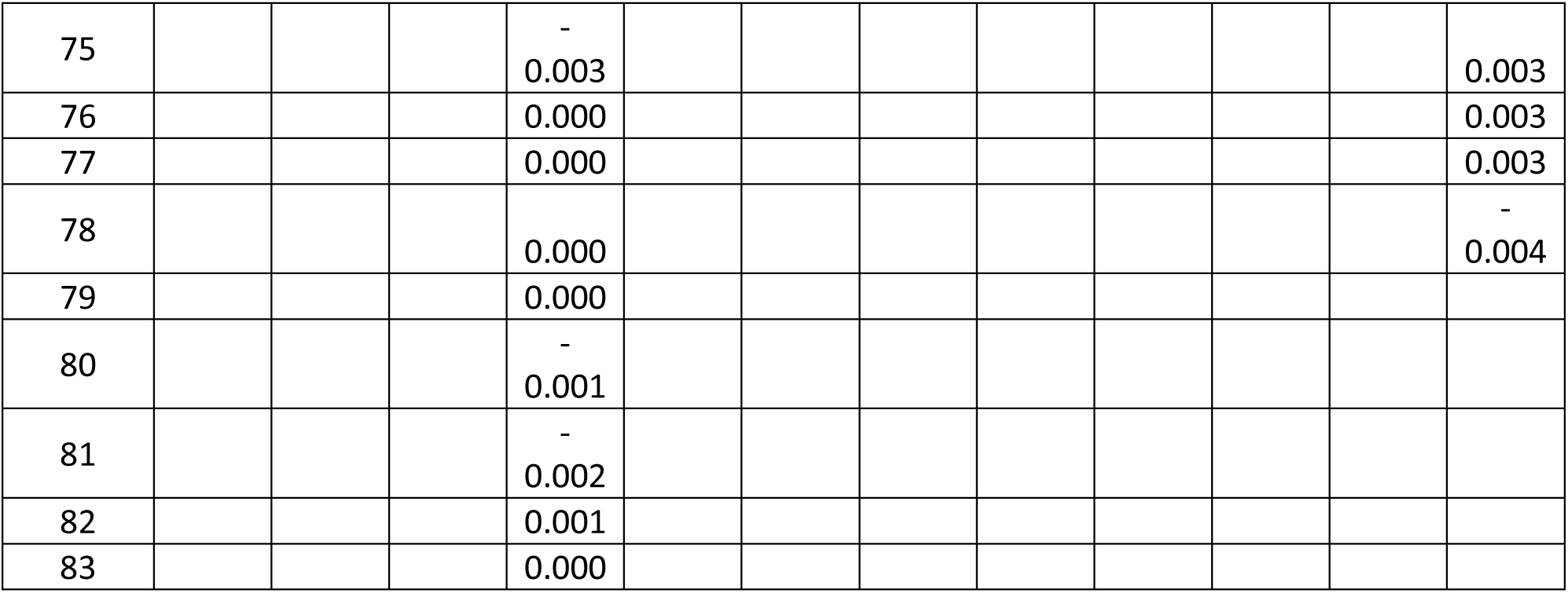
Results of MSD analysis of the 700 shuffled cell trajectories.

**Table S6.**
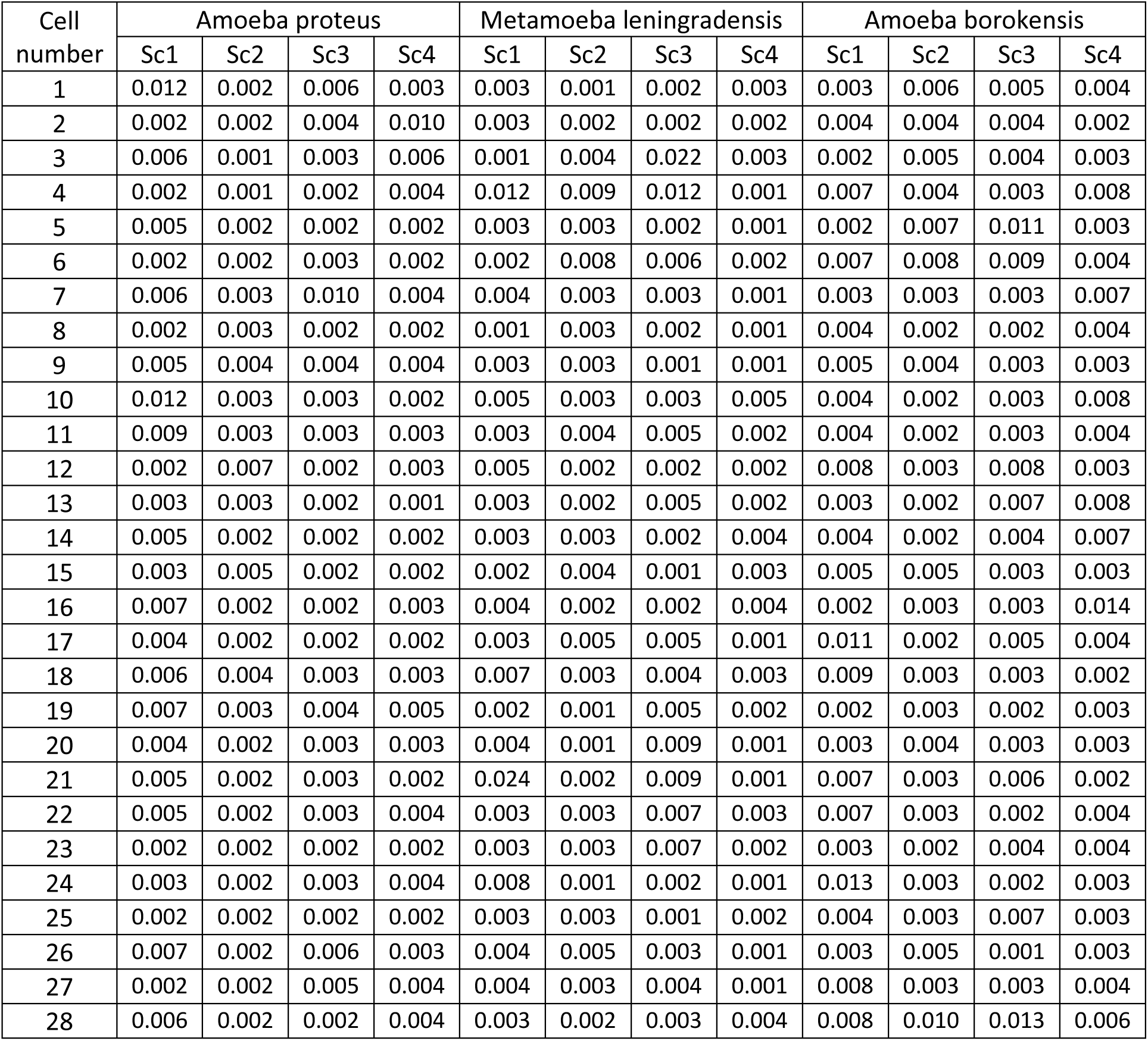
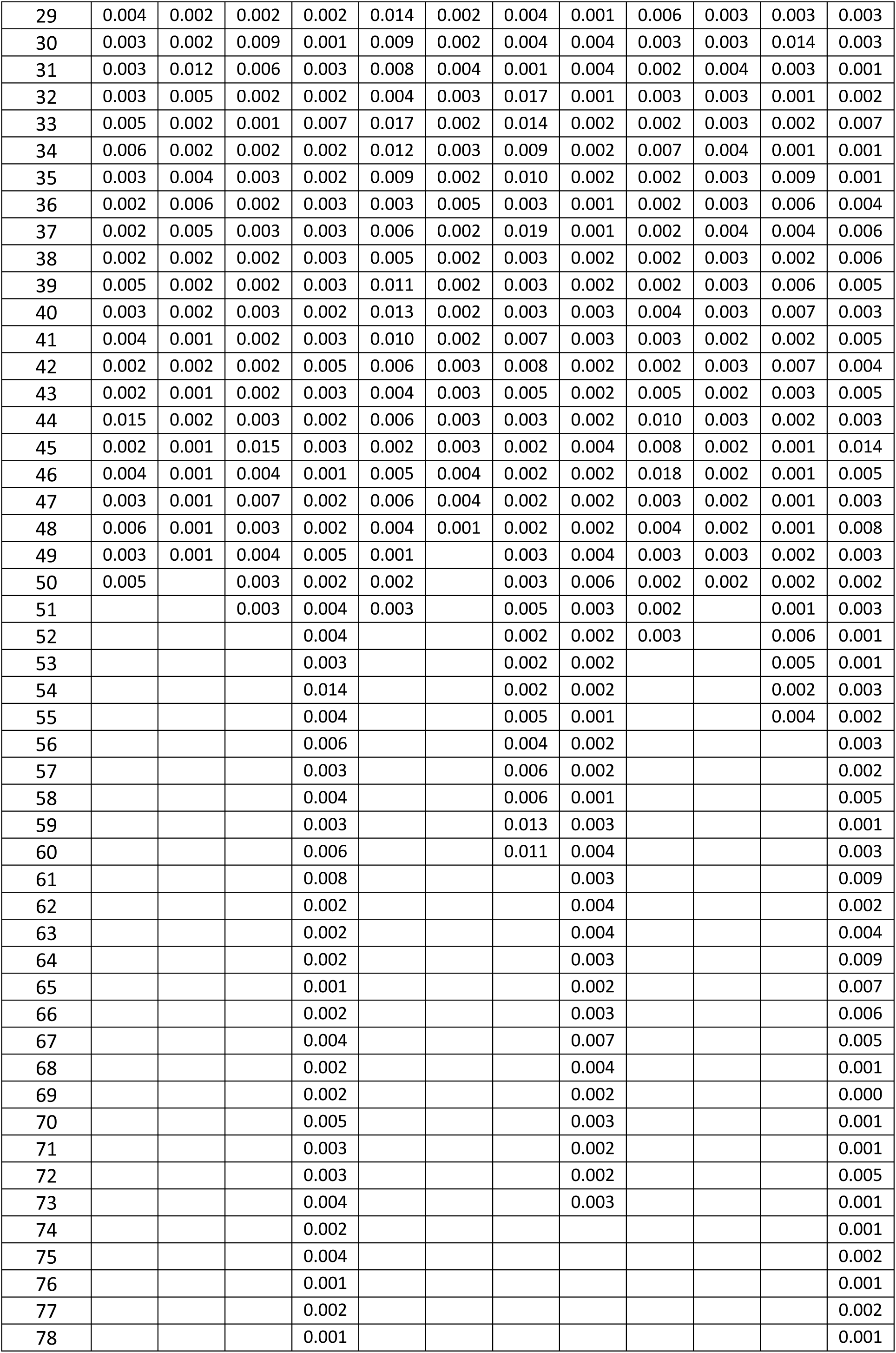
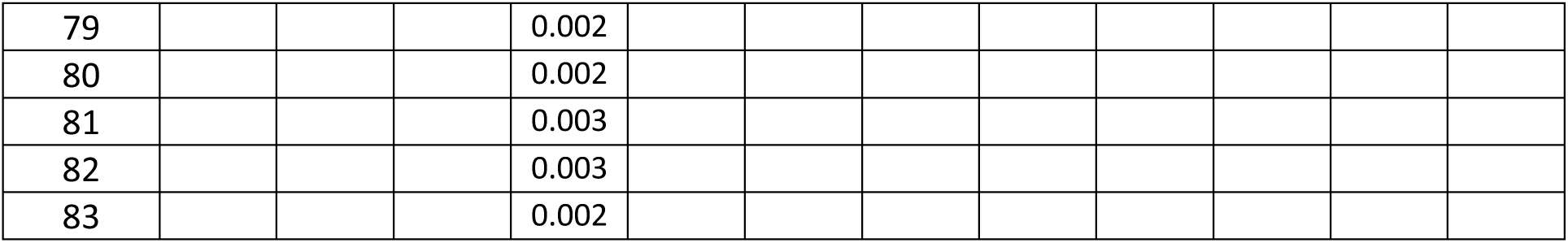
Results of Approximate Entropy estimation for the 700 experimental cell trajectories.

**Table S7.** Results of Approximate Entropy estimation for the 700 shuffled cell trajectories.

| Cell number | Amoeba proteus |  |  |  | Metamoeba leningradensis |  |  |  | Amoeba borokensis |  |  |  |
| --- | --- | --- | --- | --- | --- | --- | --- | --- | --- | --- | --- | --- |
|  | Sc1 | Sc2 | Sc3 | Sc4 | Sc1 | Sc2 | Sc3 | Sc4 | Sc1 | Sc2 | Sc3 | Sc4 |
| 1 | 2.071 | 1.964 | 1.853 | 1.857 | 1.968 | 2.067 | 1.613 | 1.927 | 1.688 | 1.896 | 1.933 | 2.045 |
| 2 | 1.849 | 1.901 | 2.004 | 1.952 | 1.762 | 1.892 | 1.963 | 2.062 | 1.922 | 1.879 | 1.840 | 2.049 |
| 3 | 1.740 | 2.097 | 1.683 | 1.826 | 2.090 | 1.974 | 2.006 | 1.901 | 1.890 | 1.669 | 1.605 | 1.970 |
| 4 | 2.009 | 2.077 | 1.947 | 1.897 | 1.830 | 1.894 | 2.000 | 1.886 | 1.988 | 2.033 | 1.871 | 1.995 |
| 5 | 1.892 | 2.031 | 2.079 | 2.015 | 1.914 | 2.079 | 1.770 | 1.656 | 1.747 | 1.960 | 2.057 | 1.761 |
| 6 | 1.979 | 1.943 | 1.953 | 1.928 | 1.574 | 1.992 | 1.982 | 1.864 | 2.041 | 1.788 | 1.836 | 1.957 |
| 7 | 2.053 | 1.986 | 2.076 | 2.015 | 1.960 | 2.004 | 1.998 | 2.016 | 1.979 | 1.807 | 1.681 | 1.813 |
| 8 | 1.700 | 2.036 | 2.028 | 1.650 | 2.007 | 2.023 | 1.991 | 2.070 | 1.908 | 1.996 | 1.806 | 2.076 |
| 9 | 1.938 | 1.856 | 1.993 | 1.763 | 1.960 | 2.074 | 1.916 | 2.003 | 1.941 | 1.911 | 1.752 | 1.993 |
| 10 | 2.003 | 2.033 | 2.032 | 1.991 | 1.996 | 2.021 | 1.823 | 1.989 | 1.608 | 1.917 | 1.789 | 2.055 |
| 11 | 2.006 | 2.034 | 2.115 | 1.762 | 1.832 | 1.931 | 1.977 | 1.962 | 1.891 | 1.984 | 1.826 | 1.859 |
| 12 | 1.931 | 1.732 | 2.027 | 1.814 | 1.958 | 1.999 | 2.003 | 1.878 | 1.734 | 2.008 | 1.660 | 2.088 |
| 13 | 1.936 | 1.969 | 1.955 | 2.046 | 1.996 | 2.069 | 1.984 | 1.904 | 2.010 | 2.102 | 1.997 | 2.003 |
| 14 | 2.028 | 2.074 | 2.008 | 1.973 | 1.636 | 2.022 | 2.024 | 1.606 | 2.026 | 2.099 | 1.881 | 1.976 |
| 15 | 2.093 | 1.815 | 2.026 | 1.976 | 2.085 | 1.595 | 2.008 | 2.079 | 1.895 | 1.807 | 2.090 | 1.830 |
| 16 | 2.013 | 2.036 | 2.050 | 1.833 | 1.699 | 2.049 | 2.065 | 2.019 | 1.891 | 1.950 | 1.945 | 1.928 |
| 17 | 2.083 | 2.046 | 1.992 | 1.849 | 2.023 | 1.978 | 1.918 | 2.059 | 2.031 | 1.983 | 1.867 | 2.013 |
| 18 | 1.865 | 1.922 | 1.957 | 1.984 | 1.894 | 2.080 | 1.981 | 1.972 | 2.032 | 2.037 | 1.888 | 2.079 |
| 19 | 1.809 | 2.043 | 1.960 | 1.745 | 1.893 | 2.081 | 1.953 | 2.096 | 2.039 | 1.960 | 1.944 | 1.942 |
| 20 | 1.866 | 2.036 | 1.973 | 1.963 | 1.981 | 2.108 | 2.039 | 2.108 | 2.086 | 1.912 | 2.005 | 2.018 |
| 21 | 1.885 | 2.104 | 1.851 | 2.037 | 1.766 | 2.041 | 1.969 | 2.082 | 1.859 | 2.093 | 1.804 | 1.925 |
| 22 | 1.857 | 1.745 | 1.945 | 1.734 | 1.809 | 1.865 | 2.049 | 2.081 | 1.912 | 2.097 | 2.032 | 1.920 |
| 23 | 1.912 | 2.097 | 1.960 | 2.075 | 1.252 | 2.106 | 1.821 | 1.971 | 2.051 | 2.042 | 1.956 | 1.968 |
| 24 | 1.901 | 2.098 | 1.862 | 1.862 | 1.954 | 2.051 | 1.919 | 2.065 | 1.879 | 2.014 | 2.076 | 2.066 |
| 25 | 1.923 | 2.053 | 1.895 | 1.826 | 2.055 | 2.062 | 1.982 | 1.953 | 1.897 | 2.101 | 1.999 | 1.870 |
| 26 | 1.969 | 2.115 | 1.747 | 2.053 | 1.976 | 1.852 | 1.999 | 2.072 | 1.952 | 1.818 | 2.106 | 1.929 |
| 27 | 2.081 | 1.834 | 1.991 | 1.989 | 1.428 | 2.088 | 1.856 | 2.052 | 1.869 | 2.070 | 1.821 | 1.613 |
| 28 | 2.052 | 2.066 | 2.007 | 1.862 | 1.357 | 2.039 | 1.877 | 1.985 | 1.974 | 1.898 | 1.972 | 2.053 |
| 29 | 1.918 | 1.888 | 2.022 | 2.001 | 1.728 | 1.969 | 1.924 | 2.081 | 1.797 | 2.085 | 1.956 | 1.979 |
| 30 | 2.001 | 2.006 | 1.899 | 1.855 | 1.605 | 2.014 | 1.759 | 1.960 | 1.858 | 2.066 | 1.896 | 2.037 |
| 31 | 2.051 | 1.589 | 1.803 | 1.860 | 1.881 | 2.018 | 1.970 | 1.830 | 1.965 | 1.973 | 1.995 | 2.048 |
| 32 | 2.074 | 1.922 | 1.899 | 1.941 | 1.885 | 2.030 | 1.999 | 2.055 | 1.653 | 2.062 | 2.055 | 2.054 |
| 33 | 2.037 | 2.036 | 1.990 | 1.671 | 1.835 | 2.007 | 2.015 | 1.963 | 2.077 | 2.100 | 1.800 | 1.952 |
| 34 | 1.947 | 2.066 | 2.088 | 1.904 | 1.988 | 2.098 | 1.942 | 1.901 | 2.006 | 1.860 | 2.021 | 2.041 |
| 35 | 2.039 | 1.961 | 1.890 | 1.879 | 1.579 | 2.072 | 1.890 | 1.957 | 1.903 | 2.091 | 1.738 | 1.997 |
| 36 | 2.085 | 1.956 | 2.013 | 1.898 | 1.749 | 1.897 | 1.819 | 2.094 | 1.916 | 2.054 | 2.081 | 1.980 |
| 37 | 2.080 | 2.034 | 1.872 | 1.896 | 1.754 | 1.697 | 2.016 | 2.075 | 1.964 | 2.088 | 1.773 | 1.707 |
| 38 | 2.013 | 2.054 | 1.894 | 2.080 | 1.844 | 1.990 | 1.782 | 2.003 | 2.016 | 1.951 | 1.913 | 1.786 |
| 39 | 1.827 | 2.028 | 1.969 | 1.677 | 1.919 | 2.078 | 1.811 | 2.036 | 1.981 | 2.079 | 1.920 | 1.901 |
| 40 | 2.029 | 2.087 | 1.910 | 1.826 | 1.945 | 1.975 | 1.543 | 1.999 | 1.834 | 2.057 | 2.011 | 1.968 |
| 41 | 1.843 | 2.077 | 1.986 | 1.780 | 1.890 | 1.980 | 1.912 | 1.830 | 1.977 | 2.077 | 1.879 | 1.778 |
| 42 | 1.927 | 2.092 | 1.822 | 1.934 | 1.610 | 2.018 | 1.998 | 2.026 | 2.126 | 1.921 | 2.059 | 1.896 |
| 43 | 2.053 | 2.064 | 2.043 | 2.049 | 1.714 | 1.963 | 1.835 | 2.008 | 2.059 | 2.068 | 1.749 | 1.928 |
| 44 | 1.952 | 2.057 | 1.862 | 1.810 | 1.935 | 2.012 | 1.651 | 2.001 | 2.082 | 2.072 | 1.711 | 2.032 |
| 45 | 2.075 | 2.085 | 1.967 | 1.899 | 1.884 | 2.017 | 1.763 | 1.699 | 1.953 | 2.089 | 2.075 | 1.841 |
| 46 | 1.931 | 2.075 | 1.999 | 1.967 | 1.921 | 1.916 | 1.824 | 1.915 | 1.995 | 2.065 | 1.968 | 2.011 |
| 47 | 2.102 | 2.053 | 1.809 | 1.909 | 1.743 | 1.927 | 1.994 | 1.942 | 1.884 | 2.026 | 2.061 | 2.093 |
| 48 | 2.007 | 2.019 | 1.791 | 1.857 | 1.514 | 2.050 | 1.904 | 1.916 | 1.986 | 1.848 | 1.968 | 1.999 |
| 49 | 2.011 | 2.050 | 2.060 | 1.971 | 2.055 |  | 1.820 | 1.934 | 1.967 | 1.880 | 1.950 | 1.986 |
| 50 | 1.831 |  | 1.870 | 1.824 | 1.628 |  | 1.896 | 1.762 | 1.893 | 2.070 | 1.672 | 1.919 |
| 51 |  |  | 1.833 | 1.974 | 1.913 |  | 1.799 | 1.984 | 2.044 |  | 2.045 | 1.821 |
| 52 |  |  |  | 1.873 |  |  | 1.934 | 1.996 | 2.033 |  | 1.809 | 2.045 |
| 53 |  |  |  | 1.689 |  |  | 2.059 | 2.004 |  |  | 1.906 | 2.015 |
| 54 |  |  |  | 1.940 |  |  | 1.805 | 2.059 |  |  | 1.671 | 1.929 |
| 55 |  |  |  | 1.772 |  |  | 1.905 | 1.925 |  |  | 1.963 | 1.831 |
| 56 |  |  |  | 1.963 |  |  | 1.796 | 2.009 |  |  |  | 1.985 |
| 57 |  |  |  | 2.060 |  |  | 1.780 | 1.811 |  |  |  | 1.943 |
| 58 |  |  |  | 1.799 |  |  | 1.630 | 2.038 |  |  |  | 2.016 |
| 59 |  |  |  | 1.992 |  |  | 2.046 | 1.950 |  |  |  | 2.040 |
| 60 |  |  |  | 2.003 |  |  | 1.943 | 1.979 |  |  |  | 1.922 |
| 61 |  |  |  | 1.921 |  |  |  | 1.904 |  |  |  | 1.860 |
| 62 |  |  |  | 1.993 |  |  |  | 1.958 |  |  |  | 2.058 |
| 63 |  |  |  | 2.092 |  |  |  | 1.832 |  |  |  | 1.973 |
| 64 |  |  |  | 1.606 |  |  |  | 1.922 |  |  |  | 1.894 |
| 65 |  |  |  | 1.924 |  |  |  | 1.956 |  |  |  | 2.013 |
| 66 |  |  |  | 2.003 |  |  |  | 1.982 |  |  |  | 1.925 |
| 67 |  |  |  | 1.849 |  |  |  | 2.057 |  |  |  | 1.941 |
| 68 |  |  |  | 1.816 |  |  |  | 1.937 |  |  |  | 1.987 |
| 69 |  |  |  | 1.974 |  |  |  | 1.998 |  |  |  | 2.084 |
| 70 |  |  |  | 1.908 |  |  |  | 1.840 |  |  |  | 2.009 |
| 71 |  |  |  | 1.993 |  |  |  | 1.979 |  |  |  | 2.082 |
| 72 |  |  |  | 1.731 |  |  |  | 2.083 |  |  |  | 1.898 |
| 73 |  |  |  | 1.694 |  |  |  | 1.962 |  |  |  | 2.088 |
| 74 |  |  |  | 1.973 |  |  |  |  |  |  |  | 2.007 |
| 75 |  |  |  | 1.552 |  |  |  |  |  |  |  | 1.972 |
| 76 |  |  |  | 1.668 |  |  |  |  |  |  |  | 1.854 |
| 77 |  |  |  | 1.980 |  |  |  |  |  |  |  | 1.928 |
| 78 |  |  |  | 2.037 |  |  |  |  |  |  |  | 1.994 |
| 79 |  |  |  | 1.993 |  |  |  |  |  |  |  |  |
| 80 |  |  |  | 1.769 |  |  |  |  |  |  |  |  |
| 81 |  |  |  | 1.863 |  |  |  |  |  |  |  |  |
| 82 |  |  |  | 1.822 |  |  |  |  |  |  |  |  |
| 83 |  |  |  | 1.804 |  |  |  |  |  |  |  |  |

**Table S8.** DFA (Detrended Fluctuation Analysis) scaling exponent γ values for the 700 experimental cell trajectories.

| Cell number | Amoeba proteus |  |  |  | Metamoeba leningradensis |  |  |  | Amoeba borokensis |  |  |  |
| --- | --- | --- | --- | --- | --- | --- | --- | --- | --- | --- | --- | --- |
|  | Sc1 | Sc2 | Sc3 | Sc4 | Sc1 | Sc2 | Sc3 | Sc4 | Sc1 | Sc2 | Sc3 | Sc4 |
| 1 | 1.644 | 1.794 | 1.715 | 1.746 | 1.704 | 1.844 | 1.830 | 1.783 | 1.827 | 1.539 | 1.803 | 1.736 |
| 2 | 1.805 | 1.670 | 1.645 | 1.688 | 1.562 | 1.840 | 1.755 | 1.766 | 1.568 | 1.674 | 1.748 | 1.771 |
| 3 | 1.778 | 1.781 | 1.868 | 1.468 | 1.814 | 1.671 | 1.350 | 1.873 | 1.818 | 1.714 | 1.869 | 1.784 |
| 4 | 1.800 | 1.809 | 1.856 | 1.809 | 1.518 | 1.521 | 1.518 | 1.862 | 1.618 | 1.741 | 1.799 | 1.639 |
| 5 | 1.736 | 1.716 | 1.818 | 1.802 | 1.828 | 1.820 | 1.799 | 1.881 | 1.832 | 1.535 | 1.551 | 1.800 |
| 6 | 1.686 | 1.857 | 1.826 | 1.790 | 1.793 | 1.582 | 1.860 | 1.824 | 1.799 | 1.707 | 1.770 | 1.723 |
| 7 | 1.895 | 1.820 | 1.536 | 1.626 | 1.589 | 1.762 | 1.787 | 1.845 | 1.745 | 1.747 | 1.873 | 1.314 |
| 8 | 1.826 | 1.764 | 1.828 | 1.693 | 1.846 | 1.832 | 1.853 | 1.821 | 1.827 | 1.703 | 1.875 | 1.776 |
| 9 | 1.750 | 1.745 | 1.668 | 1.770 | 1.839 | 1.796 | 1.870 | 1.840 | 1.772 | 1.502 | 1.764 | 1.776 |
| 10 | 1.426 | 1.818 | 1.784 | 1.781 | 1.753 | 1.823 | 1.900 | 1.787 | 1.821 | 1.747 | 1.677 | 1.736 |
| 11 | 1.739 | 1.816 | 1.780 | 1.757 | 1.836 | 1.680 | 1.856 | 1.813 | 1.781 | 1.767 | 1.842 | 1.703 |
| 12 | 1.820 | 1.723 | 1.848 | 1.735 | 1.786 | 1.807 | 1.761 | 1.831 | 1.738 | 1.770 | 1.802 | 1.748 |
| 13 | 1.633 | 1.842 | 1.801 | 1.830 | 1.793 | 1.832 | 1.816 | 1.839 | 1.794 | 1.798 | 1.644 | 1.672 |
| 14 | 1.497 | 1.799 | 1.745 | 1.755 | 1.799 | 1.775 | 1.692 | 1.870 | 1.707 | 1.793 | 1.846 | 1.824 |
| 15 | 1.772 | 1.689 | 1.836 | 1.875 | 1.827 | 1.370 | 1.737 | 1.780 | 1.633 | 1.782 | 1.785 | 1.822 |
| 16 | 1.805 | 1.791 | 1.732 | 1.851 | 1.776 | 1.858 | 1.816 | 1.881 | 1.849 | 1.822 | 1.775 | 1.324 |
| 17 | 1.740 | 1.854 | 1.662 | 1.805 | 1.767 | 1.688 | 1.691 | 1.814 | 1.659 | 1.771 | 1.761 | 1.789 |
| 18 | 1.650 | 1.780 | 1.654 | 1.820 | 1.690 | 1.816 | 1.665 | 1.717 | 1.701 | 1.657 | 1.743 | 1.801 |
| 19 | 1.445 | 1.814 | 1.872 | 1.857 | 1.607 | 1.828 | 1.627 | 1.794 | 1.824 | 1.716 | 1.836 | 1.672 |
| 20 | 1.851 | 1.827 | 1.823 | 1.605 | 1.842 | 1.809 | 1.656 | 1.792 | 1.762 | 1.867 | 1.818 | 1.815 |
| 21 | 1.792 | 1.789 | 1.597 | 1.825 | 1.276 | 1.866 | 1.728 | 1.795 | 1.515 | 1.802 | 1.574 | 1.824 |
| 22 | 1.815 | 1.850 | 1.753 | 1.763 | 1.807 | 1.647 | 1.575 | 1.792 | 1.747 | 1.782 | 1.800 | 1.729 |
| 23 | 1.873 | 1.782 | 1.825 | 1.746 | 1.358 | 1.767 | 1.744 | 1.800 | 1.812 | 1.831 | 1.838 | 1.702 |
| 24 | 1.866 | 1.811 | 1.852 | 1.826 | 1.714 | 1.777 | 1.834 | 1.817 | 1.556 | 1.791 | 1.778 | 1.775 |
| 25 | 1.715 | 1.724 | 1.862 | 1.722 | 1.823 | 1.826 | 1.876 | 1.834 | 1.728 | 1.786 | 1.722 | 1.559 |
| 26 | 1.634 | 1.777 | 1.751 | 1.793 | 1.803 | 1.784 | 1.675 | 1.802 | 1.828 | 1.785 | 1.780 | 1.649 |
| 27 | 1.787 | 1.741 | 1.815 | 1.872 | 1.766 | 1.795 | 1.827 | 1.744 | 1.751 | 1.803 | 1.833 | 1.813 |
| 28 | 1.865 | 1.805 | 1.821 | 1.707 | 1.760 | 1.832 | 1.706 | 1.769 | 1.672 | 1.647 | 1.536 | 1.590 |
| 29 | 1.861 | 1.842 | 1.776 | 1.806 | 1.445 | 1.836 | 1.674 | 1.827 | 1.766 | 1.782 | 1.764 | 1.795 |
| 30 | 1.853 | 1.801 | 1.459 | 1.765 | 1.685 | 1.861 | 1.679 | 1.713 | 1.799 | 1.775 | 1.550 | 1.827 |
| 31 | 1.846 | 1.590 | 1.670 | 1.662 | 1.730 | 1.771 | 1.797 | 1.834 | 1.847 | 1.755 | 1.722 | 1.819 |
| 32 | 1.843 | 1.516 | 1.823 | 1.662 | 1.820 | 1.833 | 1.656 | 1.833 | 1.841 | 1.798 | 1.822 | 1.797 |
| 33 | 1.790 | 1.848 | 1.855 | 1.049 | 1.296 | 1.840 | 1.439 | 1.851 | 1.806 | 1.795 | 1.873 | 1.493 |
| 34 | 1.849 | 1.794 | 1.803 | 1.827 | 1.577 | 1.734 | 1.638 | 1.878 | 1.729 | 1.812 | 1.844 | 1.763 |
| 35 | 1.793 | 1.443 | 1.768 | 1.821 | 1.473 | 1.744 | 1.343 | 1.830 | 1.856 | 1.778 | 1.725 | 1.805 |
| 36 | 1.782 | 1.618 | 1.847 | 1.818 | 1.893 | 1.850 | 1.822 | 1.796 | 1.837 | 1.804 | 1.840 | 1.806 |
| 37 | 1.781 | 1.808 | 1.894 | 1.855 | 1.507 | 1.808 | 1.237 | 1.825 | 1.796 | 1.753 | 1.454 | 1.762 |
| 38 | 1.855 | 1.767 | 1.858 | 1.827 | 1.673 | 1.644 | 1.734 | 1.864 | 1.839 | 1.726 | 1.862 | 1.264 |
| 39 | 1.764 | 1.804 | 1.856 | 1.901 | 1.500 | 1.787 | 1.838 | 1.853 | 1.807 | 1.731 | 1.745 | 1.545 |
| 40 | 1.821 | 1.793 | 1.768 | 1.804 | 1.479 | 1.806 | 1.839 | 1.757 | 1.826 | 1.764 | 1.751 | 1.790 |
| 41 | 1.862 | 1.759 | 1.816 | 1.835 | 1.442 | 1.821 | 1.696 | 1.871 | 1.793 | 1.726 | 1.849 | 1.478 |
| 42 | 1.784 | 1.764 | 1.801 | 1.755 | 1.678 | 1.824 | 1.659 | 1.849 | 1.791 | 1.857 | 1.644 | 1.673 |
| 43 | 1.826 | 1.846 | 1.824 | 1.725 | 1.805 | 1.816 | 1.590 | 1.842 | 1.744 | 1.820 | 1.462 | 1.708 |
| 44 | 1.391 | 1.797 | 1.788 | 1.846 | 1.804 | 1.726 | 1.781 | 1.837 | 1.616 | 1.747 | 1.657 | 1.790 |
| 45 | 1.820 | 1.803 | 1.462 | 1.816 | 1.814 | 1.802 | 1.831 | 1.612 | 1.655 | 1.801 | 1.805 | 1.656 |
| 46 | 1.717 | 1.776 | 1.782 | 1.829 | 1.791 | 1.678 | 1.831 | 1.786 | 1.414 | 1.785 | 1.808 | 1.636 |
| 47 | 1.747 | 1.815 | 1.862 | 1.884 | 1.628 | 1.754 | 1.760 | 1.815 | 1.862 | 1.827 | 1.828 | 1.739 |
| 48 | 1.838 | 1.822 | 1.728 | 1.903 | 1.767 | 1.804 | 1.701 | 1.821 | 1.765 | 1.861 | 1.735 | 1.649 |
| 49 | 1.841 | 1.773 | 1.717 | 1.712 | 1.815 |  | 1.859 | 1.784 | 1.858 | 1.667 | 1.794 | 1.808 |
| 50 | 1.831 |  | 1.720 | 1.892 | 1.862 |  | 1.757 | 1.721 | 1.837 | 1.783 | 1.791 | 1.778 |
| 51 |  |  | 1.823 | 1.852 | 1.757 |  | 1.714 | 1.704 | 1.814 |  | 1.728 | 1.825 |
| 52 |  |  |  | 1.664 |  |  | 1.626 | 1.748 | 1.759 |  | 1.724 | 1.841 |
| 53 |  |  |  | 1.861 |  |  | 1.793 | 1.734 |  |  | 1.762 | 1.837 |
| 54 |  |  |  | 1.543 |  |  | 1.801 | 1.777 |  |  | 1.892 | 1.810 |
| 55 |  |  |  | 1.799 |  |  | 1.644 | 1.732 |  |  | 1.569 | 1.905 |
| 56 |  |  |  | 1.720 |  |  | 1.576 | 1.721 |  |  |  | 1.826 |
| 57 |  |  |  | 1.815 |  |  | 1.486 | 1.598 |  |  |  | 1.776 |
| 58 |  |  |  | 1.818 |  |  | 1.301 | 1.790 |  |  |  | 1.796 |
| 59 |  |  |  | 1.765 |  |  | 1.319 | 1.687 |  |  |  | 1.842 |
| 60 |  |  |  | 1.624 |  |  | 1.338 | 1.661 |  |  |  | 1.830 |
| 61 |  |  |  | 1.762 |  |  |  | 1.773 |  |  |  | 1.600 |
| 62 |  |  |  | 1.745 |  |  |  | 1.737 |  |  |  | 1.771 |
| 63 |  |  |  | 1.795 |  |  |  | 1.590 |  |  |  | 1.766 |
| 64 |  |  |  | 1.551 |  |  |  | 1.821 |  |  |  | 1.601 |
| 65 |  |  |  | 1.747 |  |  |  | 1.870 |  |  |  | 1.499 |
| 66 |  |  |  | 1.845 |  |  |  | 1.738 |  |  |  | 1.474 |
| 67 |  |  |  | 1.714 |  |  |  | 1.616 |  |  |  | 1.741 |
| 68 |  |  |  | 1.803 |  |  |  | 1.771 |  |  |  | 1.791 |
| 69 |  |  |  | 1.796 |  |  |  | 1.839 |  |  |  | 1.805 |
| 70 |  |  |  | 1.740 |  |  |  | 1.659 |  |  |  | 1.833 |
| 71 |  |  |  | 1.836 |  |  |  | 1.736 |  |  |  | 1.757 |
| 72 |  |  |  | 1.894 |  |  |  | 1.738 |  |  |  | 1.726 |
| 73 |  |  |  | 1.767 |  |  |  | 1.673 |  |  |  | 1.800 |
| 74 |  |  |  | 1.770 |  |  |  |  |  |  |  | 1.749 |
| 75 |  |  |  | 1.293 |  |  |  |  |  |  |  | 1.738 |
| 76 |  |  |  | 1.633 |  |  |  |  |  |  |  | 1.594 |
| 77 |  |  |  | 1.794 |  |  |  |  |  |  |  | 1.698 |
| 78 |  |  |  | 1.818 |  |  |  |  |  |  |  | 1.687 |
| 79 |  |  |  | 1.841 |  |  |  |  |  |  |  |  |
| 80 |  |  |  | 1.843 |  |  |  |  |  |  |  |  |
| 81 |  |  |  | 1.768 |  |  |  |  |  |  |  |  |
| 82 |  |  |  | 1.867 |  |  |  |  |  |  |  |  |
| 83 |  |  |  | 1.847 |  |  |  |  |  |  |  |  |

**Table S9.** DFA (Detrended Fluctuation Analysis) scaling exponent γ values for the 700 shuffled cell trajectories.

| Cell number | Amoeba proteus |  |  |  | Metamoeba leningradensis |  |  |  | Amoeba borokensis |  |  |  |
| --- | --- | --- | --- | --- | --- | --- | --- | --- | --- | --- | --- | --- |
|  | Sc1 | Sc2 | Sc3 | Sc4 | Sc1 | Sc2 | Sc3 | Sc4 | Sc1 | Sc2 | Sc3 | Sc4 |
| 1 | 0.582 | 0.412 | 0.344 | 0.615 | 0.325 | 0.660 | 0.531 | 0.350 | 0.410 | 0.474 | 0.310 | 0.659 |
| 2 | 0.570 | 0.670 | 0.577 | 0.505 | 0.477 | 0.624 | 0.507 | 0.426 | 0.312 | 0.389 | 0.519 | 0.351 |
| 3 | 0.318 | 0.470 | 0.472 | 0.482 | 0.291 | 0.329 | 0.506 | 0.507 | 0.500 | 0.349 | 0.563 | 0.443 |
| 4 | 0.496 | 0.346 | 0.421 | 0.411 | 0.529 | 0.510 | 0.694 | 0.293 | 0.512 | 0.393 | 0.351 | 0.410 |
| 5 | 0.577 | 0.579 | 0.568 | 0.350 | 0.280 | 0.603 | 0.499 | 0.610 | 0.435 | 0.456 | 0.497 | 0.536 |
| 6 | 0.435 | 0.435 | 0.483 | 0.383 | 0.462 | 0.323 | 0.454 | 0.448 | 0.352 | 0.373 | 0.627 | 0.372 |
| 7 | 0.493 | 0.411 | 0.472 | 0.464 | 0.699 | 0.591 | 0.357 | 0.539 | 0.509 | 0.467 | 0.424 | 0.509 |
| 8 | 0.582 | 0.467 | 0.570 | 0.592 | 0.566 | 0.563 | 0.421 | 0.500 | 0.393 | 0.633 | 0.621 | 0.417 |
| 9 | 0.725 | 0.491 | 0.249 | 0.330 | 0.402 | 0.496 | 0.400 | 0.508 | 0.628 | 0.519 | 0.620 | 0.520 |
| 10 | 0.515 | 0.637 | 0.615 | 0.615 | 0.322 | 0.413 | 0.560 | 0.439 | 0.421 | 0.545 | 0.397 | 0.347 |
| 11 | 0.320 | 0.637 | 0.618 | 0.493 | 0.476 | 0.501 | 0.491 | 0.514 | 0.380 | 0.417 | 0.376 | 0.555 |
| 12 | 0.492 | 0.631 | 0.420 | 0.331 | 0.510 | 0.640 | 0.585 | 0.275 | 0.580 | 0.450 | 0.314 | 0.584 |
| 13 | 0.479 | 0.326 | 0.488 | 0.505 | 0.397 | 0.628 | 0.274 | 0.667 | 0.520 | 0.554 | 0.522 | 0.680 |
| 14 | 0.591 | 0.435 | 0.542 | 0.382 | 0.443 | 0.314 | 0.396 | 0.431 | 0.434 | 0.447 | 0.390 | 0.561 |
| 15 | 0.480 | 0.353 | 0.456 | 0.546 | 0.444 | 0.433 | 0.401 | 0.358 | 0.607 | 0.461 | 0.337 | 0.460 |
| 16 | 0.559 | 0.500 | 0.517 | 0.460 | 0.330 | 0.322 | 0.439 | 0.375 | 0.567 | 0.538 | 0.493 | 0.432 |
| 17 | 0.498 | 0.523 | 0.641 | 0.525 | 0.315 | 0.590 | 0.288 | 0.566 | 0.460 | 0.462 | 0.419 | 0.583 |
| 18 | 0.618 | 0.530 | 0.499 | 0.450 | 0.362 | 0.578 | 0.500 | 0.560 | 0.515 | 0.616 | 0.335 | 0.446 |
| 19 | 0.480 | 0.476 | 0.375 | 0.434 | 0.492 | 0.339 | 0.523 | 0.654 | 0.458 | 0.453 | 0.473 | 0.437 |
| 20 | 0.565 | 0.440 | 0.504 | 0.563 | 0.369 | 0.616 | 0.385 | 0.330 | 0.411 | 0.372 | 0.544 | 0.517 |
| 21 | 0.573 | 0.505 | 0.409 | 0.564 | 0.470 | 0.567 | 0.401 | 0.395 | 0.701 | 0.290 | 0.435 | 0.599 |
| 22 | 0.361 | 0.581 | 0.346 | 0.397 | 0.507 | 0.384 | 0.538 | 0.407 | 0.566 | 0.375 | 0.496 | 0.492 |
| 23 | 0.448 | 0.507 | 0.451 | 0.381 | 0.430 | 0.346 | 0.572 | 0.534 | 0.471 | 0.541 | 0.450 | 0.558 |
| 24 | 0.605 | 0.530 | 0.403 | 0.526 | 0.631 | 0.368 | 0.278 | 0.584 | 0.388 | 0.485 | 0.444 | 0.552 |
| 25 | 0.348 | 0.491 | 0.545 | 0.344 | 0.302 | 0.405 | 0.597 | 0.602 | 0.536 | 0.531 | 0.587 | 0.443 |
| 26 | 0.427 | 0.401 | 0.455 | 0.481 | 0.368 | 0.474 | 0.567 | 0.464 | 0.563 | 0.429 | 0.472 | 0.530 |
| 27 | 0.644 | 0.328 | 0.413 | 0.409 | 0.390 | 0.437 | 0.445 | 0.507 | 0.456 | 0.249 | 0.689 | 0.542 |
| 28 | 0.495 | 0.504 | 0.412 | 0.566 | 0.448 | 0.509 | 0.560 | 0.452 | 0.557 | 0.284 | 0.548 | 0.403 |
| 29 | 0.494 | 0.526 | 0.414 | 0.657 | 0.466 | 0.645 | 0.420 | 0.417 | 0.452 | 0.560 | 0.449 | 0.447 |
| 30 | 0.564 | 0.293 | 0.514 | 0.446 | 0.445 | 0.481 | 0.348 | 0.446 | 0.313 | 0.380 | 0.508 | 0.645 |
| 31 | 0.430 | 0.465 | 0.473 | 0.399 | 0.500 | 0.464 | 0.410 | 0.400 | 0.395 | 0.361 | 0.301 | 0.411 |
| 32 | 0.506 | 0.352 | 0.442 | 0.257 | 0.477 | 0.539 | 0.523 | 0.345 | 0.544 | 0.649 | 0.449 | 0.345 |
| 33 | 0.420 | 0.530 | 0.329 | 0.534 | 0.506 | 0.635 | 0.802 | 0.522 | 0.578 | 0.386 | 0.452 | 0.540 |
| 34 | 0.557 | 0.509 | 0.569 | 0.410 | 0.481 | 0.714 | 0.510 | 0.565 | 0.431 | 0.449 | 0.400 | 0.560 |
| 35 | 0.454 | 0.572 | 0.415 | 0.417 | 0.545 | 0.446 | 0.389 | 0.351 | 0.486 | 0.587 | 0.466 | 0.619 |
| 36 | 0.451 | 0.542 | 0.471 | 0.417 | 0.547 | 0.410 | 0.368 | 0.454 | 0.389 | 0.757 | 0.421 | 0.393 |
| 37 | 0.584 | 0.623 | 0.370 | 0.476 | 0.470 | 0.401 | 0.487 | 0.413 | 0.382 | 0.603 | 0.401 | 0.494 |
| 38 | 0.365 | 0.495 | 0.549 | 0.411 | 0.561 | 0.486 | 0.393 | 0.488 | 0.348 | 0.495 | 0.431 | 0.371 |
| 39 | 0.503 | 0.435 | 0.591 | 0.513 | 0.710 | 0.570 | 0.535 | 0.373 | 0.500 | 0.619 | 0.529 | 0.514 |
| 40 | 0.514 | 0.561 | 0.505 | 0.445 | 0.454 | 0.339 | 0.404 | 0.320 | 0.337 | 0.500 | 0.385 | 0.490 |
| 41 | 0.702 | 0.413 | 0.502 | 0.441 | 0.507 | 0.464 | 0.646 | 0.438 | 0.453 | 0.400 | 0.530 | 0.669 |
| 42 | 0.456 | 0.405 | 0.366 | 0.579 | 0.534 | 0.646 | 0.589 | 0.626 | 0.525 | 0.654 | 0.471 | 0.416 |
| 43 | 0.459 | 0.509 | 0.531 | 0.556 | 0.249 | 0.461 | 0.511 | 0.472 | 0.375 | 0.433 | 0.571 | 0.484 |
| 44 | 0.483 | 0.516 | 0.247 | 0.391 | 0.446 | 0.483 | 0.494 | 0.465 | 0.393 | 0.378 | 0.549 | 0.394 |
| 45 | 0.542 | 0.411 | 0.480 | 0.762 | 0.442 | 0.337 | 0.516 | 0.350 | 0.306 | 0.507 | 0.655 | 0.415 |
| 46 | 0.488 | 0.460 | 0.489 | 0.321 | 0.595 | 0.563 | 0.419 | 0.521 | 0.447 | 0.486 | 0.273 | 0.412 |
| 47 | 0.642 | 0.461 | 0.458 | 0.316 | 0.471 | 0.571 | 0.555 | 0.592 | 0.396 | 0.511 | 0.378 | 0.571 |
| 48 | 0.525 | 0.425 | 0.495 | 0.521 | 0.311 | 0.517 | 0.516 | 0.577 | 0.537 | 0.473 | 0.660 | 0.476 |
| 49 | 0.525 | 0.352 | 0.636 | 0.463 | 0.467 |  | 0.508 | 0.640 | 0.597 | 0.349 | 0.424 | 0.633 |
| 50 | 0.462 |  | 0.667 | 0.441 | 0.462 |  | 0.539 | 0.706 | 0.515 | 0.526 | 0.418 | 0.431 |
| 51 |  |  | 0.326 | 0.488 | 0.427 |  | 0.488 | 0.478 | 0.472 |  | 0.341 | 0.447 |
| 52 |  |  |  | 0.481 |  |  | 0.407 | 0.509 | 0.401 |  | 0.479 | 0.373 |
| 53 |  |  |  | 0.480 |  |  | 0.586 | 0.646 |  |  | 0.677 | 0.445 |
| 54 |  |  |  | 0.377 |  |  | 0.527 | 0.243 |  |  | 0.374 | 0.533 |
| 55 |  |  |  | 0.308 |  |  | 0.626 | 0.545 |  |  | 0.403 | 0.366 |
| 56 |  |  |  | 0.355 |  |  | 0.509 | 0.408 |  |  |  | 0.361 |
| 57 |  |  |  | 0.491 |  |  | 0.396 | 0.458 |  |  |  | 0.633 |
| 58 |  |  |  | 0.556 |  |  | 0.570 | 0.603 |  |  |  | 0.460 |
| 59 |  |  |  | 0.695 |  |  | 0.469 | 0.445 |  |  |  | 0.369 |
| 60 |  |  |  | 0.560 |  |  | 0.351 | 0.333 |  |  |  | 0.493 |
| 61 |  |  |  | 0.426 |  |  |  | 0.562 |  |  |  | 0.351 |
| 62 |  |  |  | 0.585 |  |  |  | 0.464 |  |  |  | 0.628 |
| 63 |  |  |  | 0.475 |  |  |  | 0.562 |  |  |  | 0.604 |
| 64 |  |  |  | 0.603 |  |  |  | 0.564 |  |  |  | 0.544 |
| 65 |  |  |  | 0.449 |  |  |  | 0.361 |  |  |  | 0.537 |
| 66 |  |  |  | 0.349 |  |  |  | 0.527 |  |  |  | 0.417 |
| 67 |  |  |  | 0.638 |  |  |  | 0.512 |  |  |  | 0.586 |
| 68 |  |  |  | 0.578 |  |  |  | 0.474 |  |  |  | 0.384 |
| 69 |  |  |  | 0.478 |  |  |  | 0.394 |  |  |  | 0.565 |
| 70 |  |  |  | 0.469 |  |  |  | 0.369 |  |  |  | 0.325 |
| 71 |  |  |  | 0.331 |  |  |  | 0.480 |  |  |  | 0.568 |
| 72 |  |  |  | 0.470 |  |  |  | 0.551 |  |  |  | 0.320 |
| 73 |  |  |  | 0.510 |  |  |  | 0.559 |  |  |  | 0.628 |
| 74 |  |  |  | 0.510 |  |  |  |  |  |  |  | 0.472 |
| 75 |  |  |  | 0.659 |  |  |  |  |  |  |  | 0.632 |
| 76 |  |  |  | 0.599 |  |  |  |  |  |  |  | 0.414 |
| 77 |  |  |  | 0.323 |  |  |  |  |  |  |  | 0.426 |
| 78 |  |  |  | 0.503 |  |  |  |  |  |  |  | 0.492 |
| 79 |  |  |  | 0.442 |  |  |  |  |  |  |  |  |
| 80 |  |  |  | 0.549 |  |  |  |  |  |  |  |  |
| 81 |  |  |  | 0.431 |  |  |  |  |  |  |  |  |
| 82 |  |  |  | 0.511 |  |  |  |  |  |  |  |  |
| 83 |  |  |  | 0.550 |  |  |  |  |  |  |  |  |

**Table S10.** Kruskal-Wallis and Wilcoxon analyses to assess the variability in kinetic properties.

| Species | Scenario | Intensity of Response | Directionality Ratio | Average Speed |
| --- | --- | --- | --- | --- |
| Amoeba Proteus | Sc1-Sc2 | 0.006 | $10^{-8}$ | 0.002 |
| | Sc1-Sc3 | 0.959 | 0.080 | $10^{-4}$ |
| | Sc1-Sc4 | 0.155 | 0.084 | $10^{-12}$ |
| | Sc2-Sc3 | $10^{-4}$ | $10^{-7}$ | 0.544 |
| | Sc2-Sc4 | $10^{-8}$ | $10^{-7}$ | 0.001 |
|  | Sc3-Sc4 | 0.100 | 0.865 | 0.005 |
| | All | $10^{-6}$ | $10^{-9}$ | $10^{-10}$ |
| Metamoeba leningradensis | Sc1-Sc2 | $10^{-8}$ | $10^{-8}$ | $10^{-4}$ |
|  | Sc1-Sc3 | 0.098 | 0.734 | 0.004 |
| | Sc1-Sc4 | $10^{-13}$ | $10^{-6}$ | $10^{-13}$ |
| | Sc2-Sc3 | $10^{-5}$ | $10^{-9}$ | 0.899 |
| | Sc2-Sc4 | 0.010 | 0.050 | $10^{-8}$ |
| | Sc3-Sc4 | $10^{-11}$ | $10^{-7}$ | $10^{-8}$ |
| | All | $10^{-16}$ | $10^{-12}$ | $10^{-15}$ |
| Amoeba borokensis | Sc1-Sc2 | $10^{-4}$ | $10^{-5}$ | 0.160 |
|  | Sc1-Sc3 | 0.884 | 0.918 | 0.326 |
| | Sc1-Sc4 | 0.099 | 0.862 | $10^{-4}$ |
| | Sc2-Sc3 | 0.003 | $10^{-4}$ | 0.982 |
| | Sc2-Sc4 | 0.029 | $10^{-4}$ | 0.119 |
|  | Sc3-Sc4 | 0.133 | 0.659 | 0.230 |
| | All | 0.002 | $10^{-5}$ | 0.022 |
| Amoeba proteus Vs. Metamoeba leningradensis | Sc1-Sc2 | 0.263 | $10^{-7}$ | $10^{-8}$ |
| | Sc1-Sc3 | 0.006 | 0.318 | $10^{-7}$ |
| | Sc1-Sc4 | $10^{-4}$ | $10^{-5}$ | 0.417 |
| | Sc2-Sc3 | $10^{-8}$ | $10^{-10}$ | 0.010 |
| | Sc2-Sc4 | 0.573 | 0.002 | $10^{-4}$ |
| | Sc3-Sc4 | $10^{-4}$ | 0.003 | $10^{-5}$ |
|  | All | 0.293 | 0.793 | 0.065 |
| Amoeba proteus Vs. Amoeba borokensis | Sc1-Sc2 | 0.537 | $10^{-5}$ | $10^{-11}$ |
| | Sc1-Sc3 | 0.002 | 0.895 | $10^{-8}$ |
| | Sc1-Sc4 | 0.009 | 0.559 | $10^{-12}$ |
| | Sc2-Sc3 | $10^{-8}$ | $10^{-7}$ | $10^{-4}$ |
| | Sc2-Sc4 | $10^{-9}$ | $10^{-7}$ | $10^{-5}$ |
| | Sc3-Sc4 | 0.004 | 0.106 | $10^{-4}$ |
| | All | $10^{-10}$ | 0.032 | $10^{-17}$ |
| Metamoeba leningradensis Vs. Amoeba borokensis | Sc1-Sc2 | $10^{-5}$ | $10^{-6}$ | 0.064 |
|  | Sc1-Sc3 | 0.383 | 0.515 | 0.131 |
|  | Sc1-Sc4 | 0.005 | 0.228 | 0.001 |
| | Sc2-Sc3 | $10^{-6}$ | $10^{-7}$ | 0.085 |
| | Sc2-Sc4 | $10^{-5}$ | $10^{-7}$ | 0.375 |
|  | Sc3-Sc4 | 0.529 | 0.117 | 0.401 |
| | All | $10^{-5}$ | 0.067 | $10^{-7}$ |

